# Herbarium Specimen Sequencing Allows Precise Datation of *Xanthomonas citri* pv. *citri* Diversification History

**DOI:** 10.1101/2022.12.08.519547

**Authors:** PE Campos, O Pruvost, K Boyer, F Chiroleu, TT Cao, M Gaudeul, C Baider, TMA Utteridge, S Dominick, N Becker, A Rieux, L Gagnevin

## Abstract

Over the past decade, the field of ancient genomics has triggered considerable progress in the study of various pathogens, including those affecting crops. In this context, herbarium collections have been an important source of dated, identified and preserved DNA, whose use in comparative genomics and phylogeography may shed light into the emergence and evolutionary history of plant pathogens. In this study, we reconstructed 13 historical genomes of the bacterial crop pathogen *Xanthomonas citri* pv. *citri* (*Xci*) from infected citrus herbarium specimens using a shotgun-based deep sequencing strategy. Following authentication of the historical genomes based on ancient DNA damage patterns, we compared them to a large set of modern genomes to reconstruct their phylogenetic relationships, pathogeny-associated genes content and estimate several evolutionary parameters, using Bayesian tip-dating calibration and phylogeography inferences. Our results reveal that *Xci* originated in Southern Asia ~11,500 years ago and diversified during the beginning of the 13^th^ century, after *Citrus* diversification and before spreading to the rest of the world. This updated scenario links *Xci* specialization to Neolithic climatic change and the development of agriculture, and its diversification to the human-driven expansion of citriculture through the early East-West trade and later colonization. The analysis of data obtained from such historical specimens is challenging and must undergo adapted treatment before being compared to modern samples. Nevertheless, we confirm here that herbarium collections are a precious tool to improve the knowledge of the evolutionary history of plant pathogens.

## Introduction

Plant pathogens have plagued human societies since the beginning of agriculture (Dark *et al.* 2001). In such human-engineered ecosystems, the high density and low genetic diversity of hosts, as well as the environmental homogeneity caused by agricultural practices, facilitated the rise and propagation of diseases, with the evolution of host-adapted and virulent pathogens (Mira *et al.* 2006, Stukenbrock *et al.* 2008). Intensification of agriculture, monocultures size rise and trade globalization contributed to the emergence and expansion of pathogens, with opportunities to meet new naive host populations and realize host shift or host jump (Anderson *et al.* 2004, McCann 2020, Stukenbrock *et al.* 2008).

Today, plant pathogens and pests cause up to 40% yield loss in major crops, threatening food security (Savary *et al.* 2019), agrobiodiversity conservation and public health (Anderson *et al.* 2004, Bernardes *et al.* 2015). A better understanding of the factors underlying the origin, evolution and emergence of pathogens would help assess the risks they pose to crops and improve tools for surveillance and disease control. The combination of genetic material obtained from historic biological collections such as herbaria and modern samples provides heterochronous datasets which can improve phylogenetic estimates of evolutionary parameters and the timelines of their emergence and spread by bringing robust time component to inferences (Duchêne *et al.* 2020, Malmstrom *et al.* 2022, Rieux *et al.* 2014). Indeed, adding ancient or historical sequences expands the temporal range of the dataset, increasing the chance to detect evolutionary change, *i.e.,* temporal signal, which can be used to infer substitution rates and divergence time between lineages, as well as sudden modifications in genetic diversity (Drummond *et al.* 2002, Drummond *et al.* 2003, Rieux *et al.* 2016a).

The most well-studied crop pathosystem using historical herbarium genetic material is *Phytophthora infestans,* the oomycete responsible for potato late blight. Through the sequencing of 19^th^ century infected specimens, the strain which caused the great potato famine in 1845-1849 has been identified and its genome characterized. Phylogeny reconstruction showed the historical strain to have originated from a secondary diversification area of the pathogen in North America from where one or a few dispersal events caused *P. infestans* emergence in Europe (Martin *et al.* 2013, Ristaino 2020, Saville *et al.* 2016, Yoshida *et al.* 2014, Yoshida *et al.* 2015, Yoshida *et al.* 2013). Similar studies reconstructing the evolutionary history of crop pathogens from full genomes have been successfully realized on viruses as well (Al Rwahnih *et al.* 2015, Malmstrom *et al.* 2007, Rieux *et al.* 2021, Smith *et al.* 2014). We recently described the history of the local emergence of the bacterial crop pathogen *Xanthomonas citri* pv. *citri (Xci)* in the South West Indian Ocean, using the first ancient bacterial genome retrieved from a herbarium specimen (Campos *et al.* 2021).

*Xci,* responsible for Asiatic Citrus Canker (ACC) and found in most subtropical citrus-producing regions, is a serious threat to citriculture. With no available definitive control measure, the disease causes important economic losses, both by decreasing fruit yield and quality, and because of *Xci* quarantine organism status (Gottwald *et al.* 2002, Talon *et al.* 2020). *Xci* comprises three major pathotypes, discriminated by genetic diversity and host range. Pathotype A, with the broadest host-range (nearly all *Citrus* and several related rutaceous genera), is the most prevalent worldwide (Graham *et al.* 2004). Pathotypes A* and A^W^, primarily reported from Asia, are restricted to *Citrus aurantiifolia* and its close relative *Citrus macrophylla* (Schubert *et al.* 2001, Vernière *et al.* 1998). Pathotype A* also occasionally infects Tahiti lime (*C. latifolia*) or sweet lime (*C. limettioides*). A specificity of pathotype A^W^ is to elicit a hypersensitive response on several *Citrus* species, including *C. paradisi* and *C. sinensis* (Sun *et al.* 2004). The phylogenetic relationships between the different pathotypes, first reconstructed using minisatellite molecular markers (Pruvost *et al.* 2014), before obtaining more resolutive data from whole genomes (Gordon *et al.* 2015, Zhang *et al.* 2015), suggested that pathotypes A and A^W^ are more closely related to each other than they are to pathotype A*. Recently, comparative genomic and phylogenomic analysis of 95 contemporary genomes were used to identify pathotype-specific virulence-associated genes and infer a probable scenario for *Xci* origin and diversification (Patané *et al.* 2019). This study revealed that the origin of *Xci* occurred much more recently than the main phylogenetic splits of *Citrus* plants, suggesting dispersion, rather than host-directed vicariance, as the main driver of this pathogen geographic expansion. However, the sole use of modern genomes impeded the detection of sufficient *de novo* evolutionary change within the dataset (as referring to “measurably evolving populations” (Biek *et al.* 2015, Drummond *et al.* 2003)). The authors were thus compelled to build a timeframe of evolution based on both the extrapolation of rates from external measures (*i.e.* rate dating) and a constraint on the distribution of a single external node age (*i.e.* node dating), two dating methodologies known to yield potential misleading estimates (Ho *et al.* 2008, Rieux *et al.* 2014).

In the present study, we took advantage of an extensive sampling covering the last 70 years of *Xci* strains evolution, along with a broad representation of *Citrus* specimens in herbaria dating back to the 19^th^ century. We sequenced historical bacterial genomes of *Xci* from 13 herbarium samples showing typical canker symptoms, and originating from the putative center of origin of the pathogen. Authentication based on DNA degradation patterns was established, allowing us to identify, in more detail, a significant contribution of sample age and library production protocol to deamination rate. We then compared the historical genomes to those of 171 modern strains representative of the worldwide genetic diversity, 57 of which specifically sequenced for the purpose of this study. We aimed to improve knowledge about *Xci* origin and diversification history by *1*) reconstructing a thorough time-calibrated phylogeny and inferring evolutionary parameters based on a robust dating approach within the measurably evolving populations framework, *2*) inferring the ancestral geographical state of lineages and estimating source populations of epidemics, *3*) assessing the pathogenicity-associated gene content across all lineages.

## Material & Methods

### Herbarium material sampling

The collections of the Royal Botanic Gardens, Kew (K) (https://www.kew.org/science/collections-and-resources/collections/herbarium), the Mauritius Herbarium (MAU) (https://agriculture.govmu.org/Pages/Departments/Departments/The-Mauritius-Herbarium.aspx), the Muséum national d’Histoire naturelle (P) (https://www.mnhn.fr/fr/collections/ensembles-collections/botanique), the US National Fungus Collections (BPI) (https://nt.ars-grin.gov/fungaldatabases/specimens/specimens.cfm) and the U.S. National Herbarium (US) (https://collections.nmnh.si.edu/search/botany/) were prospected between May 2016 and October 2017. *Citrus* specimens displaying typical Asiatic citrus canker lesions were sampled on site using sterile equipment and transported back to the laboratory inside individual envelopes where they were stored at 17°C in vacuum-sealed boxes until use. Thirteen historic specimens collected between 1845 to 1974 and conserved in five different herbaria were selected for analysis (Table 1). Those were chosen as the oldest available from Asia, the supposed geographic origin of *Xci,* as well as from Oceania and the Southwest Indian Ocean. Among the 13 herbarium specimens, 12 were processed during the course of this study, while HERB_1937 was processed previously (Campos *et al.* 2021).

**Table 1.** General characteristics of the 13 herbarium specimens.

| ID | Herbarium code | Herbarium specimen ID | Collection year | Location | Host |
| --- | --- | --- | --- | --- | --- |
| HERB_1845 | P | P05297986 | 1845 | Indonesia, Java | *Citrus aurantiifolia* |
| HERB_1852 | K | Q1874 | 1852 | India, Khasi hills | *Citrus medica* |
| HERB_1854 | K | Q1954 | 1854 | Indonesia, Java | *Citrus aurantiifolia* |
| HERB_1859 | P | P05240716 | 1859 | Bangladesh | *Citrus medica* |
| HERB_1865 | K | Q1889 | 1865 | India | *Citrus medica* |
| HERB_1884 | K | 1206 | 1884 | Philippines, Luzon | *Citrus medica* |
| HERB_1911 | P | P05297996 | 1911 | Indonesia, Java | *Citrus aurantiifolia* |
| HERB_1915 | P | P05297992 | 1915 | Philippines | *Citrus* sp. |
| HERB_1922 | US | 1756364 | 1922 | China, Yunnan | *Citrus medica* |
| HERB_1937 | MAU | MAU0015151 | 1937 | Mauritius | *Citrus* sp. |
| HERB_1946 | BPI | 686249 | 1946 | Guam, Tlofofo | *Citrus* sp. |
| HERB_1963 | K | 630116 | 1963 | Nepal, Sanichare | *Citrus medica* |
| HERB_1974 | MAU | MAU0015154 | 1974 | Mauritius | *Citrus* sp. |

### Ancient DNA extraction and library preparation

DNA extraction from herbarium samples was performed in a bleach-cleaned facility room with no exposure to modern *Xci* DNA, as described in Campos *et al.* (2021), along with herbarium samples of *Coffea* sp., a non *Xci*-host species acting as negative control. Following extraction, fragment size and concentration were controlled using TapeStation (Agilent Technologies) high sensitivity assays, according to the manufacturers’ recommendations. Seven herbarium samples were converted into double-stranded libraries using the aDNA-adapted BEST (Blunt-End-Single-Tube) protocol from Carøe *et al.* (2017). Library preparation of the remaining six herbarium samples was outsourced to Fasteris (https://www.fasteris.com/dna/) where DNA was converted into a double-stranded library following a custom TruSeq Nano DNA protocol (Illumina) without prior fragmentation and using a modified bead ratio adapted to small fragments.

### Modern bacterial strains culture, DNA extraction and library preparation

Fifty-seven bacterial strains isolated between 1963 and 2008, mainly from Asia (S1 Table) and stored as lyophiles at −80°C, were chosen to complete the collection of available modern genomes. Strains were grown at 28°C on YPGA (7 g/L yeast extract, 7 g/L peptone, 7 g/L glucose, 18 g/L agar, supplemented by 20 mg/L propiconazole, pH 7.2). Single cultures were used for DNA extraction using the Wizard® genomic DNA purification kit (Promega) following the manufacturer’s instructions. Quality assessment was realized for concentration using QuBit (Invitrogen) and Nanodrop (Thermo Fisher Scientific) fluorometers. Library preparation of the modern strains was outsourced to Fasteris where classic TruSeq Nano DNA protocol following Nextera enzymatic DNA fragmentation was applied (Illumina). Sequencing for both historical and modern DNA was performed in a paired-end 2×150 cycles configuration on a NextSeq500 machine in several batches, with samples from both types of libraries being independently treated.

### Initial reads trimming and merging

Artefactual homopolymer sequences were removed from libraries when presenting entropy inferior than 0.6 using BBDuk from BBMap 37.92 (DOE Joint Genome Institute). Adaptors were trimmed using the Illuminaclip option from Trimmomatic 0.36 (Bolger *et al.* 2014). Such reads were processed into the *post-mortem* DNA damage assessment pipeline detailed in the section below. Additional quality-trimming was realized with Trimmomatic based on base-quality (LEADING:15; TRAILING:15; SLIDINGWINDOW:5:15) and read length (MINLEN:30). Paired reads were then merged using AdapterRemoval 2.2.2 (Schubert *et al.* 2016) with default options.

### Ancient DNA damage assessment and statistical analyses

*Post-mortem* DNA damage was measured by DNA fragment length distribution and terminal deamination patterns using mapDamage 2.2.1 (Jonsson *et al.* 2013). Alignments required for mapDamage were performed with an aligner adapted to short reads BWA-aln 0.7.15 (default options, seed disabled) (Li *et al.* 2009) for the herbarium samples and Bowtie 2 (options --non-deterministic --very-sensitive) (Langmead *et al.* 2012) for modern strains, using *Xci* reference strain IAPAR 306 genome (chromosome NC_003919.1, plasmids pXAC33 NC_003921.3 and pXAC64 NC_003922.1). MarkDuplicates in picardtools 2.7.0 (Broad Institute) was run to remove PCR duplicates. For each sample, reads were grouped in nucleotide length classes of 25 nucleotide-long intervals, from 15 to 290 nucleotides (*i.e.,* 11 classes). Analysis of variance aov function (“stats” R package) was used to test the effect of protocol (BEST or TruSeq), DNA type (plasmid or chromosome), and age (years) on nucleotide length classes. When significant, a *Student t.test* was performed. Deamination rate data of each sample took into account: the number of terminal 3’ G>A substitutions, divided by the total number of reads bearing a G at the same position (according to the reference sequence). Effect of protocol (BEST or TrueSeqNano) and age (in years) on deamination rate was assessed using a glm model (“stats” R package), under quasi-binomial distribution. Effect of DNA type (plasmid or chromosome) was specifically tested using glmmPQL function, allowing the analysis of paired variables.

### Genome reconstruction

Genomes were reconstructed by mapping quality-trimmed reads to *Xci* reference strain IAPAR 306 genome using BWA-aln for short reads (Li *et al.* 2009) and Bowtie 2 for longer reads (Langmead *et al.* 2012), as defined above. Sequencing depths were computed using BEDTools genomecov 2.24.0 (Quinlan *et al.* 2010). For herbarium specimens, BAM (Binary Alignment Map) files were extremity-trimmed on their 5 external nucleotides at each end using BamUtil 1.0.14 (Jun *et al.* 2015). SNPs (Single Nucleotide Polymorphisms) were called with GATK UnifiedGenotyper (DePristo *et al.* 2011); they were considered dubious and filtered out if they met at least one of the following conditions: “depth<20”, “minor allelic frequency<0.9” and “mapping quality<30”. Consensus sequences were then reconstructed by introducing the high-quality SNPs in the *Xci* reference genome and replacing dubious SNPs and non-covered sites (depth=0) by an N.

### Phylogeny & tree-calibration

A dataset of 171 modern *Xci* genomes (date range: 1948 - 2017) representative of *Xci* global diversity was built from 114 previously published genomes and 57 new genomes generated within the course of this study (S1 Table). An alignment of the 13 historical chromosome sequences with the 171 modern sequences was constructed for phylogenetic analyses, with strains of *Xanthomonas axonopodis* pv. *vasculorum* NCPPB-796 from Mauritius (isolated in 1960, GCF_013177355.1), *Xanthomonas citri* pv. *cajani* LMG558 from India (1950, GCF_002019105.1) and *Xanthomonas citri* pv. *clitoriae* LMG9045 from India (1974, GCA_002019345.1) as outgroups. Variants from modern strains were independently called and filtered using the same parameters as for historical genomes. Recombinant regions were identified inside the *Xci* dataset (ingroup only) with ClonalFrameML (Didelot *et al.* 2015) and removed, to avoid production of incongruent trees during phylogenetic reconstruction. Two SNP (Single Nucleotide Polymorphism) datasets were constructed, either within the *Xci* ingroup only, or from across the whole dataset (ingroup + outgroups). A Maximum Likelihood (ML) tree was constructed on both SNP alignments using RAxML 8.2.4 (Stamatakis 2014) using a rapid Bootstrap analysis, a General Time-Reversible model of evolution following a Γ distribution with four rate categories (GTRGAMMA) and 1,000 alternative runs (Lanave *et al.* 1984).

As a requirement to build tip-calibrated phylogenies, the existence of a temporal signal was investigated thanks to three different tests. First, a linear regression test between sample age and root-to-tip distances was computed at each internal node of the ML tree using Phylostems (Doizy *et al.* 2020). Temporal signal was considered present at nodes displaying a significant positive correlation. Secondly, a date-randomization test (DRT) (Duchêne *et al.* 2015b) was performed with 20 independent date-randomized datasets generated using the R package “TipDatingBeast” (Rieux *et al.* 2016b). Temporal signal was considered present when there was no overlap between the inferred root height 95% Highest Posterior Density (95% HPD) of the initial dataset and that of 20 date-randomized datasets. Finally, a Mantel test with 1,000 date-randomized iterations investigating whether closely related sequences were more likely to have been sampled at similar times was also performed to ensure no confounding effect between temporal and genetic structure, as recent work suggested that temporal signal investigation through root to-tip-regression and DRT could be misled in such a case (Murray *et al.* 2016).

Tip-dating calibration Bayesian inferences (BI) were performed on the primary SNP alignment (*Xci* ingroup) with BEAST 1.8.4 (Drummond *et al.* 2007). Leaf heights were constrained to be proportional to sample ages. Flat priors (*i.e.,* uniform distributions) for the substitution rate (10^-12^ to 10^-2^ substitutions/site/year) and for the age of all internal nodes in the tree were applied. We also considered a GTR substitution model with a Γ distribution and invariant sites (GTR+G+I), an uncorrelated relaxed log-normal clock to account for variations between lineages, and a tree prior for demography of coalescent extended Bayesian skyline. The Bayesian topology was conjointly estimated with all other parameters during the Markov chain Monte-Carlo (MCMC) and no prior information from the tree was incorporated in BEAST. Five independent chains were run for 200 million steps and sampled every 20,000 steps, discarding the first 20,000 steps as burn-in. BEAGLE (Broad-platform Evolutionary Analysis General Likelihood Evaluator) library was used to improve computational speed (Ayres *et al.* 2012, Suchard *et al.* 2009). Convergence to the stationary, sufficient sampling (effective sample size > 200) and mixing were checked by inspecting posterior samples with Tracer 1.7.1 (Rambaut *et al.* 2018). Final parameters estimation was based on the combination of the different chains. Maximum clade credibility method in TreeAnnotator (Drummond *et al.* 2007) was used to determine the best-supported tree of the combined chains.

Rate-dating calibration outside the *Xci* clade was performed on the secondary SNP alignment (ingroup + outgroups) with BEAST 1.8.4 (Drummond *et al.* 2007). Instead of using tip-dates, we applied a prior on the substitution rate by drawing values from a normal distribution with mean and standard deviation values fixed as those inferred using tip-dating calibration within *Xci* ingroup. All other parameters were applied as described previously.

### Phylogeography & ancestral location state reconstruction

The presence of geographic structure in the ML tree was measured through calculation of the Association Index (AI) and the comparison of its value with the ones computed from 1,000 location-randomized trees (Parker *et al.* 2008). Non-random association between phylogeny and location was assumed when less than 5% of AI values computed from the randomized trees were smaller than the AI value of the real ML tree.

Ancestral location state was reconstructed using BEAST 1.8.4 under the same parameters as tip-dating calibration but adding a partition for location character. We modelled discrete location transitioning between areas throughout *Xci* phylogenetic history using a continuous-time Markov chain (CTMC) process under an asymmetric substitution model with a Bayesian stochastic search variable selection (BSSVS) procedure. States were recoded from countries to greater areas: East Africa (Ethiopia), West Africa (Mali and Senegal), the Caribbean (Martinique, France), North America (United States of America), South America (Argentina and Brazil), East Asia 1 (China and Taiwan), East Asia 2 (Japan), South East Asia 1 (Cambodia, Malaysia, Myanmar, Thailand and Vietnam), South East Asia 2 (Indonesia and Philippines), South Asia 1 (Bangladesh, India and Nepal), South Asia 2 (Iran and Pakistan), West Asia (Oman and Saudi Arabia), Oceania and Pacific (Fiji, Guam, New Zealand and Papua New Guinea), North Indian Ocean islands (Maldives and Seychelles), and South West Indian Ocean islands (Comoros, Mauritius and Rodrigues, Mayotte and La Réunion (France)).

### Pathogenicity-associated genes content analysis

The presence of pathogenicity-associated genes was investigated using a list of 66 type III effectors (T3E) found in *Xanthomonas* (Escalon *et al.* 2013, The Xanthomonas Resource) as well as 24 genes involved in type III secretion system (T3SS) (Buttner 2016) and 54 genes more distantly involved in pathogenicity (S4 Table). Alignments were performed either with BWA-aln or Bowtie 2 (same conditions as above), for herbarium samples and modern strains, respectively. The sequences used to assess homology were the reference strain IAPAR 306 CDS (coding sequences) when available; when not, variants from other *Xci* strains or other *Xanthomonas* CDS were used. Depth was recovered using BEDTools genomecov 2.24.0 (Quinlan *et al.* 2010) and coverage was calculated with R. Genes were considered present if their sequence was covered on more than 75% of its length. The pathogenicity-associated genes content was then projected on BI phylogenetic tree using the gheatmap function of R “ggtree” package (Yu *et al.* 2017).

## Results

### Laboratory procedure & high-throughput sequencing

Thirteen herbarium samples were processed into libraries, using a TruSeq Nano (Illumina) protocol for seven of them, and a home-made BEST protocol for the six others (Carøe *et al.* 2017). Sequencing produced between 56.3 and 365.2 M paired-end reads. Following quality checking and adaptor trimming, reads were merged, presenting insert median length of 32 to 92 nt (mean lengths of 42.1 ± 12.8 to 102.9 ± 45.1 nt).

### Historical genomes reconstruction & ancient DNA damage pattern assessment

Thirteen historical draft *Xci* genomes were reconstructed by mapping processed reads on reference strain IAPAR 306 genome (chromosome, plasmids pXAC33 and pXAC64) (da Silva *et al.* 2002), with one of them, HERB_1937, detailed previously (Campos *et al.* 2021). The proportion of reads mapping to *Xci* reference genome ranged from 0.82% (HERB_1937) to 27.10% (HERB_1922) (Table 2). Respective values of 0.0027, 0.0090 and 0.0196% were obtained for three herbarium *Coffea* sp. control samples.

**Table 2.** Summary of mapping, depth, coverage and damage statistics for the 13 historic *Xci* genomes. SD: standard deviation; nt: nucleotides.

| ID | Protocol | Million reads | Endogenous *Xci* DNA (%) | Chromosome |  |  |  |
| --- | --- | --- | --- | --- | --- | --- | --- |
| | | | | Depth (X) | Coverage at 1X (%) | Insert length (mean ± SD, nt) | Deamination rate at terminal position (%) |
| HERB_1845 | TruSeq Nano | 414.3 | 10.39 | 82.1 | 98.3 | 50.4±23.0 | 3.68 |
| HERB_1884 | TruSeq Nano | 246.8 | 10.48 | 55.6 | 98.1 | 46.2±16.6 | 3.22 |
| HERB_1911 | TruSeq Nano | 365.2 | 2.05 | 32.4 | 98.2 | 69.1±22.3 | 3.70 |
| HERB_1915 | TruSeq Nano | 217.3 | 6.04 | 39.3 | 98.1 | 47.9±17.9 | 2.81 |
| HERB_1937 | TruSeq Nano | 220.9 | 0.82 | 6.2 | 94.6 | 42.7±12.7 | 3.65 |
| HERB_1946 | TruSeq Nano | 262.5 | 7.52 | 64.3 | 98.2 | 57.9±29.5 | 1.81 |
| HERB_1974 | TruSeq Nano | 260.8 | 6.80 | 35.9 | 97.9 | 41.1±9.3 | 2.09 |
| HERB_1852 | BEST | 314.9 | 4.81 | 63.7 | 97.8 | 70.9±43.6 | 1.89 |
| HERB_1854 | BEST | 113.0 | 10.76 | 54.1 | 98.3 | 75.7±42.8 | 1.22 |
| HERB_1859 | BEST | 159.5 | 5.31 | 41.8 | 97.7 | 73.9±33.8 | 1.76 |
| HERB_1865 | BEST | 156.5 | 4.91 | 49.8 | 98.2 | 88.1±39.2 | 1.57 |
| HERB_1922 | BEST | 120.9 | 27.10 | 96.2 | 97.3 | 67.9±29.3 | 1.22 |
| HERB_1963 | BEST | 56.3 | 14.63 | 43 | 98.0 | 74.7±32.3 | 1.25 |

The chromosome sequences displayed a coverage (proportion of reference genome covered) at 1X between 94.6 and 98.2%, with mean depth (average number of mapped reads at each base of the reference genome) of 6.2 (HERB_1937) to 96.2X (HERB_1922) (Table 2). Lower coverages (49.7 to 97.1%) but higher mean depths (17.9 to 130.3X) were obtained for reads mapping to plasmid references (S2 Table).

Ancient DNA typically presents short fragments and cytosine deamination at fragment extremities (Dabney *et al.* 2013). We analyzed such degradation patterns using the dedicated tool mapDamage2 (Jonsson *et al.* 2013). For reads aligning to the chromosome sequence, herbarium samples displayed mean fragments length of 41.1 ± 9.3 to 88.1 ± 39.2 nt, and 3’G>A substitution rates at terminal nucleotides of 1.22 to 3.70%, decreasing exponentially along the DNA molecule for all historical genomes. Modern DNA controls from three *Xci* strains displayed no such decay (Figure 1).

**Fig 1.**
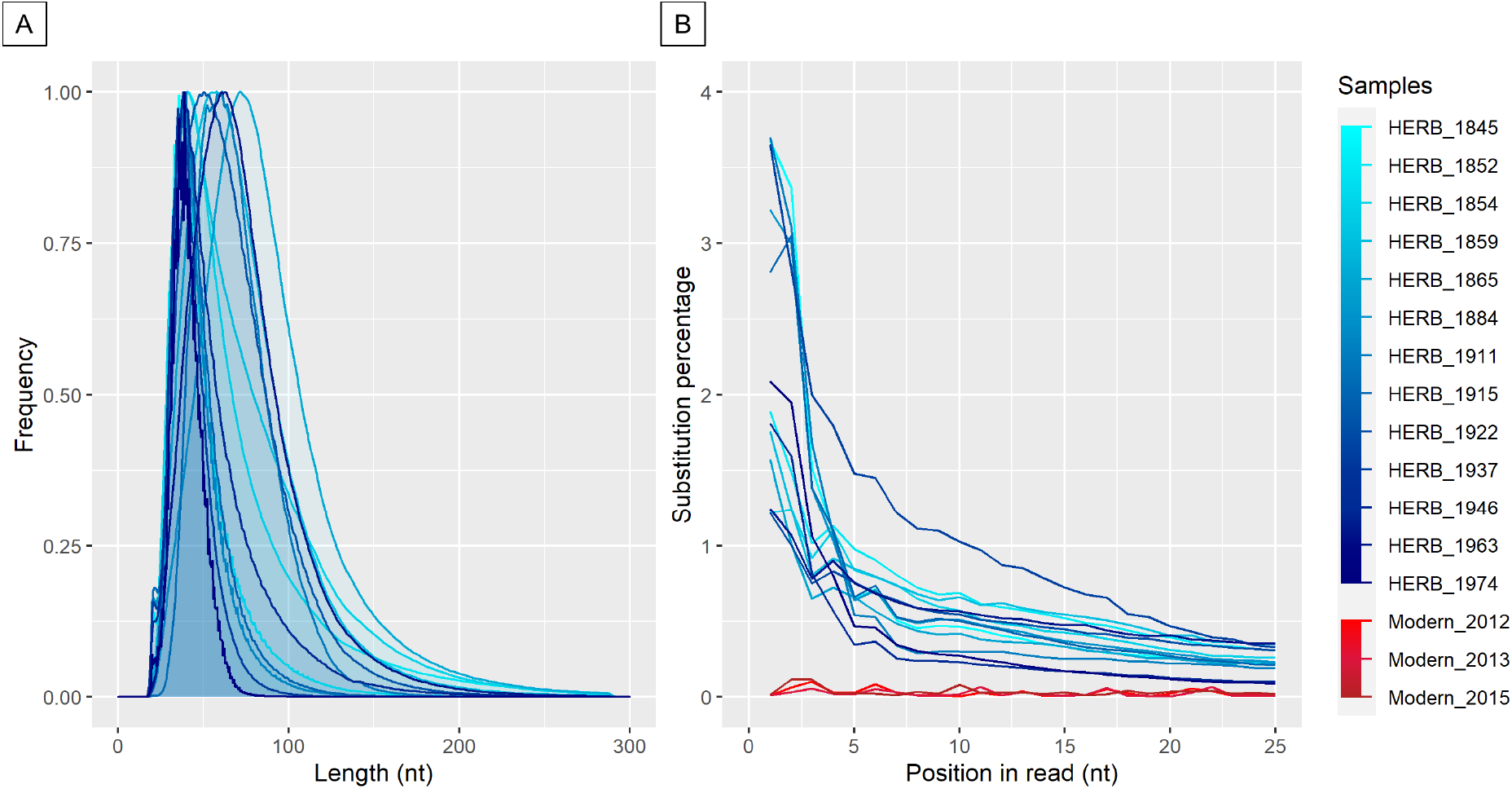
*Post-mortem* DNA damage patterns measured on reads mapping to the *Xci* chromosome. (A) Fragment length distribution (nt: nucleotides; relative frequency in arbitrary units). (B) G>A substitution percentage of the first 25 nucleotides from the 3’ end of the 13 historical genomes (blue lines, light to dark gradient from the oldest to the youngest) and three modern *Xci* strains (red lines, light to dark gradient from the oldest to the youngest).

Similar patterns were obtained for plasmid-like sequences (S1 Fig). Interestingly, significantly lower deamination rates were observed for BEST protocol (95%CI: 1.27-1.64%, *versus* 2.71-3.24% for TruSeq Nano protocol, S2 Fig.). When comparing fragment length categories, the BEST library was enriched in longer fragments (66-140 nt), while the TruSeq Nano library was enriched in smaller fragments (15-40 nt and 41-65 nt) (p-value<0.0001 in all cases, aov function, R package, data not shown). Within each protocol, neither sequence type (aligned on either plasmid or chromosome), nor age, were associated with fragment length differences. More significantly, within each protocol, a time-dependency of the deamination rate was observed (p-value<0.0001, S2 Fig). Finally, within each library construction protocol, deamination rates were significantly higher for plasmid-type sequences (S2 Fig, p<0.0001 for BEST protocol and p=0.0259 for TruSeq Nano protocol). Examining 5’C>T substitution rates gave the same results.

In the first published HERB_1937 herbarium sample, a significantly higher terminal substitution rate was observed for plasmid *versus* chromosome sequences, irrespective of fragment length (Campos *et al.* 2021). In this study, in addition to deamination rates *per se,* the total number of analyzed reads, as well as 11 fragment length classes, were taken into account for each sample, within each library protocol (BEST, TruSeq Nano) and within each sequence type (plasmid, chromosome) (S1 Fig and S2 Table). All the 26 (but one) plasmid-type sequences harbored higher deamination rates, as compared to their respective chromosome-type sequences.

### Phylogenetic reconstruction, dating and ancestral geographic state estimation

Alignment of the chromosome sequence of the 13 historical genomes and 171 modern genomes (17 modern strains from pathotype A*, 4 from pathotype A^W^ and 150 from pathotype A) allowed for the identification of 15,292 high-quality SNPs (Single Nucleotide Polymorphisms). ClonalFrameML identified four major recombining regions (S3 Table), from which 2,285 SNPs were removed from further inferences. On the 13,007 recombination-free SNPs alignment, a paraphyletic outgroup was added, formed of *X. a.* pv. *vasculorum* NCPPB-796 and two strains phylogenetically close to *Xci*, *X. c.* pv. *cajani* LMG558 and *X. c.* pv. *clitoriae* LMG9045. A Maximum-Likelihood (ML) phylogeny was built with RAxML (Stamatakis 2014) and rooted with *X. a.* pv. *vasculorum* (S3 Fig). Strains from each pathotype grouped together and formed distinct clades. Clade A* was at the root of clade *Xci,* while clade A^W^ was a sister-group of clade A. This topology (A*, (A^W^, A)) was highly supported with bootstraps values of 100. Clade A displayed three major, highly supported lineages which we named A1, A2 and A3: lineage A1 corresponds to the main group of strains (lineage A in Patané *et al.* (2019), DAPC1 in Pruvost *et al.* (2014)), contained ten historical specimens, and has a polytomic structure. Lineage A2 contains strains from India and Pakistan but also from Senegal and Mali and corresponds to lineage A2 (Patané *et al.* 2019). Lineage A3 contains seven strains from Bangladesh as well as three historical specimens from Bangladesh, India, and China (Yunnan). Lineages A2 and A3 both contain strains classified as DAPC2 by Pruvost *et al.* (2014). Within these lineages, strains mostly grouped according to their geographic origin, with a few exceptions. Asia is represented in all groups and subgroups. Historical specimens were mainly clustered with modern strains of the same geographical origin.

The ML tree was used to test the presence of temporal signal (*i.e.,* progressive accumulation of mutations over time) within the *Xci* clade using three different tests. The linear regression test between root-to-tip distances and sampling ages displayed a significantly positive slope (value=36.8×10^-6^, adjusted R^2^=0.0235 with a p-value=0.0209) (Fig 2A). Interestingly, such a pattern was also conserved at several other internal nodes (S4 Fig). Second, the BEAST inferred root age and substitution rates of the real *versus* date-randomized datasets exhibited no overlap of the 95% HPD (Highest Posterior Density) (Fig 2B). Finally, the Mantel test displayed no confounding effect (r=-0.73, p-value=0.89) between temporal and genetic structures. To specifically evaluate the contribution of historical genomes to the magnitude of temporal signal, we repeated the above tests on a dataset containing modern genomes only, producing no temporal signal at the *Xci* clade scale (Fig 2A & B).

**Fig 2.**
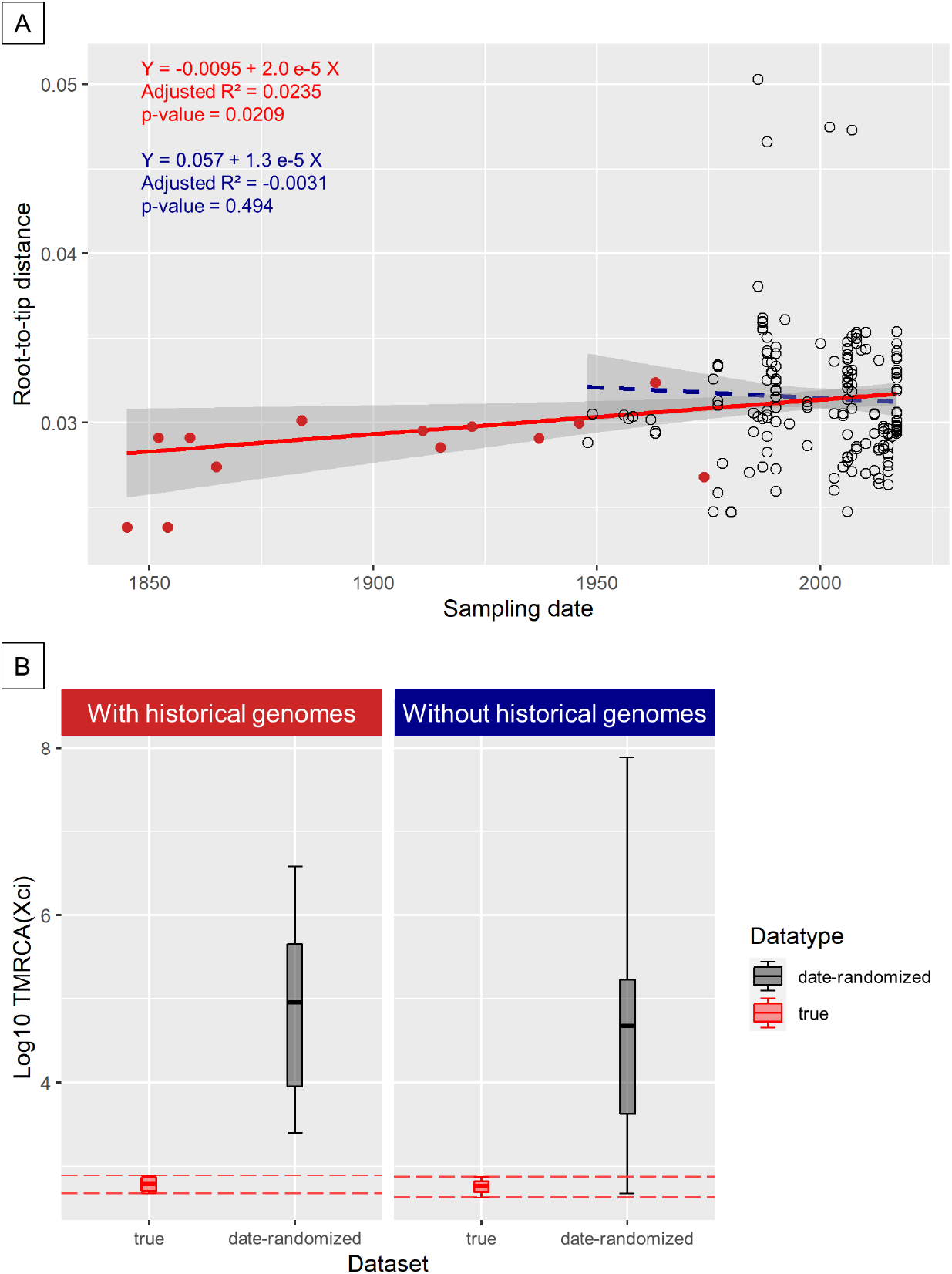
Root-to-tip regression and date-randomization temporal test. (A) Regression lines are plotted in red when integrating historical genomes, and in blue dots when not. Grey areas indicate 95% confidence intervals. Associated values are the regression equation, adjusted R^2^ (Adj R^2^) and p-value (significant if including historical genomes). (B) Evaluating temporal signal in the dataset by date-randomization test showed no overlap between the age of the root estimated from the real dataset (red) and 20 date-randomized datasets (summed up in a single box, black) when historical genomes were included (left). Vertical bars represent 95% Highest Posterior Density intervals.

A Bayesian time-calibrated tree was built with BEAST (Fig 3), and found globally congruent (similar topology and node supports) with the ML tree. The root of the *Xci* clade (node at which *Xci* diversified into numerous pathotypes) was inferred to date to 1218 [95% HPD: 962 - 1437]. We obtained a mean substitution rate of 14.30×10^-8^ [95% HPD: 12.47×10^-8^ - 16.14×10^-8^] per site per year with a standard deviation for the uncorrelated log-normal clock of 0.507 [95% HPD: 0.428 - 0.594], suggesting low rate heterogeneity among tree branches. The inferred dates of other internal nodes of interest, including the MRCA for each of the three pathotypes, as well as for some geographically structured clades, are given in Table 3.

**Fig 3.**
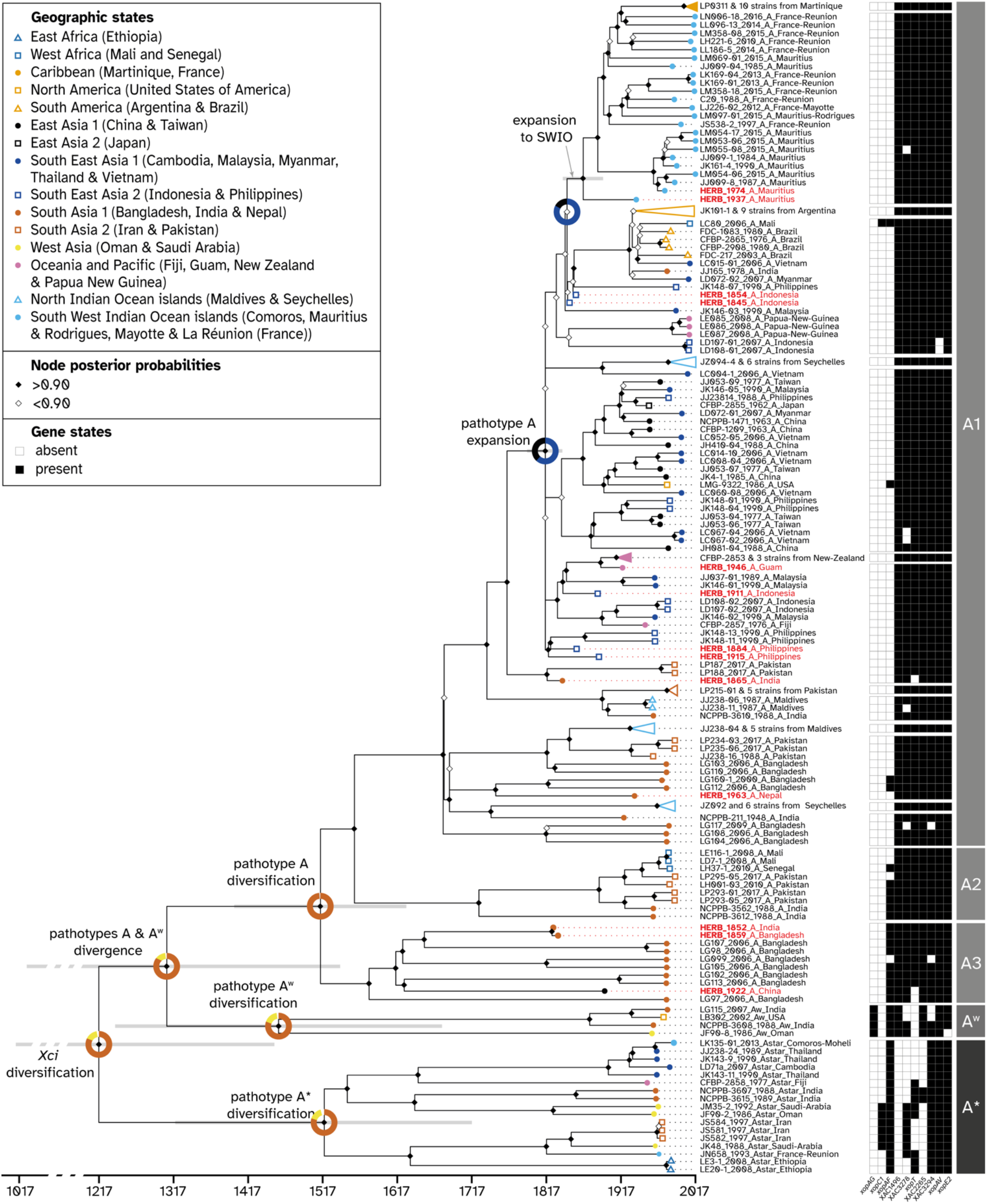
Spatiotemporal Bayesian reconstruction of *Xci* evolutionary history. Dated phylogenetic tree including 13 historical specimens (red labels) and 171 modern strains (black labels) built from 13,007 recombination-free SNPs. Node support values are displayed by diamonds; node bars cover 95% Highest Probability Density of node height. Branch tips are colored according to the sample’s geographic origin. Groups of closely related strains were collapsed for better visibility, details of each group can be found in S2 Fig. Reconstructed ancestral geographic states are represented at some nodes of interest with pie charts representing the posterior probability of geographical regions as the origin of the node. Strain labels include strain name, collection year, pathotype and country of origin. Presence and absence of ten variably-present pathogeny-associated genes is indicated next to strain labels by black squares and white squares, respectively. Pathotypes and lineages are indicated to the right.

**Table 3.** Inferred spatiotemporal data at major nodes. Estimations inferred by Patané *et al.* (2019) (in gray italics) were indicated when possible. Abbreviations: SWIO, South West Indian Ocean. South Asia 1: Bangladesh, India and Nepal; South East Asia 2: Indonesia and Philippines; SWIO islands: Comoros, Mauritius, Rodrigues, Mayotte and La Réunion. * For nodes indicated with an asterisk, reconstructed locations are given for the parent node, i.e., the split between the MRCA of the strains from the indicated geographic areas and its closest relative.

| Node | Date<br>[95% Highest Probability Density] | Location<br>(posterior probability) |
| --- | --- | --- |
| *Xci* origin | -9501 [-13099 – -6446]<br>*[-12052 – -3851]* | South Asia 1 (0.85)<br>*Bangladesh, India, Pakistan* |
| *Xci* diversification | 1218 [962 – 1437]<br>*[-3648 – 285]* | South Asia 1 (0.81)<br>*Bangladesh, India, Pakistan* |
| *Xci* (A<sup>w</sup>, A) divergence | 1305 [1061 – 1511] | South Asia 1 (0.82) |
| *Xci* A* diversification | 1523 [1377 – 1650] | South Asia 1 (0.80)<br>*Iran, Oman, Saudi Arabia* |
| *Xci* A<sup>w</sup> diversification | 1461 [1227 – 1675] | South Asia 1 (0.80)<br>*Bangladesh, India, Pakistan* |
| *Xci* A diversification | 1524 [1390 – 1637] | South Asia 1 (0.99)<br>*Bangladesh, India, Pakistan* |
| *Xci* A - (A2, A1) divergence | 1570 [1408 – 1684] | South Asia 1 (1.00)<br>*Bangladesh, India, Pakistan* |
| *Xci* A1 diversification | 1696 [1607 – 1768] | South Asia 1 (1.00)<br>*Bangladesh, India, Pakistan* |
| *Xci* A expansion (polytomy) | 1829 [1799 – 1845] | South East Asia 2 (0.62) |
| *Xci* A - New Zealand MRCA* | 1925 [1896 – 1949] | South East Asia 2 (0.79) |
| *Xci* A - South America MRCA* | 1930 [1907 – 1951] | South East Asia 2 (0.96) |
| *Xci* A - SWIO islands MRCA* | 1868 [1844 – 1899] | South East Asia 2 (0.83) |
| *Xci* A - Caribbean MRCA* | 2000 [1991 – 2008] | SWIO islands (1.00) |

To date the origin of *Xci, i.e.,* the date at which it diverged from its closest relatives, a secondary alignment was realized with the 13 historical sequences, the 171 modern sequences and the three outgroup sequences. A total of 209,306 chromosomal high-quality SNPs was found, 198,249 of which were outside recombining regions. The presence of temporal signal was tested as previously described. On this dataset, temporal signal was still present at the root of *Xci* clade but the signal disappeared at the external node connecting the outgroup, with *X. c.* pv. *cajani* as *Xci* closest relative (root-to-tip regression test: slope value=-4.6×10^-6^, adjusted R^2^=0.0006 with a p-value=0.293; date-randomization test: overlapping of the 95% HPD of the root age (S5 Fig); Mantel’s test: r=0.04, p-value=0.09). As the prerequisite for tip-dating was not met, we realized a rate-dating analysis integrating the mean substitution rate inferred in the tip-dating calibration. The node date at which *Xci* split from other *Xanthomonas citri* pathovars was inferred to −9501 [95% HPD: -13099 - -6446].

In order to infer ancestral location state at nodes, geographic signal in the dataset must first be detected. By measuring the association index between topology and the location trait data on both the real tree and 1,000 location-randomized trees, our results highlighted the existence of a non-random association between spatial and phylogenetic structure (p-value<0.0001). A discrete phylogeographic analysis was therefore run with BEAST (Fig 3, Table 3), inferring a highly-supported South Asia 1 (Bangladesh, India and Nepal) origin where *Xci* split from its *Xanthomonas citri* relatives and then diversified through its three lineages A*, A^W^ and A. All of them, comprising A1, A2 and A3 lineages included in A, were also inferred to have diversified in the same South Asia 1 region. In the lineage A1, the polytomy was inferred to have a Southeast Asia 2 (Indonesia or Philippines) origin (node support of 1, state posterior probability of 0.62). Southeast Asia 2 origin was also inferred for the lineages composed of: 1) all New Zealand strains (herbarium specimen HERB_1946_A_Guam at the root); 2) all Argentina strains (with HERB_1854_A_Indonesia and HERB_1845_A_Indonesia branching closely) and 3) nearly all strains from the SWIO islands and from Martinique (with HERB_1974_A_Mauritius and HERB_1937_A_Mauritius at their root).

### Pathogenicity-associated genes content

We investigated the presence or absence of 144 pathogenicity-associated genes (S4 Table) under a mapping approach. The twenty-four genes coding for the type III secretion system (T3SS) were present in all *Xci* strains. Of the 66 T3E genes tested, we found 32 present and 28 absent in all *Xci* strains, whereas 6 were of variable presence. In addition, among 54 genes potentially involved in virulence but not necessarily dependent on the T3SS, only 4 were of variable presence in *Xci,* the 50 others being systematically present. The ten identified variable virulence factors were *xopAF, xopAG, xopAV, xopC1, xopE, xopT,* XAC1496, XAC2265/*helD*, XAC3278 and XAC3294 (Fig 3 & S5 Table). In general variation was localized in the “distant” branches of the tree, and concerned strains from pathotypes A* and A^W^. Conversely, strains belonging to pathotype A were rather homogenous in their pathogenicity factor contents.

## Discussion

In this study, we successfully reconstructed the genome of 13 *Xci* historical strains from herbarium material collected between 1845 and 1974, which we compared with a set of 171 modern genomes representative of the bacterial global diversity, 57 of them having been specifically generated for this study. A better understanding of *Xci* evolutionary history is a subject of great interest since it may help deciphering how bacterial pathogens specialize on their hosts and diversify while expanding their geographical range.

At a molecular level, the analysis of historical genomes was highly informative. First of all, assessment of *post-mortem* DNA degradation patterns specific to ancient DNA, such as fragmentation and deamination, confirmed the historical nature of the reconstructed genomes. Terminal deamination values, consistent with those of Weiss *et al.* (2016) herbarium samples (roughly, between 1.5 and 5.0%), were higher for plasmidic versus chromosomal sequences, possibly due to chromosome-specific cytosine methylation patterns (discussed in (Campos *et al.* 2021)). Different deamination values were also observed between library protocols. The usage of specific DNA polymerases incapable of recovering uracils during library amplification (Kistler *et al.* 2017), or specific bead-purifications resulting in longer fragment size enrichment (as observed in our study with the BEST protocol), could lead to the observation of lower deamination rates. Finally, for each protocol we showed for the first time an age-dependent deamination for bacterial DNA from herbarium material, as described elsewhere for nuclear and chloroplastic plant DNA (Weiss *et al.* 2016).

Adopting a shotgun-based deep sequencing strategy revealed between 0.8% to 25.5% of endogenous *Xci* DNA amongst the 13 historical samples, a wide variation falling in the range of previous studies that attempted to retrieve non vascular pathogen DNA from infected herbarium leaves (Martin *et al.* 2013, Yoshida *et al.* 2015, Yoshida *et al.* 2013). We aligned the 13 historical genomes with 171 modern representatives of the bacterial global diversity, and built a phylogenetic tree from the chromosome-wide non-recombining SNPs. This tree confidently associated the pathotypes to be monophyletic groups, and displayed a (A*, (A^W^, A)) topology, as previously reported from genome-wide SNPs (Gordon *et al.* 2015) and unicopy gene families analyses (Patané *et al.* 2019). Interestingly, the relationships inside the pathotype A (A3, (A2, A1)) agreed with the former analysis but not the latter, whose discrepancy could be explained by the under-representation of A3 lineage (only LG98_2006_A_Bangladesh present). Globally, the observed geographic clustering inside the pathotype A clade is consistent with previous studies (Patané *et al.* 2019, Pruvost *et al.* 2014, Richard *et al.* 2021). Clade A1 had a somehow polytomic structure (most lineages seem to have branched from a single ancestor at the same moment). This can either be the result of a rapid and synchronous expansion of *Xci* in contrasted environments, leading to the divergence and maintenance of multiple lineages, or of an artefact of building topologies with insufficient data, when lack of information does not allow to differentiate distinct divergence events (Lin *et al.* 2011). Our datation (see below) of the beginning of the A1 diversification fits with the first scenario, but as many nodes in the A1 cluster are not well supported (white diamonds in Fig 3) we cannot conclude.

The presence of temporal structure is an essential prerequisite to perform tip-calibrated inferences (Drummond *et al.* 2002, Drummond *et al.* 2003, Rieux *et al.* 2016a). While the dataset containing contemporary genomes only (1948 - 2017) did not reveal the existence of any measurably evolving population, inclusion of the 13 historical genomes (1845 - 1974) brought the required temporal signal within the *Xci* clade. This allowed us building a time-calibrated phylogeny without making any underlying assumption on the age of any node in the tree, nor on the rate of evolution and proposing new evolutionary scenarios for the origin and diversification of the pathogen. To our knowledge, this is the first study attempting to elucidate the evolutionary history of a bacterial crop pathogen at such a global scale using herbarium specimens. Previous ones did not use historical strains (Patané *et al.* 2019), focused on a recent and local emergence (Campos *et al.* 2021), or were limited by the exploitation of a few partial genetic markers only (Li *et al.* 2007).

We inferred a mean substitution rate of 14.30×10^-8^ [95% HPD: 12.47×10^-8^ - 16.14×10^-8^] substitutions per site per year, a value ~1.5x faster than the one (9.4×10^-8^ [95% HPD: 7.3×10^-8^ - 11.4×10^-8^]) obtained by our team on a single lineage within the pathotype A clade of *Xci,* at the local scale of the South West Indian Ocean islands (Campos *et al.* 2021). We dated the MRCA of all *Xci* strains, the node leading to bacterial diversification, to the beginning of the 13^th^ century (1218 [95% HPD: 962 - 1437]), a much more recent timespan than the one [-3648 - 285] inferred previously (Patané *et al.* 2019). The discrepancy between those estimates probably arises from differences in the considered molecular dating methodologies. Indeed, their molecular clock was calibrated by applying a prior on both the rate of evolution (estimated on house-keeping genes of non-*Xci Xanthomonas* species) and the age of a node external to the *Xci* clade (indirectly deriving from the same rate of evolution), while the methodology used in our work makes use of the age of the strains only, a method shown to yield far more accurate and robust estimates (Ho *et al.* 2008, Rieux *et al.* 2016a, Rieux *et al.* 2014). In addition of dating the emergence of the three *Xci* lineages A*, A^W^ & A, we also inferred the ones of geographically structured lineages such as the one in New Zealand in 1925 [95% HPD: 1896 - 1949], in South America in 1930 [95% HPD: 1907 - 1951] or in Martinique in 2000 [95% HPD: 1991 - 2008] with values always predating disease first reports made in 1937 (Dye 1969), 1957 (Rossetti 1977), and 2014 (Richard *et al.* 2016), respectively. Finally, as the divergence between a pathogen and its closest known relative places a maximum bound on the timing of its emergence (Duchêne *et al.* 2020), we included three outgroup sequences, of which *X. c.* pv*. cajani,* a pathogen of the *Fabaceae* plant family, was the first to branch out of the *Xci* clade. As the inclusion of divergent outgroup genomes precluded the application of tip-dating methodology we extrapolated the rate of substitution previously estimated within the *Xci* clade to date the split between *Xci* and *X. c.* pv*. cajani* to -9501 [95% HPD: -13099 - - 6446], a value which partly overlap with the inferred interval of [-12052 - -3851] found by Patané *et al.* (2019), although on a more restrained length of time.

Our phylogeographic analysis inferred an origin and diversification of *Xci* in an area of South Asia neighboring to Bangladesh, India and Nepal, consistently with previous reconstructions and estimations based on genetic diversity (Gordon *et al.* 2015, Patané *et al.* 2019, Pruvost *et al.* 2014). It also corresponds to the area of origin of the *Citrus* genus, which is believed to have emerged within the southern foothills of the Himalayas (including Assam, Western Yunnan and Northern Myanmar) 6 to 8 Mya (Wu *et al.* 2018). Our temporal calibrations indicate a mean age of 11.5 ky for the origin of *Xci,* a period which coincides with the beginning of the Holocene (~-9700 - present) (Walker *et al.* 2009) following the Bølling-Allerød warming global event (~-12700 - -10900) (Rasmussen *et al.* 2006). Such warmer and wetter climates could have facilitated plant expansion into new areas previously occupied by ice such as the mountainous regions and the northern parts of South Asia (Staubwasser *et al.* 2006). This was followed by the development of societies and agriculture in Northern India and in China during which movements of plants between and outside these regions may have gathered favorable conditions for *Xci* emergence on *Citrus* through bacterial host jump, as previously proposed (Patané *et al.* 2019). The diversification of *Xci* was dated to the early 13^th^ century in its area of origin, which was crossed at the time by the Southern Silk Road (Talon *et al.* 2020), linking Eastern and Western civilizations through trading. The westward commerce of goods, including citrus, which have been found in Mediterranean countries since -500 (Zech-Matterne *et al.* 2017), as well as the breeding of citrus varieties for cooking and for raw eating (Talon *et al.* 2020), could have dispersed and isolated the pathogen into its three known pathotypes. More recent global changes observed between the 17^th^ and 20^th^ centuries, such as the spice trade and the development of a worldwide colonial agriculture might also be important factors in the apparently intense diversification of genotypes (and their global spread) (Campos *et al.* 2021).

We based our analysis of the contents in virulence-related genes on lists of proven and hypothetical factors, a large part of which are T3SS elements and T3E. As expected, and as reported previously all *Xci* strains contained genes for a full type III secretion apparatus and for a large number of T3E: with 32 genes it is larger than previously reported (Escalon *et al.* 2013), mostly because new (often hypothetical) genes were described since then (Teper *et al.* 2016). Variation was detected for six T3E among *Xci* (detailed below). Factors not related to the T3SS were also assessed: we confirmed that most are present in all of the strains, with variation concerning four of them.

Inconsistencies with other works can be due to our identification strategy (mapping on more than 75% of the CDS). For example, XAC3278 (XAC_RS16605) was considered to be present in all strains by Patané *et al.* (2019), but we found that there can be two paralogs of the gene. One is distant (65% nucleotidic identity) from the functionally demonstrated copy but is present in all strains, while the other, 100% homologous to XAC3278, is only present in some strains. Similarly, we found variability in the length of homologues to XAC2265 (XAC_RS11510) between strains, some being truncated of more than 30% of their length. Finally, *xopF1* is pseudogenized, and probably not functional (Jalan *et al.* 2013). These results confirm the general tendency of “core sets of genes” to be reduced to nothing when enough strains are analyzed and that the concept of core effectome is not especially relevant for *Xanthomonas* at the genus level (Roux *et al.* 2015).

Variation in some virulence factors (for example, *xopE2, xopAV,* XAC3294) was anecdotal, concerning only one or a few strains. In others cases their distribution was found to be more or less clade-dependent: *xopAG* is present only in pathotype A^W^ strains, as expected (Rybak *et al.* 2009). *XopC1* is present in a few A* strains. XAC1496 is specifically absent from all A* strains, suggesting an acquisition during the pathotype differentiation process. *XopAF* is present only in the deeply rooted branches of the tree, suggesting a loss of the effector for strains of clade A1. XAC2265 is absent from A* strains except for two of them, suggesting reacquisition of this factor. Conversely, XAC3278 and *xopT* seem to have been lost several times in the recent past.

Herbarium samples were not different in their effector contents from their close phylogenetic relatives, indicating that most of the effector shuffling process occurred before the 1850s. This is consistent with evolutionary hypotheses that postulate that fundamental factors of pathogenicity (in our case the type III secretion apparatus and probably many non-T3 factors) are acquired early in the evolutionary history of plant-associated bacteria, providing a general adaptation to interactions with plants, with a subsequent host specialization (and differentiation in further clades) correlated to variable T3E contents and coevolutionary arms race (Hajri *et al.* 2009, Ma *et al.* 2006, Merda *et al.* 2017).

Our work presents two main limitations. First, although with 163 genomes our dataset displayed the best representation of pathotype A genetic diversity published to date, the reconstructed phylogenetic tree exhibits high level of imbalance (with only 17 and 4 A* and A^W^ genomes, respectively), a property previously shown to lead to reduced accuracy or precision of phylogenetic timescale estimates (Duchêne *et al.* 2015a). Bias in representation of populations, such as overrepresentation of one compared to the others or the absence of representants from the true founder lineage, can also lead to the reconstruction of ancestral state tending to correspond to the oversampled population rather than the true founder lineage (Rasmussen *et al.* 2021). Although this feature arises from the fact that *Xci* worldwide expansion mostly involved pathotype A strains (Pruvost *et al.* 2014), future work should aim to better characterize the genomic diversity of A* and A^W^ strains. Secondly, as gene content variation analysis was performed by mapping reads to reference sequences, we were unable to identify potential genomic rearrangements among strains, a process known to be frequent within *Xanthomonas* species (Jacques *et al.* 2016, Merda *et al.* 2017, Richard *et al.* 2022). Similarly, this approach impeded us from identifying genetic content absent from the reference sequences. To overcome those limitations and better recover pathotype-specific genes, comparative genomic analysis based on *de novo* assembly and/or without *a priori* on the targeted genes would be interesting to perform.

To conclude, our study emphasizes how historical genomes from herbarium samples can provide a wealth of genetic and temporal information on bacterial crop pathogens evolution. Similar studies could be applied to other plant pathogens to infer the temporal dynamic of their populations and elucidate their evolutionary history with more resolutive estimations, which in turn may provide clues to improve disease monitoring and achieve sustainable control.

## Acknowledgments

We are grateful to P. Lefeuvre, D. Richard, F. Balloux, V. Llaurens, R. Debruyne & Á. Pérez-Quintero for valuable comments and discussions. We thank L Bui Thi Ngoc (SOFRI, Viet Nam), B. Canteros (INTA, Argentina), B. Carter (FERA, UK), R. Davis (NASQ, Australia), A. Hamza (INRAPE, the Comoros), G. Johnson (Horticulture4Development, Australia), N. Le Mai (PPRI, Viet Nam), W. Wu (National Chung Hsing University, Taiwan) and M. Zakria (NARC, Pakistan) for providing citrus canker lesions and/or bacterial strains. We also thank Meghann Toner, collections management, Department of Botany, National Museum of Natural History, Washington, D.C., USA for allowing us to consult and sample Citrus specimens at the U.S. National Herbarium. We also thank L Bui Thi Ngoc, B. Canteros, B. Carter, R. Davis, A. Hamza, G. Johnson, N. Le Mai, W. Wu and M. Zakria for providing diseased citrus material and/or bacterial strains. We would also like to acknowledge herbarium curators who allowed us to collect samples not included in this study. Collection of any plant material used in this study complies with institutional, national, and international guidelines. Computational work was performed on the CIRAD - UMR AGAP HPC data center of the South Green bioinformatics platform (http://www.southgreen.fr/) and MESO@LR-Platform at the University of Montpellier (https://hal.umontpellier.fr/MESO). This work was conducted on the Plant Protection Platform (3P, IBISA).

## Data, script and code availability

The authors confirm that all data used in this study are fully available without restriction. Both historical and modern raw reads were deposited to the Sequence Read Archive (under accession numbers listed in S1 Table). Accession numbers of any previously published data used in this study are also listed in S1 Table.

## Supplementary information

S1 Fig. Post-mortem DNA damage patterns on Xci plasmids pXAC33 and pXAC64.

S2 Fig. Deamination rate (in percentage) at terminal position from the 3’ end (G to A substitutions) as a function of the collection year for the 13 Xci genomes reconstructed from herbarium specimens.

S3 Fig. Maximum Likelihood (ML) phylogenetic tree of historical and modern Xci genomes

S4 Fig. Root-to-tip regression temporal test visualised on online tool PhyloStemS

S5 Fig. Root-to-tip regression and date-randomisation temporal test results when performed on the dataset including outgroups

S1 Table. General characteristics of the historical and modern strains of the study

S2 Table. Summary of mapping, depth, coverage and damage statistics for the 13 historic Xci plasmids pXAC33 and pXAC64.

S3 Table. Recombining regions among 185 historical or modern Xci strains

S4 Table. List and presence status of 144 pathogenicity-associated genes investigated among 184 historical or modern Xci strains

S5 Table. Coverage (in percentage) of the 10 pathogenicity-associated genes of variable presence investigated among 184 historical or modern Xci strains

## References

Al Rwahnih M, Rowhani A & Golino D (2015). First report of grapevine red blotch-associated virus in archival grapevine material from Sonoma County, California. Plant Dis. 99(6): 895. https://doi.org/10.1094/PDIS-12-14-1252-PDN

Anderson PK, Cunningham AA, Patel NG, Morales FJ, Epstein PR & Daszak P (2004). Emerging infectious diseases of plants: pathogen pollution, climate change and agrotechnology drivers. Trends Ecol. Evol. 19(10): 535–544. https://doi.org/10.1016/j.tree.2004.07.021

Ayres DL, Darling A, Zwickl DJ, Beerli P, Holder MT, Lewis PO, Huelsenbeck JP, Ronquist F, Swofford DL, Cummings MP, Rambaut A & Suchard MA (2012). BEAGLE: An application programming interface and high- performance computing library for statistical phylogenetics. Syst. Biol. 61(1): 170–173. https://doi.org/10.1093/sysbio/syr100

Bernardes MFF, Pazin M, Pereira LC & Dorta DJ (2015). Impact of pesticides on environmental and human health. In “Toxicology studies-cells, drugs and environment”. (Andreazza AC & Scola G, Eds.) Rijeka, Croatia, InTech. pp 195–233. https://doi.org/10.5772/58714.

Biek R, Pybus OG, Lloyd-Smith JO & Didelot X (2015). Measurably evolving pathogens in the genomic era. Trends Ecol. Evol. 30(6): 306–313. https://doi.org/10.1016/j.tree.2015.03.009

Bolger AM, Lohse M & Usadel B (2014). Trimmomatic: a flexible trimmer for Illumina sequence data. Bioinformatics 30(15): 2114–2120. https://doi.org/10.1093/bioinformatics/btu170

Broad Institute. Picard Tools. Accessed November 2020, http://broadinstitute.github.io/picard/

Buttner D (2016). Behind the lines-actions of bacterial type III effector proteins in plant cells. FEMS Microbiol. Rev. 40(6): 894–937. https://doi.org/10.1093/femsre/fuw026

Campos PE, Groot Crego C, Boyer K, Gaudeul M, Baider C, Richard D, Pruvost O, Roumagnac P, Szurek B, Becker N, Gagnevin L & Rieux A (2021). First historical genome of a crop bacterial pathogen from herbarium specimen: Insights into citrus canker emergence. PLoS Pathog. 17(7): e1009714. https://doi.org/10.1371/journal.ppat.1009714

Carøe C, Gopalakrishnan S, Vinner L, Mak SST, Sinding MHS, Samaniego JA, Wales N, Sicheritz-Pontén T & Gilbert MT (2017). Single-tube library preparation for degraded DNA. Methods Ecol. Evol. 9: 410–419. https://doi.org/10.1111/2041-210X.12871

da Silva AC, Ferro JA, Reinach FC, Farah CS, Furlan LR, Quaggio RB, Monteiro-Vitorello CB, Van Sluys MA, Almeida NF, Alves LM, do Amaral AM, Bertolini MC, Camargo LE, Camarotte G, Cannavan F, Cardozo J, Chambergo F, Ciapina LP, Cicarelli RM, Coutinho LL, Cursino-Santos JR, El-Dorry H, Faria JB, Ferreira AJ, Ferreira RC, Ferro MI, Formighieri EF, Franco MC, Greggio CC, Gruber A, Katsuyama AM, Kishi LT, Leite RP, Lemos EG, Lemos MV, Locali EC, Machado MA, Madeira AM, Martinez-Rossi NM, Martins EC, Meidanis J, Menck CF, Miyaki CY, Moon DH, Moreira LM, Novo MT, Okura VK, Oliveira MC, Oliveira VR, Pereira HA, Rossi A, Sena JA, Silva C, de Souza RF, Spinola LA, Takita MA, Tamura RE, Teixeira EC, Tezza RI, Trindade dos Santos M, Truffi D, Tsai SM, White FF, Setubal JC & Kitajima JP (2002). Comparison of the genomes of two *Xanthomonas* pathogens with differing host specificities. Nature 417(6887): 459–463. https://doi.org/10.1038/417459a

Dabney J, Meyer M & Paabo S (2013). Ancient DNA damage. Cold Spring Harb. Perspect. Biol. 5(7): a012567. https://doi.org/10.1101/cshperspect.a012567

Dark P & Gent H (2001). Pests and diseases of prehistoric crops: a yield ‘honeymoon’ for early grain crops in Europe? Oxford J. Archaeol. 20(1): 59–78. https://doi.org/10.1111/1468-0092.00123

DePristo MA, Banks E, Poplin R, Garimella KV, Maguire JR, Hartl C, Philippakis AA, del Angel G, Rivas MA, Hanna M, McKenna A, Fennell TJ, Kernytsky AM, Sivachenko AY, Cibulskis K, Gabriel SB, Altshuler D & Daly MJ (2011). A framework for variation discovery and genotyping using next-generation DNA sequencing data. Nat. Genet. 43(5): 491–498. https://doi.org/10.1038/ng.806

Didelot X & Wilson DJ (2015). ClonalFrameML: Efficient inference of recombination in whole bacterial genomes. PLoS Comput. Biol.11 (2). https://doi.org/10.1371/journal.pcbi.1004041

DOE Joint Genome Institute. BBTools. Accessed November 2020, https://jgi.doe.gov/data-and-tools/software-tools/

Doizy A, Prin A, Cornu G, Chiroleu F & Rieux A (2020). Phylostems: a new graphical tool to investigate temporal signal of heterochronous sequences at various evolutionary scales. bioRxiv. https://doi.org/10.1101/2020.10.19.346429

Drummond AJ, Nicholls GK, Rodrigo AG & Solomon W (2002). Estimating mutation parameters, population history and genealogy simultaneously from temporally spaced sequence data. Genetics 161(3): 1307–1320. https://doi.org/10.1093/genetics/161.3.1307

Drummond AJ, Pybus OG, Rambaut A, Forsberg R & Rodrigo AG (2003). Measurably evolving populations. Trends Ecol. Evol. 18(9): 481–488. https://doi.org/10.1016/S0169-5347(03)00216-7

Drummond AJ & Rambaut A (2007). BEAST: Bayesian evolutionary analysis by sampling trees. BMC Evol. Biol. 7: 214. https://doi.org/10.1186/1471-2148-7-214

Duchêne D, Duchêne S & Ho SYW (2015a). Tree imbalance causes a bias in phylogenetic estimation of evolutionary timescales using heterochronous sequences. Mol. Ecol. Resour. 15(4): 785–794. https://doi.org/10.1111/1755-0998.12352

Duchêne S, Duchêne D, Holmes EC & Ho SYW (2015b). The performance of the date-randomization test in phylogenetic analyses of time-structured virus data. Mol. Biol. Evol. 32(7): 1895–1906. https://doi.org/10.1093/molbev/msv056

Duchêne S, Ho SYW, Carmichael AG, Holmes EC & Poinar H (2020). The recovery, interpretation and use of ancient pathogen genomes. Curr. Biol. 30(19): R1215–1231. https://doi.org/10.1016/j.cub.2020.08.081

Dye DW (1969). Eradicating citrus canker from New Zealand. N. Z. J. Agric. Res. 118(2): 20–21.

Escalon A, Javegny S, Vernière C, Noël LD, Vital K, Poussier S, Hajri A, Boureau T, Pruvost O, Arlat M & Gagnevin L (2013). Variations in type III effector repertoires, pathological phenotypes and host range of *Xanthomonas citri* pv. *citri* pathotypes. Mol. Plant Pathol. 14(5): 483–496. https://doi.org/10.1111/mpp.12019

Gordon JL, Lefeuvre P, Escalon A, Barbe V, Curveiller S, Gagnevin L & Pruvost O (2015). Comparative genomics of 43 strains of *Xanthomonas citri* pv. *citri* reveals the evolutionary events giving rise to pathotypes with different host ranges. BMC Genomics 16: 1098. https://doi.org/10.1186/s12864-015-2310-x

Gottwald TR, Graham JH & Schubert TS (2002). Citrus canker: The pathogen and its impact. Plant Health Prog. 3(1): 15. https://doi.org/10.1094/php-2002-0812-01-rv

Graham JH, Gottwald TR, Cubero J & Achor DS (2004). *Xanthomonas axonopodis* pv. *citri:* factors affecting successful eradication of citrus canker. Mol. Plant Pathol. 5(1): 1–15. https://doi.org/10.1046/j.1364-3703.2004.00197.x

Hajri A, Brin C, Hunault G, Lardeux F, Lemaire C, Manceau C, Boureau T & Poussier S (2009). A “repertoire for repertoire” hypothesis: repertoires of type three effectors are candidate determinants of host specificity in *Xanthomonas*. PLoS One 4(8): e6632. https://doi.org/10.1371/journal.pone.0006632

Ho SYW, Saarma U, Barnett R, Haile J & Shapiro B (2008). The effect of inappropriate calibration: three case studies in molecular ecology. PLoS One 3(2). https://doi.org/10.1371/journal.pone.0001615

Jacques MA, Arlat M, Boulanger A, Boureau T, Carrère S, Cesbron S, Chen NWG, Cociancich S, Darrasse A, Denancé N, Fischer-Le Saux M, Gagnevin L, Koebnik R, Lauber E, Noël LD, Pieretti I, Portier P, Pruvost O, Rieux A, Robène I, Royer M, Szurek B, Verdier V & Vernière C (2016). Using ecology, physiology and genomics to understand host specificity in *Xanthomonas:* French Network on Xanthomonads (FNX). Annu. Rev. Phytopathol. 54(1): 163–187. https://doi.org/10.1146/annurev-phyto-080615-100147

Jalan N, Kumar D, Andrade MO, Yu F, Jones JB, Graham JH, White FF, Setubal JC & Wang N (2013). Comparative genomic and transcriptome analyses of pathotypes of *Xanthomonas citri* subsp. *citri* provide insights into mechanisms of bacterial virulence and host range. BMC Genomics 14: 551. https://doi.org/10.1186/1471-2164-14-551

Jonsson H, Ginolhac A, Schubert M, Johnson PL & Orlando L (2013). mapDamage2.0: fast approximate Bayesian estimates of ancient DNA damage parameters. Bioinformatics 29(13): 1682–1684. https://doi.org/10.1093/bioinformatics/btt193

Jun G, Wing MK, Abecasis GR & Kang HM (2015). An efficient and scalable analysis framework for variant extraction and refinement from population-scale DNA sequence data. Genome Res. 25(6): 918–925. https://doi.org/10.1101/gr.176552.114

Kistler L, Ware R, Smith O, Collins M & Allaby RG (2017). A new model for ancient DNA decay based on paleogenomic meta-analysis. Nucleic Acids Res. 45(11): 6310–6320. https://doi.org/10.1093/nar/gkx361

Lanave C, Preparata G, Saccone C & Serio G (1984). A new method for calculating evolutionary substitution rates. J. Mol. Evol. 20(1): 86–93. https://doi.org/10.1007/BF02101990

Langmead B & Salzberg SL (2012). Fast gapped-read alignment with Bowtie 2. Nat. Methods 9(4): 357–U354. https://doi.org/10.1038/Nmeth.1923

Li H & Durbin R (2009). Fast and accurate short read alignment with Burrows-Wheeler transform. Bioinformatics 25(14): 1754–1760. https://doi.org/10.1093/bioinformatics/btp324

Li W, Song Q, Brlansky RH & Hartung JS (2007). Genetic diversity of citrus bacterial canker pathogens preserved in herbarium specimens. P. Natl. Acad. Sci. USA 104(47): 18427–18432. https://doi.org/10.1073/pnas.0705590104

Lin GN, Zhang C & Xu D (2011). Polytomy identification in microbial phylogenetic reconstruction. BMC Syst Biol 5 Suppl 3: S2. https://doi.org/10.1186/1752-0509-5-S3-S2

Ma W, Dong FF, Stavrinides J & Guttman DS (2006). Type III effector diversification via both pathoadaptation and horizontal transfer in response to a coevolutionary arms race. PLoS Genet. 2(12): e209. https://doi.org/10.1371/journal.pgen.0020209

Malmstrom CM, Martin MD & Gagnevin L (2022). Exploring the emergence and evolution of plant pathogenic microbes using historical and paleontological sources. Annu. Rev. Phytopathol. 60(1): 187–209. https://doi.org/10.1146/annurev-phyto-021021-041830

Malmstrom CM, Shu R, Linton EW, Newton L & Cook MA (2007). Barley yellow dwarf viruses (BYDVs) preserved in herbarium specimens illuminate historical disease ecology of invasive and native grasses. J. Ecol. 95(6): 1153–1166. https://doi.org/10.1111/j.1365-2745.2007.01307.x

Martin MD, Cappellini E, Samaniego JA, Zepeda ML, Campos PF, Seguin-Orlando A, Wales N, Orlando L, Ho SY, Dietrich FS, Mieczkowski PA, Heitman J, Willerslev E, Krogh A, Ristaino JB & Gilbert MT (2013). Reconstructing genome evolution in historic samples of the Irish potato famine pathogen. Nat. Commun. 4: 2172. https://doi.org/10.1038/ncomms3172

McCann HC (2020). Skirmish or war: the emergence of agricultural plant pathogens. Curr. Opin. Plant Biol. 56: 147–152. https://doi.org/10.1016/j.pbi.2020.06.003

Merda D, Briand M, Bosis E, Rousseau C, Portier P, Barret M, Jacques MA & Fischer-Le Saux M (2017). Ancestral acquisitions, gene flow and multiple evolutionary trajectories of the type three secretion system and effectors in *Xanthomonas* plant pathogens. Mol. Ecol. 26(21): 5939–5952. https://doi.org/10.1111/mec.14343

Mira A, Pushker R & Rodriguez-Valera F (2006). The Neolithic revolution of bacterial genomes. Trends Microbiol. 14(5): 200–206. https://doi.org/10.1016/j.tim.2006.03.001

Murray GGR, Wang F, Harrison EM, Paterson GK, Mather AE, Harris SR, Holmes MA, Rambaut A & Welch JJ (2016). The effect of genetic structure on molecular dating and tests for temporal signal. Methods Ecol. Evol. 7(1): 80–89. https://doi.org/10.1111/2041-210x.12466

Parker J, Rambaut A & Pybus OG (2008). Correlating viral phenotypes with phylogeny: Accounting for phylogenetic uncertainty. Infect. Genet. Evol. 8(3): 239–246. https://doi.org/10.1016/j.meegid.2007.08.001

Patané JSL, Martins J, Jr., Rangel LT, Belasque J, Digiampietri LA, Facincani AP, Ferreira RM, Jaciani FJ, Zhang Y, Varani AM, Almeida NF, Wang N, Ferro JA, Moreira LM & Setubal JC (2019). Origin and diversification of *Xanthomonas citri* subsp. *citri* pathotypes revealed by inclusive phylogenomic, dating, and biogeographic analyses. BMC Genomics 20(1): 700. https://doi.org/10.1186/s12864-019-6007-4

Pruvost O, Magne M, Boyer K, Leduc A, Tourterel C, Drevet C, Ravigné V, Gagnevin L, Guérin F, Chiroleu F, Koebnik R, Verdier V & Vernière C (2014). A MLVA genotyping scheme for global surveillance of the citrus pathogen *Xanthomonas citri* pv. *citri* suggests a worldwide geographical expansion of a single genetic lineage. PLoS One 9(6): e98129. https://doi.org/10.1371/journal.pone.0098129

Quinlan AR & Hall IM (2010). BEDTools: a flexible suite of utilities for comparing genomic features. Bioinformatics 26(6): 841–842. https://doi.org/10.1093/bioinformatics/btq033

Rambaut A, Drummond AJ, Xie D, Baele G & Suchard MA (2018). Posterior summarization in Bayesian phylogenetics using Tracer 1.7. Syst. Biol. 67(5): 901–904. https://doi.org/10.1093/sysbio/syy032

Rasmussen DA & Grunwald NJ (2021). Phylogeographic approaches to characterize the emergence of plant pathogens. Phytopathology 111(1): 68–77. https://doi.org/10.1094/PHYTO-07-20-0319-FI

Rasmussen SO, Andersen KK, Svensson AM, Steffensen JP, Vinther BM, Clausen HB, Siggaard-Andersen ML, Johnsen SJ, Larsen LB, Dahl-Jensen D, Bigler M, Rothlisberger R, Fischer H, Goto-Azuma K, Hansson ME & Ruth U (2006). A new Greenland ice core chronology for the last glacial termination. J. Geophys. Res. Atmos. 111(D6). https://doi.org/10.1029/2005jd006079

Richard D, Boyer C, Javegny S, Boyer K, Grygiel P, Pruvost O, Rioualec AL, Rakotobe V, Iotti J, Picard R, Vernière C, Audusseau C, Francois C, Olivier V, Moreau A & Chabirand A (2016). First report of *Xanthomonas citri* pv. *citri* pathotype A causing Asiatic citrus canker in Martinique, France. Plant Dis. 100(9): 1946–1946. https://doi.org/10.1094/Pdis-02-16-0170-Pdn

Richard D, Pruvost O, Balloux F, Boyer C, Rieux A & Lefeuvre P (2021). Time-calibrated genomic evolution of a monomorphic bacterium during its establishment as an endemic crop pathogen. Mol. Ecol. 30(8): 1823–1835. https://doi.org/10.1111/mec.15770

Richard D, Roumagnac P, Pruvost O & Lefeuvre P (2022). A network approach to decipher the dynamics of Lysobacteraceae plasmid gene sharing. Mol. Ecol. (00): 1–14. https://doi.org/10.1111/mec.16536

Rieux A & Balloux F (2016a). Inferences from tip-calibrated phylogenies: a review and a practical guide. Mol. Ecol. 25(9): 1911–1924. https://doi.org/10.1111/mec.13586

Rieux A, Campos P, Duvermy A, Scussel S, Martin D, Gaudeul M, Lefeuvre P, Becker N & Lett JM (2021). Contribution of historical herbarium small RNAs to the reconstruction of a cassava mosaic geminivirus evolutionary history. Sci. Rep. 11(1): 21280. https://doi.org/10.1038/s41598-021-00518-w

Rieux A, Eriksson A, Li M, Sobkowiak B, Weinert LA, Warmuth V, Ruiz-Linares A, Manica A & Balloux F (2014). Improved calibration of the human mitochondrial clock using ancient genomes. Mol. Biol. Evol. 31(10): 2780–2792. https://doi.org/10.1093/molbev/msu222

Rieux A & Khatchikian CE (2016b). TIPDATINGBEAST: an R package to assist the implementation of phylogenetic tip-dating tests using BEAST. Mol. Ecol. Resour. 17: 608–613. https://doi.org/10.1111/1755-0998.12603

Ristaino JB (2020). The importance of mycological and plant herbaria in tracking plant killers. Front. Ecol. Evol. 7: 521. https://doi.org/10.3389/fevo.2019.00521

Rossetti V (1977). Citrus Canker in Latin America: a review. Proceedings of the International Society of Citriculture. 3:918–924.

Roux B, Bolot S, Guy E, Denancé N, Lautier M, Jardinaud MF, Fischer-Le Saux M, Portier P, Jacques MA, Gagnevin L, Pruvost O, Lauber E, Arlat M, Carrère S, Koebnik R & Noël LD (2015). Genomics and transcriptomics of *Xanthomonas campestris* species challenge the concept of core type III effectome. BMC Genomics 16(1): 975. https://doi.org/10.1186/s12864-015-2190-0

Rybak M, Minsavage GV, Stall RE & Jones JB (2009). Identification of *Xanthomonas citri* ssp. *citri* host specificity genes in a heterologous expression host. Mol. Plant Pathol. 10(2): 249–262. https://doi.org/10.1111/j.1364-3703.2008.00528.x

Savary S, Willocquet L, Pethybridge SJ, Esker P, McRoberts N & Nelson A (2019). The global burden of pathogens and pests on major food crops. Nat. Ecol. Evol. 3(3): 430–439. https://doi.org/10.1038/s41559-018-0793-y

Saville AC, Martin MD & Ristaino JB (2016). Historic late blight outbreaks caused by a widespread dominant lineage of *Phytophthora infestans* (Mont.) de Bary. PLoS One 11(12): e0168381. https://doi.org/10.1371/journal.pone.0168381

Schubert M, Lindgreen S & Orlando L (2016). AdapterRemoval v2: rapid adapter trimming, identification, and read merging. BMC Res. Notes 9: 88. https://doi.org/10.1186/s13104-016-1900-2

Schubert TS, Rizvi SA, Sun X, Gottwald TR, Graham JH & Dixon WN (2001). Meeting the challenge of eradicating Citrus Canker in Florida-again. Plant Dis. 85(4): 340–356. https://doi.org/10.1094/PDIS.2001.85.4.340

Smith O, Clapham A, Rose P, Liu Y, Wang J & Allaby RG (2014). A complete ancient RNA genome: identification, reconstruction and evolutionary history of archaeological Barley Stripe Mosaic Virus. Sci. Rep. 4: 4003. https://doi.org/10.1038/srep04003

Stamatakis A (2014). RAxML version 8: a tool for phylogenetic analysis and post-analysis of large phylogenies. Bioinformatics 30(9): 1312–1313. https://doi.org/10.1093/bioinformatics/btu033

Staubwasser M & Weiss H (2006). Holocene climate and cultural evolution in late prehistoric–early historic West Asia. Quat. Res 66(3): 372–387. https://doi.org/10.1016/j.yqres.2006.09.001

Stukenbrock EH & McDonald BA (2008). The origins of plant pathogens in agro-ecosystems. Annu. Rev. Phytopathol. 46(1): 75–100. https://doi.org/10.1146/annurev.phyto.010708.154114

Suchard MA & Rambaut A (2009). Many-core algorithms for statistical phylogenetics. Bioinformatics 25(11): 1370–1376. https://doi.org/10.1093/bioinformatics/btp244

Sun X, Stall RE, Jones JB, Cubero J, Gottwald TR, Graham JH, Dixon WN, Schubert TS, Chaloux PH, Stromberg VK, Lacy GH & Sutton BD (2004). Detection and characterization of a new strain of citrus canker bacteria from Key/Mexican lime and alemow in South Florida. Plant Dis. 88(11): 1179–1188. https://doi.org/10.1094/PDIS.2004.88.11.1179

Talon M, Caruso M & Gmitter Jr FG, Eds. (2020). “The genus Citrus”. Duxford, Elsevier. https://doi.org/10.1016/C2016-0-02375-6

Teper D, Burstein D, Salomon D, Gershovitz M, Pupko T & Sessa G (2016). Identification of novel *Xanthomonas euvesicatoria* type III effector proteins by a machine-learning approach. Mol. Plant Pathol. 17(3): 398–411. https://doi.org/10.1111/mpp.12288

The Xanthomonas Resource. Accessed December 2021, http://internet.myds.me/dokuwiki/doku.php?id=bacteria:t3e:t3e

Vernière C, Hartung JS, Pruvost OP, Civerolo EL, Alvarez AM, Maestri P & Luisetti J (1998). Characterization of phenotypically distinct strains of *Xanthomonas axonopodis* pv. *citri* from Southwest Asia. Eur. J. Plant Pathol. 104(5): 477–487. https://doi.org/10.1023/A:1008676508688

Walker M, Johnsen S, Rasmussen SO, Popp T, Steffensen JP, Gibbard P, Hoek W, Lowe J, Andrews J, Bjorck S, Cwynar LC, Hughen K, Kershaw P, Kromer B, Litt T, Lowe DJ, Nakagawa T, Newnham R & Schwander J (2009). Formal definition and dating of the GSSP (Global Stratotype Section and Point) for the base of the Holocene using the Greenland NGRIP ice core, and selected auxiliary records. J. Quaternary Sci. 24(1): 3–17. https://doi.org/10.1002/jqs.1227

Weiss CL, Schuenemann VJ, Devos J, Shirsekar G, Reiter E, Gould BA, Stinchcombe JR, Krause J & Burbano HA (2016). Temporal patterns of damage and decay kinetics of DNA retrieved from plant herbarium specimens. R. Soc. Open Sci. 3(6): 160239. https://doi.org/10.1098/rsos.160239

Wu GA, Terol J, Ibanez V, López-García A, Pérez-Román E, Borredá C, Domingo C, Tadeo FR, Carbonell-Caballero J, Alonso R, Curk F, Du D, Ollitrault P, Roose ML, Dopazo J, Gmitter FG, Rokhsar DS & Talon M (2018). Genomics of the origin and evolution of Citrus. Nature 554(7692): 311–316. https://doi.org/10.1038/nature25447

Yoshida K, Burbano HA, Krause J, Thines M, Weigel D & Kamoun S (2014). Mining herbaria for plant pathogen genomes: back to the future. PLoS Pathog. 10(4): e1004028. https://doi.org/10.1371/journal.ppat.1004028

Yoshida K, Sasaki E & Kamoun S (2015). Computational analyses of ancient pathogen DNA from herbarium samples: challenges and prospects. Front. Plant Sci. 6: 771. https://doi.org/10.3389/fpls.2015.00771

Yoshida K, Schuenemann VJ, Cano LM, Pais M, Mishra B, Sharma R, Lanz C, Martin FN, Kamoun S, Krause J, Thines M, Weigel D & Burbano HA (2013). The rise and fall of the *Phytophthora infestans* lineage that triggered the Irish potato famine. eLife 2: e00731. https://doi.org/10.7554/eLife.00731

Yu GC, Smith DK, Zhu HC, Guan Y & Lam TTY (2017). GGTREE: an R package for visualization and annotation of phylogenetic trees with their covariates and other associated data. Methods Ecol. Evol. 8(1): 28–36. https://doi.org/10.1111/2041-210x.12628

Zech-Matterne V & Fiorentino G (2017). “Agrumed: archaeology and history of citrus fruit in the mediterranean”. Napoli, Italy, Publications du Centre Jean Bérard. https://doi.org/10.4000/books.pcjb.2107

Zhang Y, Jalan N, Zhou X, Goss E, Jones JB, Setubal JC, Deng X & Wang N (2015). Positive selection is the main driving force for evolution of citrus canker-causing *Xanthomonas*. ISME J. 9(10): 2128–2138. https://doi.org/10.1038/ismej.2015.15

