## Supplementary files for "Herbarium Specimen Sequencing Allows Precise Datation of *Xanthomonas citri* pv. *citri* Diversification History"

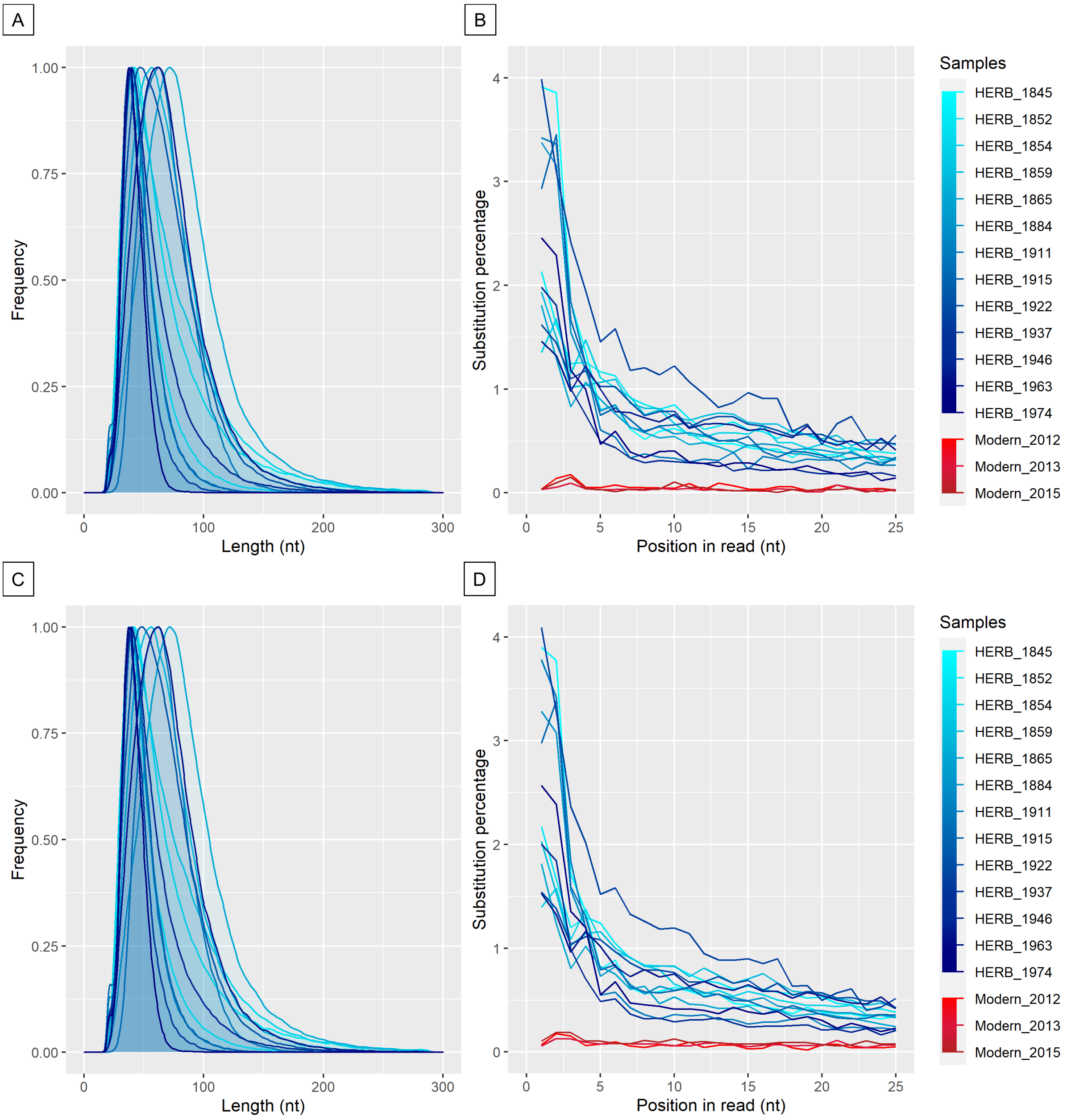


**S1 Fig. *Post-mortem* DNA damage patterns on reads mapping to *Xci* plasmids pXAC33 and pXAC64.** (A&C) Fragment length distribution (relative frequency in arbitrary units) of reads mapping to plasmids pXAC33 and pXAC64 of the reference strain, respectively. (B&D) G>A substitution percentage of the first 25 nucleotides from the 3' end of the 13 historical genomes (blue lines, light to dark gradient from the oldest to the youngest) and three modern *Xci* strains (red lines, light to dark gradient from the oldest to the youngest) for reads mapping to plasmids pXAC33 and pXAC64 of the reference strain, respectively.

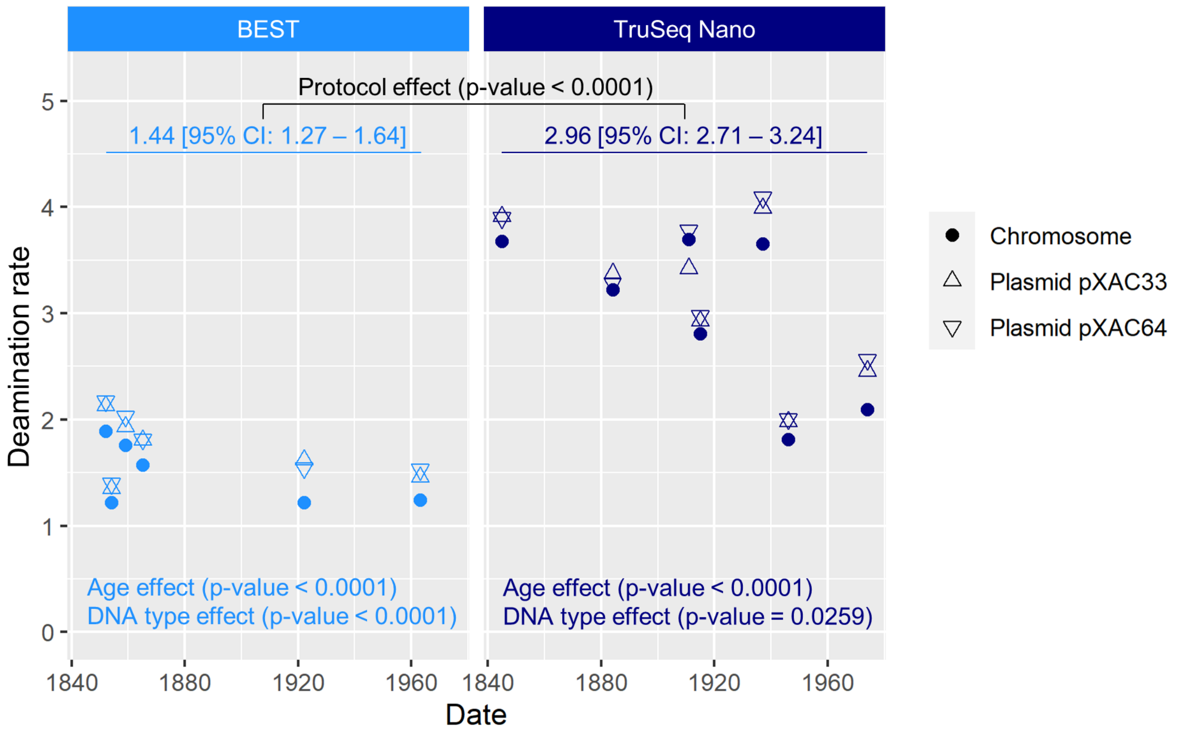


**S2 Fig. Deamination rate (in percentage) at terminal position from the 3' end (G to A substitutions) as a function of age of sample for the 13 *Xci* genomes reconstructed from herbarium specimens.** *Xci* sequencing reads (mapped to the chromosome, plasmids pXAC33 and pXAC64; circle, triangle and inverted triangle, respectively) were split into two datasets depending on the library protocol used (BEST or TruSeq Nano), according to a significant protocol effect on deamination rates. Within each dataset, a significant age effect was found. Furthermore, reads mapping to plasmid references were found significantly more deaminated than the chromosome DNA (DNA type effect).

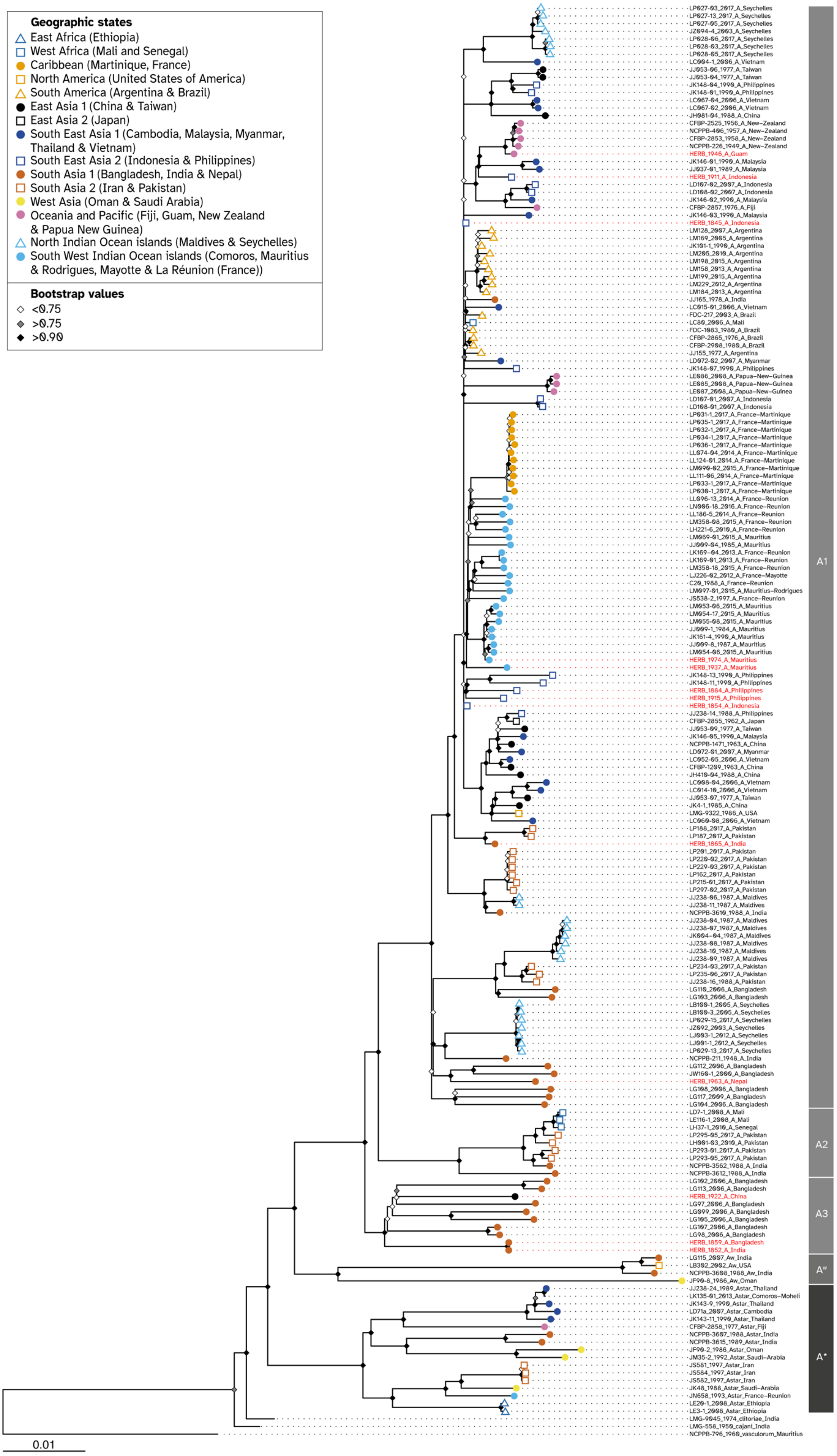
Geographic states
East Africa (Ethiopia)
West Africa (Mali and Senegal)
Caribbean (Martinique, France)
North America (United States of America)
South America (Argentina & Brazil)
East Asia 1 (China & Taiwan)
East Asia 2 (Japan)
South East Asia 1 (Cambodia, Malaysia, Myanmar, Thailand & Vietnam)
South East Asia 2 (Indonesia & Philippines)
South Asia 1 (Bangladesh, India & Nepal)
South Asia 2 (Iran & Pakistan)
West Asia (Oman & Saudi Arabia)
Oceania and Pacific (Fiji, Guam, New Zealand & Papua New Guinea)
North Indian Ocean islands (Maldives & Seychelles)
South West Indian Ocean islands (Comoros, Mauritius & Rodrigues, Mayotte & La Réunion (France))
Bootstrap values
<0.75
>0.75
>0.90
LP027-03_2017_A_Seychelles
LP027-13_2017_A_Seychelles
LP027-05_2017_A_Seychelles
JZ094-4_2003_A_Seychelles
LP028-06_2017_A_Seychelles
LP028-03_2017_A_Seychelles
LP028-05_2017_A_Seychelles
LC004-1_2006_A_Vietnam
JJ053-06_1977_A_Taiwan
JJ053-04_1977_A_Taiwan
JK148-04_1990_A_Philippines
JK148-01_1990_A_Philippines
LC067-04_2006_A_Vietnam
LC067-02_2006_A_Vietnam
JH081-04_1988_A_China
CFBP-2525_1956_A_New-Zealand
NCPPB-406_1957_A_New-Zealand
CFBP-2853_1958_A_New-Zealand
NCPPB-226_1949_A_New-Zealand
HERB_1946_A_Guam
JK146-01_1990_A_Malaysia
JJ037-01_1989_A_Malaysia
HERB_1911_A_Indonesia
LD107-02_2007_A_Indonesia
LD108-02_2007_A_Indonesia
JK146-02_1990_A_Malaysia
CFBP-2857_1976_A_Fiji
JK146-03_1990_A_Malaysia
HERB_1845_A_Indonesia
LM128_2007_A_Argentina
LM169_2005_A_Argentina
JK101-1_1990_A_Argentina
LM205_2010_A_Argentina
LM198_2015_A_Argentina
LM158_2013_A_Argentina
LM199_2015_A_Argentina
LM229_2012_A_Argentina
LM184_2013_A_Argentina
JJ165_1978_A_India
LC015-01_2006_A_Vietnam
FDC-217_2003_A_Brazil
LC80_2006_A_Mali
FDC-1083_1980_A_Brazil
CFBP-2865_1976_A_Brazil
CFBP-2908_1980_A_Brazil
JJ155_1977_A_Argentina
LD072-02_2007_A_Myanmar
JK148-07_1990_A_Philippines
LE086_2008_A_Papua-New-Guinea
LE085_2008_A_Papua-New-Guinea
LE087_2008_A_Papua-New-Guinea
LD107-01_2007_A_Indonesia
LD108-01_2007_A_Indonesia
LP031-1_2017_A_France-Martinique
LP035-1_2017_A_France-Martinique
LP032-1_2017_A_France-Martinique
LP034-1_2017_A_France-Martinique
LP036-1_2017_A_France-Martinique
LL074-04_2014_A_France-Martinique
LL124-01_2014_A_France-Martinique
LM090-02_2015_A_France-Martinique
LL111-06_2014_A_France-Martinique
LP033-1_2017_A_France-Martinique
LP030-1_2017_A_France-Martinique
LL096-13_2014_A_France-Reunion
LN006-18_2016_A_France-Reunion
LL186-5_2014_A_France-Reunion
LM358-08_2015_A_France-Reunion
LH221-6_2010_A_France-Reunion
LM069-01_2015_A_Mauritius
JJ009-04_1985_A_Mauritius
LK169-04_2013_A_France-Reunion
LK169-01_2013_A_France-Reunion
LM358-18_2015_A_France-Reunion
LJ226-02_2012_A_France-Mayotte
C20_1988_A_France-Reunion
LM097-01_2015_A_Mauritius-Rodrigues
JS538-2_1997_A_France-Reunion
LM053-06_2015_A_Mauritius
LM054-17_2015_A_Mauritius
LM055-08_2015_A_Mauritius
JJ009-1_1984_A_Mauritius
JK161-4_1990_A_Mauritius
JJ009-8_1987_A_Mauritius
LM054-06_2015_A_Mauritius
HERB_1974_A_Mauritius
HERB_1937_A_Mauritius
JK148-13_1990_A_Philippines
JK148-11_1990_A_Philippines
HERB_1884_A_Philippines
HERB_1915_A_Philippines
HERB_1854_A_Indonesia
JJ238-14_1988_A_Philippines
CFBP-2855_1962_A_Japan
JJ053-09_1977_A_Taiwan
JK146-05_1990_A_Malaysia
NCPPB-1471_1963_A_China
LD072-01_2007_A_Myanmar
LC052-05_2006_A_Vietnam
CFBP-1209_1963_A_China
JH410-04_1988_A_China
LC008-04_2006_A_Vietnam
LC014-10_2006_A_Vietnam
JJ053-07_1977_A_Taiwan
JK4-1_1985_A_China
LMG-9322_1986_A_USA
LC060-08_2006_A_Vietnam
LP188_2017_A_Pakistan
LP187_2017_A_Pakistan
HERB_1865_A_India
LP201_2017_A_Pakistan
LP220-02_2017_A_Pakistan
LP229-03_2017_A_Pakistan
LP162_2017_A_Pakistan
LP215-01_2017_A_Pakistan
LP297-02_2017_A_Pakistan
JJ238-06_1987_A_Maldives
JJ238-11_1987_A_Maldives
NCPPB-3610_1988_A_India
JJ238-04_1987_A_Maldives
JJ238-07_1987_A_Maldives
JK004-04_1987_A_Maldives
JJ238-08_1987_A_Maldives
JJ238-10_1987_A_Maldives
JJ238-09_1987_A_Maldives
LP234-03_2017_A_Pakistan
LP235-06_2017_A_Pakistan
JJ238-16_1988_A_Pakistan
LG110_2006_A_Bangladesh
LG103_2006_A_Bangladesh
LB100-1_2005_A_Seychelles
LB100-3_2005_A_Seychelles
LP029-15_2017_A_Seychelles
JZ092_2003_A_Seychelles
LJ003-1_2012_A_Seychelles
LJ001-1_2012_A_Seychelles
LP029-13_2017_A_Seychelles
NCPPB-211_1948_A_India
LG112_2006_A_Bangladesh
JW160-1_2000_A_Bangladesh
HERB_1963_A_Nepal
LG108_2006_A_Bangladesh
LG117_2009_A_Bangladesh
LG104_2006_A_Bangladesh
LD7-1_2008_A_Mali
LE116-1_2008_A_Mali
LH37-1_2010_A_Senegal
LP295-05_2017_A_Pakistan
LH001-03_2010_A_Pakistan
LP293-01_2017_A_Pakistan
LP293-05_2017_A_Pakistan
NCPPB-3562_1988_A_India
NCPPB-3612_1988_A_India
LG102_2006_A_Bangladesh
LG113_2006_A_Bangladesh
HERB_1922_A_China
LG97_2006_A_Bangladesh
LG099_2006_A_Bangladesh
LG105_2006_A_Bangladesh
LG107_2006_A_Bangladesh
LG98_2006_A_Bangladesh
HERB_1859_A_Bangladesh
HERB_1852_A_India
LG115_2007_Aw_India
LB302_2002_Aw_USA
NCPPB-3608_1988_Aw_India
JF90-8_1986_Aw_Oman
JJ238-24_1989_Astar_Thailand
LK135-01_2013_Astar_Comoros-Moheli
JK143-9_1990_Astar_Thailand
LD71a_2007_Astar_Cambodia
JK143-11_1990_Astar_Thailand
CFBP-2858_1977_Astar_Fiji
NCPPB-3607_1988_Astar_India
NCPPB-3615_1989_Astar_India
JF90-2_1986_Astar_Oman
JM35-2_1992_Astar_Saudi-Arabia
JS581_1997_Astar_Iran
JS584_1997_Astar_Iran
JS582_1997_Astar_Iran
JK48_1988_Astar_Saudi-Arabia
JN658_1993_Astar_France-Reunion
LE20-1_2008_Astar_Ethiopia
LE3-1_2008_Astar_Ethiopia
LMG-9045_1974_clitoriae_India
LMG-558_1950_cajani_India
NCPPB-796_1960_vasculorum_Mauritius
A1
A2
A3
Aw
A*
0.01

**S3 Fig. Maximum Likelihood (ML) phylogenetic tree of historical and modern *Xci* genomes.** ML tree including 13 historical specimens (labelled in red) and 171 modern strains (black) built from 13,007 recombination-free SNPs. *Xanthomonas axonopodis* pv. *vasculorum* NCPPB-796 isolated in 1960 from Mauritius (GenBank accession number: GCF_013177355.1) was used to root the tree. Node values correspond to bootstrap values calculated on 1,000 iterations. Strain labels include strain name, collection year, pathotype and country of origin; branch tips are colored according to the sample's geographic origin (see Materials and Methods for details). Pathotypes and lineages are indicated to the right.

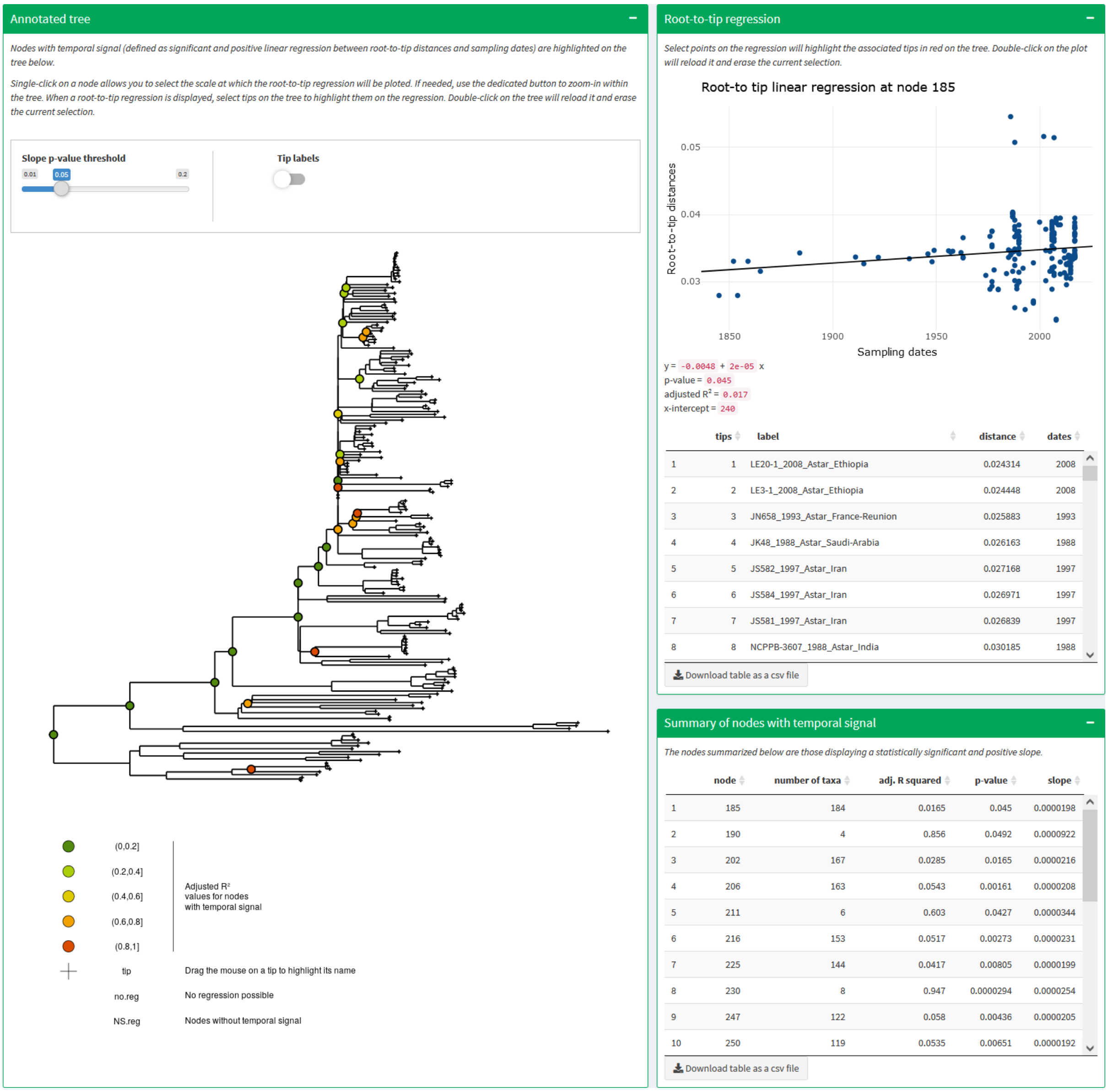


**S4 Fig. Root-to-tip regression temporal test visualized on online tool PhyloStemS.** Nodes with temporal signal, *i.e.* nodes at which root-to-tip linear regression yielded a statistically significant and positive slope, are highlighted with colors scaling to R2 value. The tool is accessible online (https://pvbmt-apps.cirad.fr/apps/phylostems/).

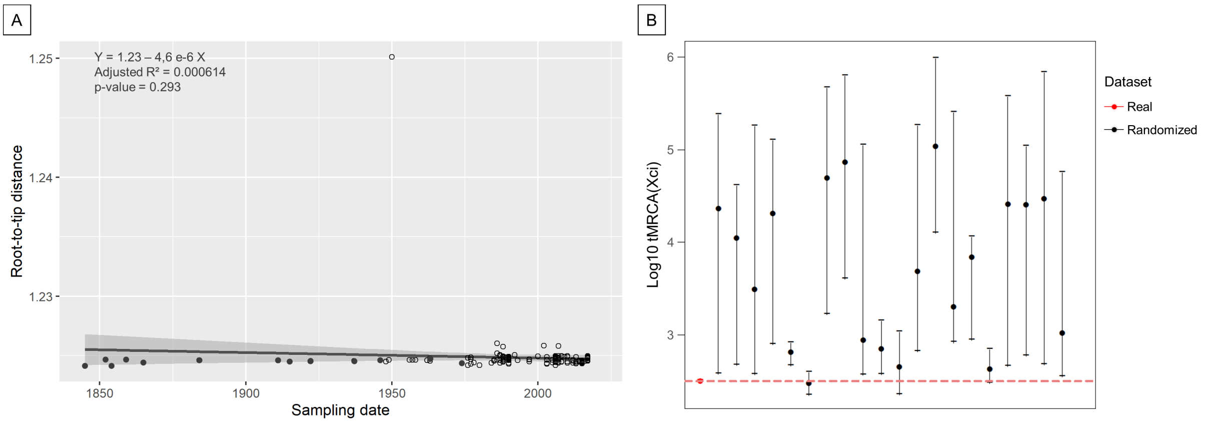


**S5 Fig. Root-to-tip regression and date-randomization temporal test results when performed on the dataset including outgroups**. (A) Regression line is plotted in black with historical genomes in black filled dots and modern ones in black empty dots. Grey areas indicate the confidence interval. Associated values are the regression equation, adjusted $R^2$ (Adj $R^2$) and p-value. (B) Evaluating temporal signal in the dataset by date-randomization test showed overlap between the age of the root estimated from the real dataset (red) vs 20 date-randomized datasets (black). Vertical bars represent 95% Highest Posterior Density intervals.

**S1 Table. General characteristics of the historical and modern strains of the study.** Labels consist in strain ID, year of collection, *Xci* pathotype (or other *Xanthomonas* pathovar: vasculorum= *Xanthomonas axonopodis* pv. *vasculorum*; clitoriae= *Xanthomonas citri* pv. *clitoriae*; cajani= *Xanthomonas citri* pv. *cajani*) and country of origin. Geographical origin areas are described in the Materials and Methods.

| **Label[1]** | **Source material** | **Geographic origin** | **Isolation Host** | **GenBank accession ID** | **Sequence Read Archive accession ID** | **Publication** | **Provider[2]** |
|---|---|---|---|---|---|---|---|
| HERB_1845_A_Indonesia | Herbarium tissue | South East Asia 2 | *Citrus aurantiifolia* | CP106931 | SRR21711661 | This study | MNHN, France[3] |
| HERB_1852_A_India | Herbarium tissue | South Asia 1 | *Citrus medica* | CP106932 | SRR21721660 | This study | Royal Botanic Gardens, UK[3] |
| HERB_1854_A_Indonesia | Herbarium tissue | South East Asia 2 | *Citrus javanica* | CP106933 | SRR21721657 | This study | Royal Botanic Gardens, UK[3] |
| HERB_1859_A_Bangladesh | Herbarium tissue | South Asia 1 | *Citrus medica* | CP106934 | SRR21721656 | This study | MNHN, France[3] |
| HERB_1865_A_India | Herbarium tissue | South Asia 1 | *Citrus medica* | CP106935 | SRR21721655 | This study | Royal Botanic Gardens, UK[3] |
| HERB_1884_A_Philippines | Herbarium tissue | South East Asia 2 | *Citrus medica* | CP106936 | SRR21721654 | This study | Royal Botanic Gardens, UK[3] |
| HERB_1911_A_Indonesia | Herbarium tissue | South East Asia 2 | *Citrus aurantiifolia* | CP106937 | SRR21721653 | This study | MNHN, France[3] |
| HERB_1915_A_Philippines | Herbarium tissue | South East Asia 2 | *Citrus aurantiifolia* | CP106938 | SRR21721652 | This study | MNHN, France[3] |
| HERB_1922_A_China | Herbarium tissue | East Asia 1 | *Citrus medica* | CP106939 | SRR21721651 | This study | U.S. National Herbarium[3] |
| HERB_1937_A_Mauritius | Herbarium tissue | South West Indian Ocean islands | *Citrus* sp. | CP072205-CP072207 | SRR12792042 | Campos *et al.*, 2021 | NA |
| HERB_1946_A_Guam | Herbarium tissue | Oceania & Pacific | *Citrus* sp. | CP106940 | SRR21721650 | This study | US Ntnl Fungus Collections[3] |
| HERB_1963_A_Nepal | Herbarium tissue | South Asia 1 | *Citrus medica* | CP106941 | SRR21721659 | This study | Royal Botanic Gardens, UK[3] |
| HERB_1974_A_Mauritius | Herbarium tissue | South West Indian Ocean islands | *Citrus* “lime” | CP106942 | SRR21721658 | This study | Mauritius Herbarium[3] |
| C20_1988_A_France-Reunion | Bacterial culture | South West Indian Ocean islands | *Citrus paradisi* | JAABHN000000000 | SRR11234852 | Richard *et al.*, 2021 | NA |
| CFBP-1209_1963_A_China-Hong-Kong | Bacterial culture | East Asia 1 | *Citrus maxima* | JAABAP000000000 | SRR11234741 | Richard *et al.*, 2021 | NA |
| CFBP-2525_1956_A_New-Zealand | Bacterial culture | Oceania & Pacific | *Citrus limon* | JAABCU000000000 | SRR11234614 | Richard *et al.*, 2021 | NA |
| CFBP-2853_1958_A_New-Zealand | Bacterial culture | Oceania & Pacific | *Citrus* sp. | JAABCV000000000 | SRR11234592 | Richard *et al.*, 2021 | NA |
| CFBP-2855_1962_A_Japan | Bacterial culture | East Asia 2 | *Citrus* sp. | JAABAW000000000 | SRR11234581 | Richard *et al.*, 2021 | NA |
| CFBP-2857_1976_A_Fiji | Bacterial culture | Oceania & Pacific | *Citrus aurantiifolia* | JAABAO000000000 | SRR11234570 | Richard *et al.*, 2021 | NA |
| CFBP-2865_1976_A_Brazil | Bacterial culture | South America | *Citrus aurantiifolia* | JAAAYX000000000 | SRR11234559 | Richard *et al.*, 2021 | NA |
| CFBP-2908_1980_A_Brazil | Bacterial culture | South America | *Citrus reticulata* | JAAAYY000000000 | SRR11234851 | Richard *et al.*, 2021 | NA |
| FDC-1083_1980_A_Brazil | Bacterial culture | South America | *Citrus reticulata* | CCVZ00000000.1 | SRR11234818 | Gordon *et al.*, 2015 | NA |
| FDC-217_2003_A_Brazil | Bacterial culture | South America | *Citrus sinensis* | CCWY00000000.1 | SRR11234807 | Gordon *et al.*, 2015 | NA |
| JH081-04_1988_A_China | Bacterial culture | East Asia 1 | *Citrus* sp. | | SRR21684687 | This study | FAO |
| JH410-04_1988_A_China | Bacterial culture | East Asia 1 | *Citrus reticulata* | | SRR21684686 | This study | FAO |
| JJ009-04_1985_A_Mauritius | Bacterial culture | South West Indian Ocean islands | *Citrus* sp. x *bergamia* | JAABBT000000000 | SRR11234774 | Richard *et al.*, 2021 | NA |
| JJ009-1_1984_A_Mauritius | Bacterial culture | South West Indian Ocean islands | *Citrus sinensis* x *Poncirus* trifoliata | JAABBU000000000 | SRR11234763 | Richard *et al.*, 2021 | NA |
| JJ009-8_1987_A_Mauritius | Bacterial culture | South West Indian Ocean islands | *Citrus sinensis* | JAABBV000000000 | SRR11234752 | Richard *et al.*, 2021 | NA |
| JJ037-01_1989_A_Malaysia | Bacterial culture | South East Asia 1 | *Citrus sinensis* | | SRR21684675 | This study | FAO |
| JJ053-04_1977_A_Taiwan | Bacterial culture | East Asia 1 | *Poncirus trifoliata* | | SRR21684664 | This study | NCHU |
| JJ053-06_1977_A_Taiwan | Bacterial culture | East Asia 1 | *Citrus* “sunki” | JAABJE000000000 | SRR11234718 | Richard *et al.*, 2021 | NA |
| JJ053-07_1977_A_Taiwan | Bacterial culture | East Asia 1 | *Citrus* “limonia” | | SRR21684653 | This study | NCHU |
| JJ053-09_1977_A_Taiwan | Bacterial culture | East Asia 1 | *Citrus aurantiifolia* | | SRR21684642 | This study | NCHU |
| JJ155_1977_A_Argentina | Bacterial culture | South America | *Citrus aurantiifolia* | JAAAYI000000000 | SRR11234707 | Richard *et al.*, 2021 | NA |
| JJ165_1978_A_India | Bacterial culture | South Asia 1 | *Citrus maxima* | JAABAR000000000 | SRR11234696 | Richard *et al.*, 2021 | NA |
| JJ238-04_1987_A_Maldives | Bacterial culture | North Indian Ocean islands | *Citrus aurantiifolia* | JAABAX000000000 | SRR11234685 | Richard *et al.*, 2021 | NA |

| Label[1] | Source material | Geographic origin | Isolation Host | GenBank accession ID | Sequence Read Archive accession ID | Publication | Provider[2] |
|---|---|---|---|---|---|---|---|
| JJ238-06_1987_A_Maldives | Bacterial culture | North Indian Ocean islands | *Citrus aurantiifolia* | JAABAY000000000 | SRR11234674 | Richard *et al.*, 2021 | NA |
| JJ238-07_1987_A_Maldives | Bacterial culture | North Indian Ocean islands | *Citrus aurantiifolia* | JAABAZ000000000 | SRR11234663 | Richard *et al.*, 2021 | NA |
| JJ238-08_1987_A_Maldives | Bacterial culture | North Indian Ocean islands | *Citrus aurantiifolia* | JAABBA000000000 | SRR11234652 | Richard *et al.*, 2021 | NA |
| JJ238-09_1987_A_Maldives | Bacterial culture | North Indian Ocean islands | *Citrus aurantiifolia* | JAABBB000000000 | SRR11234641 | Richard *et al.*, 2021 | NA |
| JJ238-10_1987_A_Maldives | Bacterial culture | North Indian Ocean islands | *Citrus aurantiifolia* | CCWC00000000.1 | SRR11234629 | Gordon *et al.*, 2015 | NA |
| JJ238-11_1987_A_Maldives | Bacterial culture | North Indian Ocean islands | *Citrus aurantiifolia* | JAABBD000000000 | SRR11234623 | Richard *et al.*, 2021 | NA |
| JJ238-14_1988_A_Philippines | Bacterial culture | South East Asia 2 | *Citrus* sp. | | SRR21684631 | This study | FAO |
| JJ238-16_1988_A_Pakistan | Bacterial culture | South Asia 2 | *Citrus sinensis* | JAABCY000000000 | SRR11234622 | Richard *et al.*, 2021 | NA |
| JK004-04_1987_A_Maldives | Bacterial culture | North Indian Ocean islands | *Citrus aurantiifolia* | JAABBE000000000 | SRR11234621 | Richard *et al.*, 2021 | NA |
| JK101-1_1990_A_Argentina | Bacterial culture | South America | *Citrus paradisi* | JAAAYJ000000000 | SRR11234620 | Richard *et al.*, 2021 | NA |
| JK146-01_1990_A_Malaysia | Bacterial culture | South East Asia 1 | *Citrus maxima* | | SRR21684630 | This study | FAO |
| JK146-02_1990_A_Malaysia | Bacterial culture | South East Asia 1 | *Citrus reticulata* | | SRR21684629 | This study | FAO |
| JK146-03_1990_A_Malaysia | Bacterial culture | South East Asia 1 | *Citrus reticulata* | | SRR21684628 | This study | FAO |
| JK146-05_1990_A_Malaysia | Bacterial culture | South East Asia 1 | *Citrus maxima* | | SRR21684685 | This study | FAO |
| JK148-01_1990_A_Philippines | Bacterial culture | South East Asia 2 | *Citrus aurantiifolia* | | SRR21684684 | This study | FAO |
| JK148-04_1990_A_Philippines | Bacterial culture | South East Asia 2 | *Citrus limon* | | SRR21684683 | This study | FAO |
| JK148-07_1990_A_Philippines | Bacterial culture | South East Asia 2 | "Citrofortunella microcarpa" | | SRR21684682 | This study | FAO |
| JK148-11_1990_A_Philippines | Bacterial culture | South East Asia 2 | *Citrus maxima* | | SRR21684681 | This study | FAO |
| JK148-13_1990_A_Philippines | Bacterial culture | South East Asia 2 | *Citrus maxima* | | SRR21684680 | This study | FAO |
| JK161-4_1990_A_Mauritius | Bacterial culture | South West Indian Ocean islands | *Citrus sinensis* | JAABBW000000000 | SRR11234619 | Richard *et al.*, 2021 | NA |
| JK4-1_1985_A_China | Bacterial culture | East Asia 1 | *Citrus* sp. | CDMR01000000 | SRR11234656 | Gordon *et al.*, 2015 | NA |
| JS538-2_1997_A_France-Reunion | Bacterial culture | South West Indian Ocean islands | *Citrus paradisi* | JAABHE000000000 | SRR11234598 | Richard *et al.*, 2021 | NA |
| JW160-1_2000_A_Bangladesh | Bacterial culture | South Asia 1 | *Citrus aurantiifolia* | CCWH00000000 | | Gordon *et al.*, 2015 | NA |
| JZ092_2003_A_Seychelles | Bacterial culture | North Indian Ocean islands | *Citrus limon* | JAABIQ000000000 | SRR11234595 | Richard *et al.*, 2021 | NA |
| JZ094-4_2003_A_Seychelles | Bacterial culture | North Indian Ocean islands | *Citrus limon* | JAABIR000000000 | SRR11234594 | Richard *et al.*, 2021 | NA |
| LB100-1_2005_A_Seychelles | Bacterial culture | North Indian Ocean islands | *Citrus sinensis* x *Poncirus trifoliata* | CDAV01000000 | SRR11234654 | Gordon *et al.*, 2015 | NA |
| LB100-3_2005_A_Seychelles | Bacterial culture | North Indian Ocean islands | *Citrus sinensis* x *Poncirus trifoliata* | JAABIT000000000 | SRR11234591 | Richard *et al.*, 2021 | NA |
| LC004-1_2006_A_Vietnam | Bacterial culture | South East Asia 1 | *Citrus maxima* | JAABJG000000000 | SRR11234590 | Richard *et al.*, 2021 | NA |
| LC008-04_2006_A_Vietnam | Bacterial culture | South East Asia 1 | *Citrus reticulata* | | SRR21684679 | This study | SOFRI |
| LC014-10_2006_A_Vietnam | Bacterial culture | South East Asia 1 | *Citrus reticulata* | | SRR21684678 | This study | SOFRI |
| LC015-01_2006_A_Vietnam | Bacterial culture | South East Asia 1 | *Citrus* sp. | | SRR21684677 | This study | SOFRI |
| LC052-05_2006_A_Vietnam | Bacterial culture | South East Asia 1 | *Citrus sinensis* | | SRR21684676 | This study | PPRI |
| LC060-08_2006_A_Vietnam | Bacterial culture | South East Asia 1 | *Citrus sinensis* | | SRR21684674 | This study | PPRI |
| LC067-02_2006_A_Vietnam | Bacterial culture | South East Asia 1 | *Citrus sinensis* | | SRR21684673 | This study | PPRI |
| LC067-04_2006_A_Vietnam | Bacterial culture | South East Asia 1 | *Citrus maxima* | | SRR21684672 | This study | PPRI |
| LC80_2006_A_Mali | Bacterial culture | West Africa | *Citrus reticulata* x *Citrus sinensis* | CCWJ00000000.1 | SRR11234586 | Gordon *et al.*, 2015 | NA |
| LD072-01_2007_A_Myanmar | Bacterial culture | South East Asia 1 | *Citrus maxima* | | SRR21684671 | This study | H4D |
| LD072-02_2007_A_Myanmar | Bacterial culture | South East Asia 1 | *Citrus sinensis* | | SRR21684670 | This study | H4D |
| LD107-01_2007_A_Indonesia | Bacterial culture | South East Asia 2 | *Citrus maxima* | | SRR21684669 | This study | H4D |
| LD107-02_2007_A_Indonesia | Bacterial culture | South East Asia 2 | *Citrus maxima* | | SRR21684668 | This study | H4D |

| Label[1] | Source material | Geographic origin | Isolation Host | GenBank accession ID | Sequence Read Archive accession ID | Publication | Provider[2] |
|---|---|---|---|---|---|---|---|
| LD108-01_2007_A_Indonesia | Bacterial culture | South East Asia 2 | *Citrus aurantiifolia* | | SRR21684667 | This study | H4D |
| LD108-02_2007_A_Indonesia | Bacterial culture | South East Asia 2 | *Citrus aurantiifolia* | | SRR21684666 | This study | H4D |
| LD7-1_2008_A_Mali | Bacterial culture | West Africa | *Citrus aurantiifolia* | CDAL00000000.1 | SRR11234651 | Gordon *et al.*, 2015 | NA |
| LE085_2008_A_Papua-New-Guinea | Bacterial culture | Oceania & Pacific | *Citrus limon* | | SRR21684665 | This study | NAQS |
| LE086_2008_A_Papua-New-Guinea | Bacterial culture | Oceania & Pacific | *Citrus limon* | | SRR21684663 | This study | NAQS |
| LE087_2008_A_Papua-New-Guinea | Bacterial culture | Oceania & Pacific | *Citrus aurantiifolia* | | SRR21684662 | This study | NAQS |
| LE116-1_2008_A_Mali | Bacterial culture | West Africa | *Citrus aurantiifolia* | CDHD01000000 | SRR11234650 | Gordon *et al.*, 2015 | NA |
| LG099_2006_A_Bangladesh | Bacterial culture | South Asia 1 | *Citrus aurantiifolia* | | SRR21684661 | This study | FERA |
| LG102_2006_A_Bangladesh | Bacterial culture | South Asia 1 | *Citrus* sp. | CDAN01000000 | SRR11234649 | Gordon *et al.*, 2015 | NA |
| LG103_2006_A_Bangladesh | Bacterial culture | South Asia 1 | *Citrus aurantiifolia* | | SRR21684660 | This study | FERA |
| LG104_2006_A_Bangladesh | Bacterial culture | South Asia 1 | *Citrus aurantiifolia* | | SRR21684659 | This study | FERA |
| LG105_2006_A_Bangladesh | Bacterial culture | South Asia 1 | *Citrus aurantiifolia* | | SRR21684658 | This study | FERA |
| LG107_2006_A_Bangladesh | Bacterial culture | South Asia 1 | *Citrus aurantiifolia* | | SRR21684657 | This study | FERA |
| LG108_2006_A_Bangladesh | Bacterial culture | South Asia 1 | *Citrus aurantiifolia* | | SRR21684656 | This study | FERA |
| LG110_2006_A_Bangladesh | Bacterial culture | South Asia 1 | *Citrus aurantiifolia* | | SRR21684655 | This study | FERA |
| LG112_2006_A_Bangladesh | Bacterial culture | South Asia 1 | *Citrus aurantiifolia* | | SRR21684654 | This study | FERA |
| LG113_2006_A_Bangladesh | Bacterial culture | South Asia 1 | *Citrus aurantiifolia* | | SRR21684652 | This study | FERA |
| LG117_2009_A_Bangladesh | Bacterial culture | South Asia 1 | *Citrus* sp. | CDAX01000000 | SRR11234648 | Gordon *et al.*, 2015 | NA |
| LG97_2006_A_Bangladesh | Bacterial culture | South Asia 1 | *Citrus* sp. | CDAK01000000 | SRR11234647 | Gordon *et al.*, 2015 | NA |
| LG98_2006_A_Bangladesh | Bacterial culture | South Asia 1 | *Citrus aurantiifolia* | CDBA01000000 | SRR11234646 | Gordon *et al.*, 2015 | NA |
| LH001-03_2010_A_Pakistan | Bacterial culture | South Asia 2 | *Citrus* sp. | | SRR21684651 | This study | NARC |
| LH221-6_2010_A_France-Reunion | Bacterial culture | South West Indian Ocean islands | *Citrus hystrix* | JAABEL000000000 | SRR11234582 | Richard *et al.*, 2021 | NA |
| LH37-1_2010_A_Senegal | Bacterial culture | West Africa | *Citrus paradisi* | CDAS00000000.1 | SRR11234645 | Gordon *et al.*, 2015 | NA |
| LJ001-1_2012_A_Seychelles | Bacterial culture | North Indian Ocean islands | *Citrus* sp. | JAABIU000000000 | SRR11234557 | Richard *et al.*, 2021 | NA |
| LJ003-1_2012_A_Seychelles | Bacterial culture | North Indian Ocean islands | *Citrus sinensis* | JAABIV000000000 | SRR11234556 | Richard *et al.*, 2021 | NA |
| LJ226-02_2012_A_France-Mayotte | Bacterial culture | South West Indian Ocean islands | *Citrus sinensis* | JAABCN000000000 | SRR11234551 | Richard *et al.*, 2021 | NA |
| LK169-01_2013_A_France-Reunion | Bacterial culture | South West Indian Ocean islands | *Citrus hystrix* | JAABFW000000000 | SRR11234810 | Richard *et al.*, 2021 | NA |
| LK169-04_2013_A_France-Reunion | Bacterial culture | South West Indian Ocean islands | *Citrus hystrix* | JAABFY000000000 | SRR11234808 | Richard *et al.*, 2021 | NA |
| LL074-04_2014_A_France-Martinique | Bacterial culture | Central America & Caribbean | *Citrus paradisi* | JAABBI000000000 | SRR11234793 | Richard *et al.*, 2021 | NA |
| LL096-13_2014_A_France-Reunion | Bacterial culture | South West Indian Ocean islands | *Citrus limon* | JAABHJ000000000 | SRR11234772 | Richard *et al.*, 2021 | NA |
| LL111-06_2014_A_France-Martinique | Bacterial culture | Central America & Caribbean | *Citrus sinensis* | JAABBJ000000000 | SRR11234768 | Richard *et al.*, 2021 | NA |
| LL124-01_2014_A_France-Martinique | Bacterial culture | Central America & Caribbean | *Citrus sinensis* | JAABBK000000000 | SRR11234764 | Richard *et al.*, 2021 | NA |
| LL186-5_2014_A_France-Reunion | Bacterial culture | South West Indian Ocean islands | *Citrus sinensis* | JAABHV000000000 | SRR11234753 | Richard *et al.*, 2021 | NA |
| LM053-06_2015_A_Mauritius | Bacterial culture | South West Indian Ocean islands | *Citrus aurantiifolia* | JAABCA000000000 | SRR11234751 | Richard *et al.*, 2021 | NA |
| LM054-06_2015_A_Mauritius | Bacterial culture | South West Indian Ocean islands | *Citrus sinensis* | JAABCC000000000 | SRR11234749 | Richard *et al.*, 2021 | NA |
| LM054-17_2015_A_Mauritius | Bacterial culture | South West Indian Ocean islands | *Citrus sinensis* | JAABCD000000000 | SRR11234748 | Richard *et al.*, 2021 | NA |
| LM055-08_2015_A_Mauritius | Bacterial culture | South West Indian Ocean islands | *Citrus* sp. | JAABCE000000000 | SRR11234747 | Richard *et al.*, 2021 | NA |
| LM069-01_2015_A_Mauritius | Bacterial culture | South West Indian Ocean islands | *Citrus* aurantiifolia | JAABCI000000000 | SRR11234743 | Richard *et al.*, 2021 | NA |
| LM090-02_2015_A_France-Martinique | Bacterial culture | Central America & Caribbean | *Citrus* sp. | JAABBL000000000 | SRR11234726 | Richard *et al.*, 2021 | NA |
| LM097-01_2015_A_Mauritius-Rodrigues | Bacterial culture | South West Indian Ocean islands | *Citrus aurantiifolia* | JAABIN000000000 | SRR11234716 | Richard *et al.*, 2021 | NA |

| Label[1] | Source material | Geographic origin | Isolation Host | GenBank accession ID | Sequence Read Archive accession ID | Publication | Provider[2] |
|---|---|---|---|---|---|---|---|
| LM128_2007_A_Argentina | Bacterial culture | South America | *Citrus reticulata* | | SRR21684650 | This study | INTA |
| LM158_2013_A_Argentina | Bacterial culture | South America | *Citrus sinensis* | JAAAYK000000000 | SRR11234710 | Richard *et al.*, 2021 | NA |
| LM169_2005_A_Argentina | Bacterial culture | South America | *Citrus paradisi* | JAAAYL000000000 | SRR11234709 | Richard *et al.*, 2021 | NA |
| LM184_2013_A_Argentina | Bacterial culture | South America | *Citrus limon* | JAAAYN000000000 | SRR11234706 | Richard *et al.*, 2021 | NA |
| LM198_2015_A_Argentina | Bacterial culture | South America | *Citrus sinensis* | JAAAYO000000000 | SRR11234705 | Richard *et al.*, 2021 | NA |
| LM199_2015_A_Argentina | Bacterial culture | South America | *Citrus sinensis* | JAAAYP000000000 | SRR11234704 | Richard *et al.*, 2021 | NA |
| LM205_2010_A_Argentina | Bacterial culture | South America | *Citrus paradisi* | JAAAYQ000000000 | SRR11234703 | Richard *et al.*, 2021 | NA |
| LM229_2012_A_Argentina | Bacterial culture | South America | *Citrus sinensis* | JAAAYR000000000 | SRR11234702 | Richard *et al.*, 2021 | NA |
| LM358-08_2015_A_France-Reunion | Bacterial culture | South West Indian Ocean islands | *Citrus limon* | JAABFN000000000 | SRR11234700 | Richard *et al.*, 2021 | NA |
| LM358-18_2015_A_France-Reunion | Bacterial culture | South West Indian Ocean islands | *Citrus limon* | JAABFP000000000 | SRR11234698 | Richard *et al.*, 2021 | NA |
| LMG-9322_1986_A_USA | Bacterial culture | North America | *Citrus aurantiifolia* | JPYD00000000.1 | SRR11234695 | Gordon *et al.*, 2015 | NA |
| LN006-18_2016_A_France-Reunion | Bacterial culture | South West Indian Ocean islands | *Citrus reticulata* x *Citrus sinensis* | JAABDQ000000000 | SRR11234689 | Richard *et al.*, 2021 | NA |
| LP027-03_2017_A_Seychelles | Bacterial culture | North Indian Ocean islands | *Citrus aurantiifolia* | JAABIW000000000 | SRR11234681 | Richard *et al.*, 2021 | NA |
| LP027-05_2017_A_Seychelles | Bacterial culture | North Indian Ocean islands | *Citrus aurantiifolia* | JAABIX000000000 | SRR11234680 | Richard *et al.*, 2021 | NA |
| LP027-13_2017_A_Seychelles | Bacterial culture | North Indian Ocean islands | *Citrus aurantiifolia* | JAABIY000000000 | SRR11234679 | Richard *et al.*, 2021 | NA |
| LP028-03_2017_A_Seychelles | Bacterial culture | North Indian Ocean islands | *Citrus aurantiifolia* | JAABIZ000000000 | SRR11234678 | Richard *et al.*, 2021 | NA |
| LP028-05_2017_A_Seychelles | Bacterial culture | North Indian Ocean islands | *Citrus aurantiifolia* | JAABJA000000000 | SRR11234677 | Richard *et al.*, 2021 | NA |
| LP028-06_2017_A_Seychelles | Bacterial culture | North Indian Ocean islands | *Citrus aurantiifolia* | JAABJB000000000 | SRR11234676 | Richard *et al.*, 2021 | NA |
| LP029-13_2017_A_Seychelles | Bacterial culture | North Indian Ocean islands | *Citrus sinensis* | JAABJC000000000 | SRR11234675 | Richard *et al.*, 2021 | NA |
| LP029-15_2017_A_Seychelles | Bacterial culture | North Indian Ocean islands | *Citrus sinensis* | JAABJD000000000 | SRR11234673 | Richard *et al.*, 2021 | NA |
| LP030-1_2017_A_France-Martinique | Bacterial culture | Central America & Caribbean | *Citrus aurantiifolia* | JAABBM000000000 | SRR11234672 | Richard *et al.*, 2021 | NA |
| LP031-1_2017_A_France-Martinique | Bacterial culture | Central America & Caribbean | *Citrus sinensis* | JAABBN000000000 | SRR11234671 | Richard *et al.*, 2021 | NA |
| LP032-1_2017_A_France-Martinique | Bacterial culture | Central America & Caribbean | *Citrus aurantiifolia* | JAABBO000000000 | SRR11234670 | Richard *et al.*, 2021 | NA |
| LP033-1_2017_A_France-Martinique | Bacterial culture | Central America & Caribbean | *Citrus paradisi* | JAABBP000000000 | SRR11234669 | Richard *et al.*, 2021 | NA |
| LP034-1_2017_A_France-Martinique | Bacterial culture | Central America & Caribbean | *Citrus aurantiifolia* | JAABBQ000000000 | SRR11234668 | Richard *et al.*, 2021 | NA |
| LP035-1_2017_A_France-Martinique | Bacterial culture | Central America & Caribbean | *Citrus aurantiifolia* | JAABBR000000000 | SRR11234667 | Richard *et al.*, 2021 | NA |
| LP036-1_2017_A_France-Martinique | Bacterial culture | Central America & Caribbean | *Citrus aurantiifolia* | JAABBS000000000 | SRR11234666 | Richard *et al.*, 2021 | NA |
| LP162_2017_A_Pakistan | Bacterial culture | South Asia 2 | *Citrus reticulata* | | SRR21684649 | This study | NARC |
| LP187_2017_A_Pakistan | Bacterial culture | South Asia 2 | *Citrus sinensis* | | SRR21684648 | This study | NARC |
| LP188_2017_A_Pakistan | Bacterial culture | South Asia 2 | *Citrus sinensis* | | SRR21684647 | This study | NARC |
| LP201_2017_A_Pakistan | Bacterial culture | South Asia 2 | *Citrus* sp. | | SRR21684646 | This study | NARC |
| LP215-01_2017_A_Pakistan | Bacterial culture | South Asia 2 | *Citrus reticulata* | | SRR21684645 | This study | NARC |
| LP220-02_2017_A_Pakistan | Bacterial culture | South Asia 2 | *Citrus sinensis* | | SRR21684644 | This study | NARC |
| LP229-03_2017_A_Pakistan | Bacterial culture | South Asia 2 | *Citrus reticulata* | | SRR21684643 | This study | NARC |
| LP234-03_2017_A_Pakistan | Bacterial culture | South Asia 2 | *Citrus reticulata* | | SRR21684641 | This study | NARC |
| LP235-06_2017_A_Pakistan | Bacterial culture | South Asia 2 | *Citrus sinensis* | | SRR21684640 | This study | NARC |
| LP293-01_2017_A_Pakistan | Bacterial culture | South Asia 2 | *Citrus sinensis* | | SRR21684639 | This study | NARC |
| LP293-05_2017_A_Pakistan | Bacterial culture | South Asia 2 | *Citrus sinensis* | | SRR21684638 | This study | NARC |
| LP295-05_2017_A_Pakistan | Bacterial culture | South Asia 2 | *Citrus sinensis* | | SRR21684637 | This study | NARC |
| LP297-02_2017_A_Pakistan | Bacterial culture | South Asia 2 | *Citrus sinensis* | | SRR21684636 | This study | NARC |

| Label[1] | Source material | Geographic origin | Isolation Host | GenBank accession ID | Sequence Read Archive accession ID | Publication | Provider[2] |
|---|---|---|---|---|---|---|---|
| NCPPB-1471_1963_A_China-Hong-Kong | Bacterial culture | East Asia 1 | *Citrus paradisi* | | SRR21684635 | This study | NCPPB |
| NCPPB-211_1948_A_India | Bacterial culture | South Asia 1 | *Citrus* sp. | JAABAS000000000 | SRR11234665 | Richard *et al.*, 2021 | NA |
| NCPPB-226_1949_A_New-Zealand | Bacterial culture | Oceania & Pacific | *Citrus* sp. | JAABCW000000000 | SRR11234664 | Richard *et al.*, 2021 | NA |
| NCPPB-3562_1988_A_India | Bacterial culture | South Asia 1 | *Citrus limon* | CCXZ01000000 | SRR11234662 | Gordon *et al.*, 2015 | NA |
| NCPPB-3610_1988_A_India | Bacterial culture | South Asia 1 | *Poncirus trifoliata* | CDAO00000000.1 | SRR11234642 | Gordon *et al.*, 2015 | NA |
| NCPPB-3612_1988_A_India | Bacterial culture | South Asia 1 | *Citrus aurantiifolia* | CDAQ00000000.1 | SRR11234640 | Gordon *et al.*, 2015 | NA |
| NCPPB-406_1957_A_New-Zealand | Bacterial culture | Oceania & Pacific | *Citrus* sp. | JAABCX000000000 | SRR11234661 | Richard *et al.*, 2021 | NA |
| CFBP-2858_1977_Astar_Fiji | Bacterial culture | Oceania & Pacific | *Citrus aurantiifolia* | | SRR21684634 | This study | CFBP |
| JF90-2_1986_Astar_Oman | Bacterial culture | West Asia | *Citrus aurantiifolia* | CCWA00000000 | | Gordon *et al.*, 2015 | NA |
| JJ238-24_1989_Astar_Thailand | Bacterial culture | South East Asia 1 | *Citrus aurantiifolia* | CCVX00000000 | | Gordon *et al.*, 2015 | NA |
| JK143-11_1990_Astar_Thailand | Bacterial culture | South East Asia 1 | *Citrus* sp. | CDMO00000000 | | Gordon *et al.*, 2015 | NA |
| JK143-9_1990_Astar_Thailand | Bacterial culture | South East Asia 1 | *Citrus* sp. | CDMQ00000000 | | Gordon *et al.*, 2015 | NA |
| JK48_1988_Astar_Saudi-Arabia | Bacterial culture | West Asia | *Citrus aurantiifolia* | CDAJ00000000 | | Gordon *et al.*, 2015 | NA |
| JM35-2_1992_Astar_Saudi-Arabia | Bacterial culture | West Asia | *Citrus aurantiifolia* | CDMS00000000 | | Gordon *et al.*, 2015 | NA |
| JN658_1993_Astar_France-Reunion | Bacterial culture | South West Indian Ocean islands | *Citrus aurantiifolia* | | SRR21684633 | This study | CIRAD |
| JS581_1997_Astar_Iran | Bacterial culture | South Asia 2 | *Citrus limetta* | CDAW00000000 | | Gordon *et al.*, 2015 | NA |
| JS582_1997_Astar_Iran | Bacterial culture | South Asia 2 | *Citrus latifolia* | CDAP00000000 | | Gordon *et al.*, 2015 | NA |
| JS584_1997_Astar_Iran | Bacterial culture | South Asia 2 | *Citrus* sp. | CCWF00000000 | | Gordon *et al.*, 2015 | NA |
| LD71a_2007_Astar_Cambodia | Bacterial culture | South East Asia 1 | *Citrus* sp. | CCWE00000000 | | Gordon *et al.*, 2015 | NA |
| LE20-1_2008_Astar_Ethiopia | Bacterial culture | East Africa | *Citrus aurantiifolia* | CCWK00000000 | | Gordon *et al.*, 2015 | NA |
| LE3-1_2008_Astar_Ethiopia | Bacterial culture | East Africa | *Citrus aurantiifolia* | CDAI00000000 | | Gordon *et al.*, 2015 | NA |
| LK135-01_2013_Astar_Comoros-Moheli | Bacterial culture | South West Indian Ocean islands | *Citrus* sp. | | SRR21684632 | This study | INRAPE |
| NCPPB-3607_1988_Astar_India | Bacterial culture | South Asia 1 | *Citrus aurantiifolia* | CDAT00000000 | | Gordon *et al.*, 2015 | NA |
| NCPPB-3615_1989_Astar_India | Bacterial culture | South Asia 1 | *Citrus aurantiifolia* | CDAM00000000 | | Gordon *et al.*, 2015 | NA |
| JF90-8_1986_Aw_Oman | Bacterial culture | West Asia | *Citrus aurantiifolia* | CCWB00000000 | | Gordon *et al.*, 2015 | NA |
| LB302_2002_Aw_USA | Bacterial culture | North America | *Citrus aurantiifolia* | CDAU00000000 | | Gordon *et al.*, 2015 | NA |
| LG115_2007_Aw_India | Bacterial culture | South Asia 1 | *Citrus* sp. | CDAY00000000 | | Gordon *et al.*, 2015 | NA |
| NCPPB-3608_1988_Aw_India | Bacterial culture | South Asia 1 | *Citrus aurantiifolia* | CCWG00000000 | | Gordon *et al.*, 2015 | NA |
| LMG558_1950_cajani_India | Bacterial culture | South Asia 1 | *Cajanus cajan* | LOKQ00000000 | | Patané *et al.*, 2019 | NA |
| LMG9045_1974_clitoriae_India | Bacterial culture | South Asia 1 | *Clitoria* sp. | LOKA00000000 | | Patané *et al.*, 2019 | NA |
| NCPPB796_1960_vasculorum_Mauritius | Bacterial culture | South West Indian Ocean islands | *Saccharum* sp. | SAMN14772551 | | | NCPPB |

[1] Labels indicate: Strain number_year of isolation_pathotype (or pathovar for non-pv. *citri* strains)_country of origin.

[2] CFBP: Collection Française de Bactéries Phytopathogènes, France; FAO: Food and Agriculture Organization of the United Nations, China; FERA: Fera Science Ltd, UK; H4D: Horticulture4Development, Australia; INRAPE: Institut National de Recherche pour l'Agriculture, la Pêche et l'Environnement, Comoros; INTA: Instituto Nacional de Tecnología Agropecuaria, Argentina; NAQS: Northern Australia Quarantine Strategy, Australia; NARC: National Agricultural Research Center, Pakistan; NCHU: National Chung Hsing

University, Taiwan; NCPPB: National Collection of Plant Pathogenic Bacteria, UK; PPRI : Plant Protection Research Institute, Viet Nam; SOFRI: Southern Horticultural Research Institute, Viet Nam.

[3] See Table 1 for details.

**S2 Table. Summary of mapping, depth, coverage and damage statistics for reads of the 13 historic *Xci* mapping to plasmids pXAC33 and pXAC64 of the reference strain.** SD: standard deviation; nt: nucleotides.

| | | Endogenous *Xci* DNA (%) | | Mean depth | | Coverage at 1X (%) | | Insert length (mean ± SD in nt) | | Deamination rate at terminal position (%) | |
|---|---|---|---|---|---|---|---|---|---|---|---|
| **ID** | **Protocol** | **pXAC33** | **pXAC64** | **pXAC33** | **pXAC64** | **pXAC33** | **pXAC64** | **pXAC33** | **pXAC64** | **pXAC33** | **pXAC64** |
| HERB_1845 | TruSeq Nano | 0.36 | 0.48 | 122.2 | 120.4 | 93.3 | 95.7 | 52.2±22.8 | 51.9±22.6 | 3.91 | 3.90 |
| HERB_1884 | TruSeq Nano | 0.35 | 0.54 | 92.1 | 94.6 | 89.8 | 93.2 | 48.4±17.0 | 48.4±16.9 | 3.38 | 3.28 |
| HERB_1911 | TruSeq Nano | 0.06 | 0.08 | 69.2 | 65.2 | 90.2 | 90.2 | 69.7±21.9 | 69.4±21.8 | 3.42 | 3.78 |
| HERB_1915 | TruSeq Nano | 0.16 | 0.24 | 66.2 | 65.8 | 88 | 92.9 | 48.5±17.1 | 48.7±17.1 | 2.93 | 2.98 |
| HERB_1937 | TruSeq Nano | 0.03 | 0.04 | 22.6 | 17.9 | 82.9 | 88.5 | 44.7±13.8 | 44.6±13.6 | 3.99 | 4.09 |
| HERB_1946 | TruSeq Nano | 0.23 | 0.32 | 114.9 | 108.1 | 94.7 | 93.2 | 58.3±28.1 | 58.4±28.2 | 1.98 | 2.01 |
| HERB_1974 | TruSeq Nano | 0.12 | 0.14 | 42.6 | 25.5 | 82.5 | 49.7 | 41.5±9.5 | 41.6±9.5 | 2.46 | 2.56 |
| HERB_1852 | BEST | 0.17 | 0.23 | 130.3 | 117.9 | 95.9 | 96.5 | 70.4±41.4 | 69.7±41.1 | 2.13 | 2.17 |
| HERB_1854 | BEST | 0.39 | 0.54 | 114 | 108.6 | 95.2 | 97.1 | 72.0±39.6 | 72.3±39.7 | 1.35 | 1.40 |
| HERB_1859 | BEST | 0.16 | 0.22 | 84.1 | 75.4 | 94.2 | 95.3 | 71.7±31.6 | 71.4±31.4 | 1.93 | 2.03 |
| HERB_1865 | BEST | 0.13 | 0.20 | 95.1 | 101.5 | 77.9 | 96.6 | 85.1±36.8 | 85.1±36.7 | 1.81 | 1.81 |
| HERB_1922 | BEST | 0.60 | 0.99 | 99.6 | 128.7 | 76.5 | 94.5 | 67.7±29.2 | 67.2±28.5 | 1.62 | 1.54 |
| HERB_1963 | BEST | 0.55 | 0.67 | 106.8 | 86.2 | 94.3 | 96 | 73.2±31.0 | 73.0±30.9 | 1.46 | 1.53 |

**S3 Table. Recombining regions among 184 historical or modern *Xci* strains.** Positions are indicated relatively to the reference genome of strain IAPAR 306.

| Starting position | Ending position | Length |
|---|---|---|
| 3,095,443 | 3,119,514 | 24,071 |
| 3,799,050 | 3,805,347 | 6,297 |
| 4,257,584 | 4,630,760 | 373,176 |
| 4,959,140 | 4,988,729 | 29,589 |

**S4 Table. List and presence status of 144 pathogenicity-associated genes investigated among 184 historical or modern *Xci* strains.** A gene was considered present if its coding sequence reached at least 75% coverage of the corresponding CDS in the organism strain. *Xci*: *Xanthomonas citri* pv. *citri*; *Xe*: *Xanthomonas euvesicatoria*; *Xcc*: *Xanthomonas campestris* pv. *campestris*; *Xoc*: *Xanthomonas oryzae* pv. *oryzicola*; *Xoo*: *Xanthomonas oryzae* pv. *oryzae*; T3E: Type III effector; T3SS: Type III secretion system. Most genes were identified and characterized in: Alegria *et al.* (2004), Astua-Monge *et al.* (2005), Bansal *et al.* (2017), Facincani *et al.* (2014), Ferreira *et al.* (2017), Ferreira *et al.* (2016), Figueiredo *et al.* (2011), Gochez *et al.* (2017), Guo *et al.* (2011), Huang *et al.* (2021), Jalan *et al.* (2013), Laia *et al.* (2009), Moreira *et al.* (2010), Patané *et al.* (2019), Rybak *et al.* (2009), Yamazaki *et al.* (2008), Yan *et al.* (2012), Zhang *et al.* (2015)

| **Gene type** | **Gene family** | **CDS** | **Function** | **Organism strain** | **Gene status across *Xci* strains** |
|---|---|---|---|---|---|
| T3E | *avrBs1* | XCVd0104 | Unknown | *Xe* 85-10 | absent |
| T3E | *avrBs2* | XAC0076 | Glycerophosphoryl diester phosphodiesterase | *Xci* IAPAR 306 | present |
| T3E | *avrBs3* | XACb0065 (PthA4) | AvrBs3/PthA-type transcription activator; nuclear localisation | *Xci* IAPAR 306 | present |
| T3E | *avrXccA1* | XCC4229 | Unknown, maybe not a T3E | *Xcc* ATCC 33913 | absent |
| T3E | *avrXccA2* | XCC2396 | Unknown, maybe not a T3E | *Xcc* ATCC 33913 | absent |
| T3E | *hpaI* | XANAC_0475 | Type III effector HpaI protein (fragment) | *Xci* IAPAR 306 | present |
| T3E | *hrpW* | XAC2922 | Pectate lyase, maybe not a T3E | *Xci* IAPAR 306 | present |
| T3E | *xopA* | XAC0416 | Harpin, maybe not a T3E | *Xci* IAPAR 306 | present |
| T3E | *xopAA* | XCV3785 | Early chlorosis factor; Proteasome/cyclosome repeat | *Xe* 85-10 | absent |
| T3E | *xopAB* | XOO3150_MAFF | Unknown | *Xoo* MAFF 311018 | absent |
| T3E | *xopAC* | XCC2565_ATCC33913 | LRR-Fic/DOC protein; vascular hypersensitivity in *Arabidopsis landrace* Col-0 | *Xcc* ATCC 33913 | absent |
| T3E | *xopAD* | XAC4213 | SKWP repeat protein | *Xci IAPAR 306* | present |
| T3E | *xopAE* | XAC0393 | LRR protein | *Xci* IAPAR 306 | present |
| T3E | *xopAF* | XCAW_b00003 | Unknown | *Xci XacW12879* | variable presence |
| T3E | *xopAG* | JX566667 | Unknown | *Xci* NCPPB 3608 | variable presence |
| T3E | *xopAH* | XCC2109_ATCC33913 | Unknown | *Xcc* ATCC 33913 | absent |
| T3E | *xopAI* | XAC3230 | Putative ADP-ribosyltransferase | *Xci* IAPAR 306 | present |
| T3E | *xopAJ* | XCV4428 | Unknown | *Xe* 85-10 | absent |
| T3E | *xopAK* | XAC3666 | Unknown | *Xci* IAPAR 306 | present |
| T3E | *xopAL1* | XCC1246_ATCC33913 | Unknown | *Xcc* ATCC 33913 | absent |
| T3E | *xopAL2* | XCCB100_0616 | Unknown | *Xcc* B100 | absent |
| T3E | *xopAM* | XCC1089_ATCC33913 | Unknown | *Xcc* ATCC 33913 | absent |
| T3E | *xopAP* | XAC2990 | Unknown | *Xci* IAPAR 306 | present |
| T3E | *xopAQ* | XAC40v3_800003 | Unknown | *Xci* C40 | present |
| T3E | *xopAU* | XAC1171 | Serine/threonine kinase | *Xci* IAPAR 306 | present |
| T3E | *xopAV* | XAC1172 | Unknown | *Xci* IAPAR 306 | variable presence |
| T3E | *xopAW* | XAC2949 | Calcium-binding protein | *Xci* IAPAR 306 | present |
| T3E | *xopAX* | XCVd0086 | Unknown | *Xe* 85-10 | absent |

| | | | | | |
|---|---|---|---|---|---|
| T3E | *xopAY* | XAC1172 | Unknown | *Xci* IAPAR 306 | present |
| T3E | *xopAZ* | XAC1358 | SlpA superfamily, FKBP-type peptidyl-prolyl cis-trans isomerase | *Xci* IAPAR 306 | present |
| T3E | *xopB* | XCV0581 | Unknown | *Xe* 85-10 | absent |
| T3E | *xopC1* | JX566666 | Phosphoribosyl transferase domain and haloacid dehalogenase-like hydrolase | *Xci JK2-10* | variable presence |
| T3E | *xopC2* | XAC1210_Ψ; XAC1209_Ψ | Haloacid dehalogenase-like hydrolase | *Xci* IAPAR 306 | present |
| T3E | *xopD* | XCV0437 | C48-family SUMO cysteine protease (Ulp1 protease family) (Clan CE); EAR motif; DNA binding; nuclear localisation | *Xe* 85-10 | absent |
| T3E | *xopE1* | XAC0286 | Putative transglutaminase | *Xci* IAPAR 306 | present |
| T3E | *xopE2* | XACb0011 | Putative transglutaminase, plasmidic | *Xci* IAPAR 306 | variable presence |
| T3E | *xopE3* | XAC3224 | Putative transglutaminase | *Xci* IAPAR 306 | present |
| T3E | *xopF1* | XANAC_0476_Ψ; XANAC_0477_ Ψ | Unknown | *Xci* IAPAR 306 | present |
| T3E | *xopF1* | XCV0414 | Unknown | *Xe* 85-10 | absent |
| T3E | *xopF2* | XAC2785_Ψ | Unknown | *Xci* IAPAR 306 | present |
| T3E | *xopF2* | XCV2942 | Unknown | *Xe* 85-10 | absent |
| T3E | *xopG1* | XCV1298 | M27-family peptidase (Clostridium toxin) | *Xe* 85-10 | absent |
| T3E | *xopG2* | XCC3258 | M27-family peptidase (Clostridium toxin) | *Xcc* ATCC 33913 | absent |
| T3E | *xopH1* | XCVd0105 | Putative tyrosine phosphatase | *Xe* 85-10 | absent |
| T3E | *xopI1* | XAC0754 | F-box protein | *Xci* IAPAR 306 | present |
| T3E | *xopJ1* | XCV2156 | C55-family cysteine protease or Ser/Thr acetyltransferase (Clan CE) | *Xe* 85-10 | absent |
| T3E | *xopJ2* | Aave_2166 | C55-family cysteine protease or Ser/Thr acetyltransferase (Clan CE) | *Acidovorax citrulli* AAC00-1 | absent |
| T3E | *xopJ3* | XCV0471 | C55-family cysteine protease or Ser/Thr acetyltransferase (Clan CE) | *Xe* 85-10 | absent |
| T3E | *xopJ4* | RSc0826 | C55-family cysteine protease or Ser/Thr acetyltransferase (Clan CE) | *Ralstonia solanacearum* GMI1000 | absent |
| T3E | *xopJ5* | XCC3731_ATCC33913 | Putative C55-family cysteine protease or Ser/Thr acetyltransferase (Clan CE) | *Xcc* ATCC 33913 | absent |
| T3E | *xopK* | XAC3085 | Unknown | *Xci* IAPAR 306 | present |
| T3E | *xopL* | XAC3090 | LRR protein | *Xci* IAPAR 306 | present |
| T3E | *xopM* | XAC0418 | Unknown | *Xci* IAPAR 306 | present |
| T3E | *xopN* | XAC2786 | ARM/HEAT repeat | *Xci* IAPAR 306 | present |
| T3E | *xopO* | XCV1055 | Unknown | *Xe* 85-10 | absent |
| T3E | *xopP* | XAC1208 | Unknown | *Xci* IAPAR 306 | present |
| T3E | *xopQ* | XAC4333 | Putative inosine-uridine nucleoside N-ribohydrolase | *Xci* IAPAR 306 | present |
| T3E | *xopR* | XAC0277 | Unknown | *Xci* IAPAR 306 | present |
| T3E | *xopS* | XAC0315 | Unknown | *Xci* IAPAR 306 | present |
| T3E | *xopT* | XANAC_p10046 | Unknown | *Xci* IAPAR 306 | variable presence |
| T3E | *xopU* | PXO_00236 | Unknown | *Xoo* PX099A | absent |
| T3E | *xopV* | XAC0601 | Unknown | *Xci* IAPAR 306 | present |
| T3E | *xopW* | PXO_03356 | Unknown | *Xoo* PX099A | absent |
| T3E | *xopX* | XAC0543 | Unknown | *Xci* IAPAR 306 | present |
| T3E | *xopY* | XOO1488_MAFF | Unknown | *Xoo* MAFF 311018 | absent |

| | | | | | |
|---|---|---|---|---|---|
| T3E | *xopZ1* | XAC2009 | Unknown | *Xci* IAPAR 306 | present |
| T3SS | *hpaA* | XAC0400 | Type III secretion control protein, maybe not a T3E | *Xci* IAPAR 306 | present |
| T3SS | *hpaB* | XAC0396 | Type III secretion system chaperone | *Xci* IAPAR 306 | present |
| T3SS | *hpaC* | XAC0404 | Type III secretion system export control protein | *Xci* IAPAR 306 | present |
| T3SS | *hpaH* | XAC0417 | Type III secretion system putative transglycosylase HpaH | *Xci* IAPAR 306 | present |
| T3SS | *hrcC* | XAC0415 | Type III secretion system outer membrane pore protein | *Xci* IAPAR 306 | present |
| T3SS | *hrcD* | XAC0399 | Type III secretion system protein | *Xci* IAPAR 306 | present |
| T3SS | *hrcJ* | XAC0409 | Type III secretion bridge between inner and outermembrane lipoprotein | *Xci* IAPAR 306 | present |
| T3SS | *hrcL* | XAC0411 | Type III secretion system cytoplasmic protein | *Xci* IAPAR 306 | present |
| T3SS | *hrcN* | XAC0412 | Type III secretion system ATP synthase | *Xci* IAPAR 306 | present |
| T3SS | *hrcQ* | XAC0403 | Type III secretion system apparatus protein | *Xci* IAPAR 306 | present |
| T3SS | *hrcR* | XAC0402 | Type III secretion system inner membrane protein | *Xci* IAPAR 306 | present |
| T3SS | *hrcS* | XAC0401 | Type III secretion system inner membrane protein | *Xci* IAPAR 306 | present |
| T3SS | *hrcT* | XAC0414 | Type III secretion system inner membrane protein | *Xci* IAPAR 306 | present |
| T3SS | *hrcU* | XAC0406 | Type III secretion system inner membrane protein | *Xci* IAPAR 306 | present |
| T3SS | *hrcV* | XAC0405 | Type III secretion system inner membrane channel protein | *Xci* IAPAR 306 | present |
| T3SS | *hrpB1* | XAC0407 | Type III secretion system protein | *Xci* IAPAR 306 | present |
| T3SS | *hrpB2* | XAC0408 | Type III secretion system protein | *Xci* IAPAR 306 | present |
| T3SS | *hrpB4* | XAC0410 | Type III secretion system protein | *Xci* IAPAR 306 | present |
| T3SS | *hrpB7* | XAC0413 | Type III secretion system protein | *Xci* IAPAR 306 | present |
| T3SS regulator | *hrpD6* | XAC0398 | Type III secretion system regulator | *Xci* IAPAR 306 | present |
| T3SS | *hrpE* | XAC0397 | Type III secretion system pilin | *Xci* IAPAR 306 | present |
| T3SS | *hrpF* | XAC0394 | Type III secretion system translocator protein | *Xci* IAPAR 306 | present |
| T3SS regulator | *hrpG* | XAC1265 | Type III secretion system OmpR-type response regulator | *Xci* IAPAR 306 | present |
| T3SS regulator | *hrpX* | XAC1266 | Type III secretion system transcriptional activator | *Xci* IAPAR 306 | present |
| | *clpA* | XAC_RS10175 | ATP-dependent Clp protease ATP-binding subunit ClpA | *Xci* IAPAR 306 | present |
| | *dksA* | XAC_RS23635 | RNA polymerase-binding protein DksA | *Xci* IAPAR 306 | present |
| | *exbD1* | XAC_RS00055 | Biopolymer transporter ExbD | *Xci* IAPAR 306 | present |
| | *exsF* | XAC_RS15890 | Response regulator | *Xci* IAPAR 306 | present |
| | *exsG* | XAC_RS15895 | PAS domain S-box protein | *Xci* IAPAR 306 | present |
| | *galU* | XAC_RS11675 | UTP—glucose-1-phosphate uridylyltransferase GalU | *Xci* IAPAR 306 | present |
| | *gltB* | XAC_RS00175 | Glutamate syntase large subunit | *Xci* IAPAR 306 | present |
| | *gumF* | XAC_RS13145 | Acyltransferase | *Xci* IAPAR 306 | present |
| | *gumK* | XAC_RS13120 | Glycosyltransferase | *Xci* IAPAR 306 | present |
| | *helD* | XAC2265 | Helicase IV | *Xci* IAPAR 306 | variable presence |
| | *hemB* | XAC_RS20350 | Porphobilinogen synthase | *Xci* IAPAR 306 | present |
| | *hrcN* | XAC_RS02160 | EscN/YscN/HrcN family type III secretion system ATPase | *Xci* IAPAR 306 | present |
| | *hrpM* | XAC_RS03215 | Glucans biosynthesis glucosyltransferase MdoH | *Xci* IAPAR 306 | present |
| | *htrA* | XAC_RS20055 | Do family serine endopeptidase | *Xci* IAPAR 306 | present |

| | | | | | |
|---|---|---|---|---|---|
| | *ispF* | XAC_RS08775 | 2-C-methyl-D-erythritol 2 4-cyclodiphosphate synthase | *Xci* IAPAR 306 | present |
| | *leuC* | XAC_RS17505 | 3-isopropylmalate dehydratase large subunit | *Xci* IAPAR 306 | present |
| | *nadD* | XAC_RS14105 | Nicotinate-nucleotide adenylyltransferase | *Xci* IAPAR 306 | present |
| | *ostA* | XAC_RS16280 | Alpha alpha-trehalose-phosphate synthase (UDP-forming) | *Xci* IAPAR 306 | present |
| | *peh-1* | XAC_RS03430 | Endopolygalacturonase | *Xci* IAPAR 306 | present |
| | *pgi* | XAC_RS09105 | Glucose-6-phosphate isomerase | *Xci* IAPAR 306 | present |
| | *pstB* | XAC_RS08000 | Phosphate ABC transporter ATP-binding protein PstB | *Xci* IAPAR 306 | present |
| | *tatB* | XAC_RS21270 | Twin-arginine translocase subunit TatB | *Xci* IAPAR 306 | present |
| | *tatC* | XAC_RS21265 | Twin-arginine translocase subunit TatC | *Xci* IAPAR 306 | present |
| | *trpC* | XAC_RS02505 | Indole-3-glycerol phosphate synthase TrpC | *Xci* IAPAR 306 | present |
| | *xpsD* | XAC_RS17865 | Type II secretion system protein GspD | *Xci* IAPAR 306 | present |
| | *xpsE* | XAC_RS17915 | Type II secretion system protein GspE | *Xci* IAPAR 306 | present |
| | *xpsF* | XAC_RS17910 | Type II secretion system F family protein | *Xci* IAPAR 306 | present |
| | *xpsG* | XAC_RS17905 | Type II secretion system protein GspG | *Xci* IAPAR 306 | present |
| | *xpsM* | XAC_RS17875 | General secretion pathway protein GspM | *Xci* IAPAR 306 | present |
| | *xpsN* | XAC_RS17870 | Hypothetical protein | *Xci* IAPAR 306 | present |
| | *yapH* | XAC_RS20725 | Filamentous hemagglutinin N-terminal domain-containing protein | *Xci* IAPAR 306 | present |
| | | XAC3278 | BLUF domain-containing protein | *Xci* IAPAR 306 | variable presence |
| | | XAC1496 | Uncharacterised protein | *Xci* IAPAR 306 | variable presence |
| | | XAC3294 | Uncharacterised protein | *Xci* IAPAR 306 | variable presence |
| | | XAC_RS05830 | 2-methylaconitate cis-trans isomerase PrpF | *Xci* IAPAR 306 | present |
| | | XAC_RS05275 | Amidophosphoribosyltransferase | *Xci* IAPAR 306 | present |
| | | XAC_RS02490 | Aminodeoxychorismate/anthranilate synthase component II | *Xci* IAPAR 306 | present |
| | | XAC_RS05160 | Cell wall hydrolase | *Xci* IAPAR 306 | present |
| | | XAC_RS15775 | Glycosyltransferase | *Xci* IAPAR 306 | present |
| | | XAC_RS07880 | GntR family transcriptional regulator | *Xci* IAPAR 306 | present |
| | | XAC_RS06130 | HDOD domain-containing protein | *Xci* IAPAR 306 | present |
| | | XAC_RS01780 | Helix-turn helix transcriptional regulator | *Xci* IAPAR 306 | present |
| | | XAC_RS06460 | Helix-turn helix transcriptional regulator | *Xci* IAPAR 306 | present |
| | | XAC_RS00350 | Hypothetical protein | *Xci* IAPAR 306 | present |
| | | XAC_RS03785 | Hypothetical protein | *Xci* IAPAR 306 | present |
| | | XAC_RS15480 | Hypothetical protein | *Xci* IAPAR 306 | present |
| | | XAC_RS22275 | Lytic murein transglycosylase | *Xci* IAPAR 306 | present |
| | | XAC_RS17065 | M23 family metallopeptidase | *Xci* IAPAR 306 | present |
| | | XAC_RS19665 | M23 family metallopeptidase | *Xci* IAPAR 306 | present |
| | | XAC_RS09355 | Methylthioribulose 1-phosphate dehydratase | *Xci* IAPAR 306 | present |
| | | XAC_RS04070 | Nudix family hydrolase | *Xci* IAPAR 306 | present |
| | | XAC_RS00125 | Peptidase M23 | *Xci* IAPAR 306 | present |
| | | XAC_RS06280 | Polyketide cyclase | *Xci* IAPAR 306 | present |
| | | XAC_RS18580 | Response regulator | *Xci* IAPAR 306 | present |

**S5 Table. Coverage (in percentage) of the 10 pathogenicity-associated genes of variable presence investigated among 184 historical or modern *Xci* strains.**

| **Strains** | ***xopAF*** | ***xopAG*** | ***xopAV*** | ***xopC1*** | ***xopE2*** | ***xopT*** | **XAC1496** | **XAC2265** | **XAC3278** | **XAC3294** |
|---|---|---|---|---|---|---|---|---|---|---|
| C20_1988_A_France-Reunion | 0.00 | 0.00 | 100.00 | 0.00 | 100.00 | 100.00 | 100.00 | 100.00 | 100.00 | 100.00 |
| CFBP-1209_1963_A_China-Hong-Kong | 0.00 | 0.00 | 100.00 | 0.00 | 100.00 | 100.00 | 100.00 | 100.00 | 100.00 | 100.00 |
| CFBP-2525_1956_A_New-Zealand | 0.00 | 0.00 | 100.00 | 0.00 | 100.00 | 100.00 | 100.00 | 100.00 | 100.00 | 100.00 |
| CFBP-2853_1958_A_New-Zealand | 0.00 | 0.00 | 100.00 | 0.00 | 100.00 | 100.00 | 100.00 | 100.00 | 100.00 | 100.00 |
| CFBP-2855_1962_A_Japan | 0.00 | 0.00 | 100.00 | 0.00 | 100.00 | 100.00 | 100.00 | 100.00 | 100.00 | 100.00 |
| CFBP-2857_1976_A_Fiji | 0.00 | 0.00 | 100.00 | 0.00 | 100.00 | 100.00 | 100.00 | 100.00 | 100.00 | 100.00 |
| CFBP-2858_1977_Astar_Fiji | 100.00 | 0.00 | 100.00 | 0.00 | 100.00 | 93.25 | 0.00 | 0.00 | 0.00 | 100.00 |
| CFBP-2865_1976_A_Brazil | 9.07 | 0.00 | 100.00 | 0.00 | 100.00 | 100.00 | 100.00 | 100.00 | 100.00 | 100.00 |
| CFBP-2908_1980_A_Brazil | 0.00 | 0.00 | 100.00 | 0.00 | 100.00 | 100.00 | 100.00 | 100.00 | 100.00 | 100.00 |
| FDC-1083_1980_A_Brazil | 0.00 | 0.00 | 100.00 | 21.20 | 100.00 | 100.00 | 100.00 | 100.00 | 100.00 | 100.00 |
| FDC-217_2003_A_Brazil | 8.60 | 0.00 | 100.00 | 0.00 | 100.00 | 100.00 | 100.00 | 100.00 | 100.00 | 100.00 |
| HERB_1845_A_Indonesia | 4.12 | 0.00 | 99.73 | 0.60 | 100.00 | 100.00 | 100.00 | 100.00 | 100.00 | 99.72 |
| HERB_1852_A_India | 100.00 | 1.24 | 99.60 | 0.00 | 100.00 | 100.00 | 100.00 | 99.89 | 99.32 | 99.17 |
| HERB_1854_A_Indonesia | 3.89 | 0.00 | 99.60 | 0.00 | 100.00 | 100.00 | 100.00 | 99.89 | 99.77 | 99.44 |
| HERB_1859_A_Bangladesh | 99.53 | 0.00 | 98.93 | 0.00 | 100.00 | 100.00 | 99.90 | 99.89 | 99.55 | 98.61 |
| HERB_1865_A_India | 0.00 | 0.00 | 99.46 | 0.00 | 100.00 | 47.37 | 100.00 | 99.79 | 99.55 | 100.00 |
| HERB_1884_A_Philippines | 0.00 | 1.95 | 99.73 | 0.64 | 100.00 | 100.00 | 98.88 | 100.00 | 100.00 | 99.72 |
| HERB_1911_A_Indonesia | 0.00 | 0.00 | 99.20 | 0.00 | 100.00 | 99.89 | 99.69 | 99.86 | 99.32 | 99.17 |
| HERB_1915_A_Philippines | 0.00 | 0.00 | 99.60 | 0.00 | 99.91 | 99.68 | 99.49 | 99.93 | 99.55 | 98.33 |
| HERB_1922_A_China | 100.00 | 0.00 | 99.60 | 0.00 | 100.00 | 66.99 | 100.00 | 99.96 | 100.00 | 100.00 |
| HERB_1937_A_Mauritius | 0.00 | 0.00 | 96.65 | 0.00 | 99.74 | 98.39 | 85.02 | 99.61 | 91.22 | 98.61 |
| HERB_1946_A_Guam | 0.00 | 0.00 | 99.60 | 0.00 | 97.90 | 99.89 | 100.00 | 99.96 | 100.00 | 100.00 |
| HERB_1963_A_Nepal | 0.00 | 0.00 | 99.60 | 0.00 | 99.83 | 99.89 | 99.90 | 99.96 | 98.65 | 98.89 |
| HERB_1974_A_Mauritius | 3.53 | 0.00 | 99.60 | 1.72 | 99.91 | 99.68 | 96.94 | 99.93 | 99.55 | 98.89 |
| JF90-2_1986_Astar_Oman | 100.00 | 4.04 | 100.00 | 100.00 | 100.00 | 93.35 | 0.00 | 11.13 | 100.00 | 100.00 |
| JF90-8_1986_Aw_Oman | 100.00 | 100.00 | 100.00 | 0.00 | 35.78 | 11.58 | 100.00 | 100.00 | 100.00 | 100.00 |
| JH081-04_1988_A_China | 0.00 | 0.00 | 100.00 | 0.00 | 100.00 | 100.00 | 100.00 | 100.00 | 100.00 | 100.00 |
| JH410-04_1988_A_China | 0.00 | 0.00 | 100.00 | 0.00 | 100.00 | 100.00 | 100.00 | 100.00 | 100.00 | 100.00 |
| JJ009-04_1985_A_Mauritius | 0.00 | 0.00 | 100.00 | 0.00 | 100.00 | 100.00 | 100.00 | 100.00 | 100.00 | 100.00 |
| JJ009-1_1984_A_Mauritius | 0.00 | 0.00 | 100.00 | 0.00 | 100.00 | 100.00 | 100.00 | 100.00 | 100.00 | 100.00 |
| JJ009-8_1987_A_Mauritius | 0.00 | 0.00 | 100.00 | 0.00 | 100.00 | 100.00 | 100.00 | 100.00 | 100.00 | 100.00 |
| JJ037-01_1989_A_Malaysia | 0.00 | 0.00 | 100.00 | 0.00 | 100.00 | 100.00 | 100.00 | 100.00 | 100.00 | 100.00 |
| JJ053-04_1977_A_Taiwan | 0.00 | 0.00 | 100.00 | 0.00 | 100.00 | 100.00 | 100.00 | 100.00 | 100.00 | 100.00 |
| JJ053-06_1977_A_Taiwan | 0.00 | 0.00 | 100.00 | 0.00 | 100.00 | 100.00 | 100.00 | 100.00 | 100.00 | 100.00 |
| JJ053-07_1977_A_Taiwan | 0.00 | 0.00 | 100.00 | 0.00 | 100.00 | 100.00 | 100.00 | 100.00 | 100.00 | 100.00 |
| JJ053-09_1977_A_Taiwan | 32.74 | 0.00 | 100.00 | 0.00 | 100.00 | 100.00 | 100.00 | 100.00 | 100.00 | 100.00 |
| JJ155_1977_A_Argentina | 0.00 | 0.00 | 100.00 | 0.00 | 100.00 | 100.00 | 100.00 | 100.00 | 100.00 | 100.00 |
| JJ165_1978_A_India | 0.00 | 0.00 | 100.00 | 0.00 | 100.00 | 100.00 | 100.00 | 100.00 | 100.00 | 100.00 |
| JJ238-04_1987_A_Maldives | 0.00 | 0.00 | 100.00 | 0.00 | 100.00 | 100.00 | 100.00 | 100.00 | 100.00 | 100.00 |
| JJ238-06_1987_A_Maldives | 0.00 | 0.00 | 100.00 | 0.00 | 100.00 | 100.00 | 100.00 | 100.00 | 100.00 | 100.00 |
| JJ238-07_1987_A_Maldives | 0.00 | 0.00 | 100.00 | 0.00 | 100.00 | 100.00 | 100.00 | 100.00 | 100.00 | 100.00 |
| JJ238-08_1987_A_Maldives | 0.00 | 0.00 | 100.00 | 0.00 | 100.00 | 100.00 | 100.00 | 100.00 | 100.00 | 100.00 |
| JJ238-09_1987_A_Maldives | 0.00 | 0.00 | 100.00 | 0.00 | 100.00 | 100.00 | 100.00 | 100.00 | 100.00 | 100.00 |
| JJ238-10_1987_A_Maldives | 16.73 | 0.00 | 100.00 | 7.35 | 100.00 | 100.00 | 100.00 | 100.00 | 100.00 | 100.00 |
| JJ238-11_1987_A_Maldives | 0.00 | 0.00 | 100.00 | 0.00 | 100.00 | 100.00 | 100.00 | 100.00 | 0.00 | 100.00 |
| JJ238-14_1988_A_Philippines | 0.00 | 0.00 | 100.00 | 0.00 | 100.00 | 100.00 | 100.00 | 100.00 | 100.00 | 100.00 |
| JJ238-16_1988_A_Pakistan | 0.00 | 0.00 | 100.00 | 0.00 | 100.00 | 100.00 | 100.00 | 100.00 | 100.00 | 100.00 |

| | | | | | | | | | | |
|---|---|---|---|---|---|---|---|---|---|---|
| JJ238-24_1989_Astar_Thailand | 100.00 | 0.00 | 100.00 | 0.00 | 100.00 | 11.36 | 0.00 | 71.60 | 0.00 | 98.61 |
| JK004-04_1987_A_Maldives | 0.00 | 0.00 | 100.00 | 0.00 | 100.00 | 100.00 | 100.00 | 100.00 | 100.00 | 100.00 |
| JK101-1_1990_A_Argentina | 0.00 | 0.00 | 100.00 | 0.00 | 100.00 | 100.00 | 100.00 | 100.00 | 100.00 | 100.00 |
| JK143-11_1990_Astar_Thailand | 100.00 | 0.00 | 100.00 | 16.13 | 100.00 | 29.05 | 10.30 | 74.04 | 22.75 | 98.61 |
| JK143-9_1990_Astar_Thailand | 100.00 | 0.00 | 100.00 | 4.03 | 100.00 | 11.68 | 9.48 | 71.60 | 0.00 | 98.61 |
| JK146-01_1990_A_Malaysia | 0.00 | 0.00 | 100.00 | 0.00 | 100.00 | 100.00 | 100.00 | 100.00 | 100.00 | 100.00 |
| JK146-02_1990_A_Malaysia | 0.00 | 0.00 | 100.00 | 0.00 | 100.00 | 100.00 | 100.00 | 100.00 | 100.00 | 100.00 |
| JK146-03_1990_A_Malaysia | 35.34 | 0.00 | 100.00 | 0.00 | 100.00 | 100.00 | 100.00 | 100.00 | 100.00 | 100.00 |
| JK146-05_1990_A_Malaysia | 0.00 | 0.00 | 100.00 | 0.00 | 100.00 | 100.00 | 100.00 | 100.00 | 100.00 | 100.00 |
| JK148-01_1990_A_Philippines | 0.00 | 0.00 | 100.00 | 0.00 | 100.00 | 100.00 | 100.00 | 100.00 | 100.00 | 100.00 |
| JK148-04_1990_A_Philippines | 0.00 | 0.00 | 100.00 | 0.00 | 100.00 | 100.00 | 100.00 | 100.00 | 100.00 | 100.00 |
| JK148-07_1990_A_Philippines | 0.00 | 0.00 | 100.00 | 0.00 | 100.00 | 100.00 | 100.00 | 100.00 | 100.00 | 100.00 |
| JK148-11_1990_A_Philippines | 0.00 | 0.00 | 100.00 | 0.00 | 100.00 | 100.00 | 100.00 | 100.00 | 100.00 | 100.00 |
| JK148-13_1990_A_Philippines | 0.00 | 0.00 | 100.00 | 0.00 | 100.00 | 100.00 | 100.00 | 100.00 | 100.00 | 100.00 |
| JK161-4_1990_A_Mauritius | 0.00 | 0.00 | 100.00 | 0.00 | 100.00 | 100.00 | 100.00 | 100.00 | 100.00 | 100.00 |
| JK4-1_1985_A_China | 18.37 | 0.00 | 100.00 | 0.00 | 100.00 | 100.00 | 100.00 | 100.00 | 100.00 | 100.00 |
| JK48_1988_Astar_Saudi-Arabia | 100.00 | 6.25 | 100.00 | 100.00 | 100.00 | 18.54 | 9.07 | 0.00 | 100.00 | 100.00 |
| JM35-2_1992_Astar_Saudi-Arabia | 100.00 | 0.00 | 100.00 | 100.00 | 100.00 | 92.82 | 9.38 | 12.20 | 100.00 | 100.00 |
| JN658_1993_Astar_France-Reunion | 100.00 | 0.00 | 100.00 | 0.00 | 100.00 | 93.25 | 0.00 | 0.00 | 100.00 | 100.00 |
| JS538-2_1997_A_France-Reunion | 0.00 | 0.00 | 100.00 | 0.00 | 100.00 | 100.00 | 100.00 | 100.00 | 100.00 | 100.00 |
| JS581_1997_Astar_Iran | 100.00 | 0.00 | 100.00 | 100.00 | 100.00 | 41.59 | 0.00 | 15.60 | 100.00 | 100.00 |
| JS582_1997_Astar_Iran | 100.00 | 0.00 | 100.00 | 100.00 | 100.00 | 11.25 | 0.00 | 6.95 | 100.00 | 100.00 |
| JS584_1997_Astar_Iran | 100.00 | 0.00 | 100.00 | 100.00 | 100.00 | 11.25 | 0.00 | 0.00 | 100.00 | 100.00 |
| JW160-1_2000_A_Bangladesh | 100.00 | 0.00 | 100.00 | 9.94 | 100.00 | 100.00 | 100.00 | 100.00 | 100.00 | 100.00 |
| JZ092_2003_A_Seychelles | 0.00 | 0.00 | 100.00 | 0.00 | 100.00 | 100.00 | 100.00 | 100.00 | 100.00 | 100.00 |
| JZ094-4_2003_A_Seychelles | 0.00 | 0.00 | 100.00 | 0.00 | 100.00 | 100.00 | 100.00 | 100.00 | 100.00 | 100.00 |
| LB100-1_2005_A_Seychelles | 5.30 | 0.00 | 100.00 | 4.03 | 100.00 | 100.00 | 100.00 | 100.00 | 100.00 | 100.00 |
| LB100-3_2005_A_Seychelles | 0.00 | 0.00 | 100.00 | 0.00 | 100.00 | 100.00 | 100.00 | 100.00 | 100.00 | 100.00 |
| LB302_2002_Aw_USA | 100.00 | 100.00 | 100.00 | 3.71 | 100.00 | 42.55 | 100.00 | 100.00 | 0.00 | 26.11 |
| LC004-1_2006_A_Vietnam | 0.00 | 0.00 | 100.00 | 0.00 | 100.00 | 100.00 | 100.00 | 100.00 | 100.00 | 100.00 |
| LC008-04_2006_A_Vietnam | 5.89 | 0.00 | 100.00 | 0.00 | 100.00 | 100.00 | 100.00 | 100.00 | 100.00 | 100.00 |
| LC014-10_2006_A_Vietnam | 0.00 | 0.00 | 100.00 | 0.00 | 100.00 | 100.00 | 100.00 | 100.00 | 100.00 | 100.00 |
| LC015-01_2006_A_Vietnam | 27.21 | 0.00 | 100.00 | 0.00 | 100.00 | 100.00 | 100.00 | 100.00 | 100.00 | 100.00 |
| LC052-05_2006_A_Vietnam | 4.83 | 0.00 | 100.00 | 0.00 | 100.00 | 100.00 | 100.00 | 100.00 | 100.00 | 100.00 |
| LC060-08_2006_A_Vietnam | 0.00 | 0.00 | 100.00 | 0.00 | 100.00 | 100.00 | 100.00 | 100.00 | 100.00 | 100.00 |
| LC067-02_2006_A_Vietnam | 0.00 | 0.00 | 100.00 | 0.00 | 100.00 | 100.00 | 100.00 | 100.00 | 0.00 | 100.00 |
| LC067-04_2006_A_Vietnam | 39.22 | 0.00 | 100.00 | 0.00 | 100.00 | 100.00 | 100.00 | 100.00 | 58.33 | 100.00 |
| LC80_2006_A_Mali | 100.00 | 0.00 | 100.00 | 100.00 | 100.00 | 100.00 | 100.00 | 100.00 | 100.00 | 100.00 |
| LD072-01_2007_A_Myanmar | 0.00 | 0.00 | 100.00 | 0.00 | 100.00 | 100.00 | 100.00 | 100.00 | 100.00 | 100.00 |
| LD072-02_2007_A_Myanmar | 30.74 | 0.00 | 100.00 | 0.00 | 100.00 | 100.00 | 100.00 | 100.00 | 100.00 | 100.00 |
| LD107-01_2007_A_Indonesia | 0.00 | 0.00 | 56.76 | 0.00 | 100.00 | 100.00 | 100.00 | 100.00 | 100.00 | 100.00 |
| LD107-02_2007_A_Indonesia | 0.00 | 0.00 | 100.00 | 0.00 | 100.00 | 100.00 | 100.00 | 100.00 | 100.00 | 100.00 |

| | | | | | | | | | | |
|---|---|---|---|---|---|---|---|---|---|---|
| LD108-01_2007_A_Indonesia | 0.00 | 0.00 | 56.76 | 0.00 | 100.00 | 100.00 | 100.00 | 100.00 | 100.00 | 100.00 |
| LD108-02_2007_A_Indonesia | 0.00 | 0.00 | 100.00 | 0.00 | 100.00 | 100.00 | 100.00 | 100.00 | 100.00 | 100.00 |
| LD7-1_2008_A_Mali | 0.00 | 0.00 | 100.00 | 0.00 | 100.00 | 100.00 | 100.00 | 100.00 | 100.00 | 100.00 |
| LD71a_2007_Astar_Cambodia | 100.00 | 0.00 | 100.00 | 8.06 | 100.00 | 11.36 | 0.00 | 71.60 | 0.00 | 98.61 |
| LE085_2008_A_Papua-New-Guinea | 0.00 | 0.00 | 100.00 | 0.00 | 100.00 | 100.00 | 100.00 | 100.00 | 100.00 | 100.00 |
| LE086_2008_A_Papua-New-Guinea | 0.00 | 0.00 | 100.00 | 0.00 | 100.00 | 100.00 | 100.00 | 100.00 | 100.00 | 100.00 |
| LE087_2008_A_Papua-New-Guinea | 0.00 | 0.00 | 100.00 | 0.00 | 100.00 | 100.00 | 100.00 | 100.00 | 100.00 | 100.00 |
| LE116-1_2008_A_Mali | 48.76 | 5.53 | 100.00 | 8.06 | 100.00 | 100.00 | 100.00 | 100.00 | 100.00 | 100.00 |
| LE20-1_2008_Astar_Ethiopia | 100.00 | 0.00 | 100.00 | 0.00 | 100.00 | 93.46 | 0.00 | 32.94 | 0.00 | 100.00 |
| LE3-1_2008_Astar_Ethiopia | 100.00 | 0.00 | 100.00 | 4.03 | 100.00 | 92.82 | 26.50 | 42.06 | 16.89 | 100.00 |
| LG099_2006_A_Bangladesh | 0.00 | 0.00 | 100.00 | 0.00 | 100.00 | 96.14 | 100.00 | 100.00 | 100.00 | 0.00 |
| LG102_2006_A_Bangladesh | 100.00 | 5.86 | 100.00 | 3.55 | 100.00 | 100.00 | 100.00 | 100.00 | 100.00 | 100.00 |
| LG103_2006_A_Bangladesh | 0.00 | 0.00 | 100.00 | 0.00 | 100.00 | 100.00 | 100.00 | 100.00 | 100.00 | 100.00 |
| LG104_2006_A_Bangladesh | 0.00 | 0.00 | 100.00 | 0.00 | 100.00 | 93.14 | 100.00 | 100.00 | 100.00 | 100.00 |
| LG105_2006_A_Bangladesh | 100.00 | 0.00 | 100.00 | 0.00 | 100.00 | 100.00 | 100.00 | 100.00 | 100.00 | 100.00 |
| LG107_2006_A_Bangladesh | 100.00 | 0.00 | 100.00 | 0.00 | 100.00 | 100.00 | 100.00 | 78.51 | 100.00 | 100.00 |
| LG108_2006_A_Bangladesh | 0.00 | 0.00 | 100.00 | 0.00 | 100.00 | 100.00 | 100.00 | 100.00 | 100.00 | 100.00 |
| LG110_2006_A_Bangladesh | 11.90 | 0.00 | 100.00 | 0.00 | 100.00 | 100.00 | 100.00 | 100.00 | 100.00 | 100.00 |
| LG112_2006_A_Bangladesh | 100.00 | 0.00 | 100.00 | 0.00 | 100.00 | 100.00 | 100.00 | 100.00 | 100.00 | 100.00 |
| LG113_2006_A_Bangladesh | 100.00 | 0.00 | 100.00 | 0.00 | 100.00 | 100.00 | 100.00 | 100.00 | 100.00 | 100.00 |
| LG115_2007_Aw_India | 100.00 | 100.00 | 100.00 | 14.77 | 100.00 | 24.97 | 100.00 | 100.00 | 0.00 | 26.11 |
| LG117_2009_A_Bangladesh | 0.00 | 4.43 | 100.00 | 4.03 | 100.00 | 100.00 | 100.00 | 100.00 | 0.00 | 0.00 |
| LG97_2006_A_Bangladesh | 100.00 | 0.00 | 100.00 | 0.00 | 100.00 | 17.68 | 100.00 | 100.00 | 100.00 | 100.00 |
| LG98_2006_A_Bangladesh | 100.00 | 0.00 | 100.00 | 3.79 | 100.00 | 100.00 | 100.00 | 78.51 | 100.00 | 100.00 |
| LH001-03_2010_A_Pakistan | 100.00 | 0.00 | 100.00 | 0.00 | 100.00 | 100.00 | 100.00 | 100.00 | 100.00 | 100.00 |
| LH221-6_2010_A_France-Reunion | 0.00 | 0.00 | 100.00 | 0.00 | 100.00 | 100.00 | 100.00 | 100.00 | 100.00 | 100.00 |
| LH37-1_2010_A_Senegal | 100.00 | 11.91 | 100.00 | 3.95 | 100.00 | 100.00 | 100.00 | 100.00 | 100.00 | 100.00 |
| LJ001-1_2012_A_Seychelles | 0.00 | 0.00 | 100.00 | 0.00 | 100.00 | 100.00 | 100.00 | 100.00 | 100.00 | 100.00 |
| LJ003-1_2012_A_Seychelles | 0.00 | 0.00 | 100.00 | 0.00 | 100.00 | 100.00 | 100.00 | 100.00 | 100.00 | 100.00 |
| LJ226-02_2012_A_France-Mayotte | 0.00 | 0.00 | 100.00 | 0.00 | 100.00 | 94.96 | 100.00 | 100.00 | 100.00 | 100.00 |
| LK135-01_2013_Astar_Comoros-Moheli | 100.00 | 0.00 | 100.00 | 0.00 | 100.00 | 11.68 | 0.00 | 71.60 | 0.00 | 98.61 |
| LK169-01_2013_A_France-Reunion | 0.00 | 0.00 | 100.00 | 0.00 | 100.00 | 100.00 | 100.00 | 100.00 | 100.00 | 100.00 |
| LK169-04_2013_A_France-Reunion | 22.73 | 0.00 | 100.00 | 0.00 | 100.00 | 100.00 | 100.00 | 100.00 | 100.00 | 100.00 |
| LL074-04_2014_A_France-Martinique | 0.00 | 0.00 | 100.00 | 0.00 | 100.00 | 100.00 | 100.00 | 100.00 | 100.00 | 100.00 |
| LL096-13_2014_A_France-Reunion | 0.00 | 0.00 | 100.00 | 0.00 | 100.00 | 100.00 | 100.00 | 100.00 | 100.00 | 100.00 |
| LL111-06_2014_A_France-Martinique | 0.00 | 0.00 | 100.00 | 0.00 | 100.00 | 100.00 | 100.00 | 100.00 | 100.00 | 100.00 |
| LL124-01_2014_A_France-Martinique | 0.00 | 0.00 | 100.00 | 0.00 | 100.00 | 100.00 | 100.00 | 100.00 | 100.00 | 100.00 |
| LL186-5_2014_A_France-Reunion | 0.00 | 0.00 | 100.00 | 0.00 | 100.00 | 100.00 | 100.00 | 100.00 | 100.00 | 100.00 |
| LM053-06_2015_A_Mauritius | 0.00 | 0.00 | 100.00 | 0.00 | 100.00 | 100.00 | 100.00 | 100.00 | 100.00 | 100.00 |
| LM054-06_2015_A_Mauritius | 0.00 | 0.00 | 100.00 | 0.00 | 100.00 | 100.00 | 100.00 | 100.00 | 100.00 | 100.00 |

| | | | | | | | | | | |
|---|---|---|---|---|---|---|---|---|---|---|
| LM054-17_2015_A_Mauritius | 0.00 | 0.00 | 100.00 | 0.00 | 100.00 | 100.00 | 100.00 | 100.00 | 100.00 | 100.00 |
| LM055-08_2015_A_Mauritius | 0.00 | 0.00 | 100.00 | 0.00 | 100.00 | 100.00 | 100.00 | 100.00 | 0.00 | 100.00 |
| LM069-01_2015_A_Mauritius | 0.00 | 0.00 | 100.00 | 0.00 | 100.00 | 100.00 | 100.00 | 100.00 | 100.00 | 100.00 |
| LM090-02_2015_A_France-Martinique | 0.00 | 0.00 | 100.00 | 0.00 | 100.00 | 100.00 | 100.00 | 100.00 | 100.00 | 100.00 |
| LM097-01_2015_A_Mauritius-Rodrigues | 0.00 | 0.00 | 100.00 | 0.00 | 100.00 | 100.00 | 100.00 | 100.00 | 100.00 | 100.00 |
| LM128_2007_A_Argentina | 0.00 | 0.00 | 100.00 | 0.00 | 100.00 | 100.00 | 100.00 | 100.00 | 100.00 | 100.00 |
| LM158_2013_A_Argentina | 0.00 | 0.00 | 100.00 | 0.00 | 100.00 | 100.00 | 100.00 | 100.00 | 100.00 | 100.00 |
| LM169_2005_A_Argentina | 0.00 | 0.00 | 100.00 | 0.00 | 100.00 | 100.00 | 100.00 | 100.00 | 100.00 | 100.00 |
| LM184_2013_A_Argentina | 0.00 | 0.00 | 100.00 | 0.00 | 100.00 | 100.00 | 100.00 | 100.00 | 100.00 | 100.00 |
| LM198_2015_A_Argentina | 0.00 | 0.00 | 100.00 | 0.00 | 100.00 | 100.00 | 100.00 | 100.00 | 100.00 | 100.00 |
| LM199_2015_A_Argentina | 0.00 | 0.00 | 100.00 | 0.00 | 100.00 | 100.00 | 100.00 | 100.00 | 100.00 | 100.00 |
| LM205_2010_A_Argentina | 0.00 | 0.00 | 100.00 | 0.00 | 100.00 | 100.00 | 100.00 | 100.00 | 100.00 | 100.00 |
| LM229_2012_A_Argentina | 0.00 | 0.00 | 100.00 | 0.00 | 100.00 | 100.00 | 100.00 | 100.00 | 100.00 | 100.00 |
| LM358-08_2015_A_France-Reunion | 0.00 | 0.00 | 100.00 | 0.00 | 100.00 | 100.00 | 100.00 | 100.00 | 100.00 | 100.00 |
| LM358-18_2015_A_France-Reunion | 0.00 | 0.00 | 100.00 | 0.00 | 100.00 | 100.00 | 100.00 | 100.00 | 100.00 | 100.00 |
| LMG-9322_1986_A_USA | 100.00 | 8.85 | 100.00 | 0.00 | 100.00 | 100.00 | 100.00 | 100.00 | 100.00 | 100.00 |
| LN006-18_2016_A_France-Reunion | 0.00 | 0.00 | 100.00 | 0.00 | 100.00 | 100.00 | 100.00 | 100.00 | 100.00 | 100.00 |
| LP027-03_2017_A_Seychelles | 0.00 | 0.00 | 100.00 | 0.00 | 100.00 | 100.00 | 100.00 | 100.00 | 100.00 | 100.00 |
| LP027-05_2017_A_Seychelles | 0.00 | 0.00 | 100.00 | 0.00 | 100.00 | 100.00 | 100.00 | 100.00 | 100.00 | 100.00 |
| LP027-13_2017_A_Seychelles | 0.00 | 0.00 | 100.00 | 0.00 | 100.00 | 100.00 | 100.00 | 100.00 | 100.00 | 100.00 |
| LP028-03_2017_A_Seychelles | 0.00 | 0.00 | 100.00 | 0.00 | 100.00 | 100.00 | 100.00 | 100.00 | 100.00 | 100.00 |
| LP028-05_2017_A_Seychelles | 0.00 | 0.00 | 100.00 | 0.00 | 100.00 | 100.00 | 100.00 | 100.00 | 100.00 | 100.00 |
| LP028-06_2017_A_Seychelles | 0.00 | 0.00 | 100.00 | 0.00 | 100.00 | 100.00 | 100.00 | 100.00 | 100.00 | 100.00 |
| LP029-13_2017_A_Seychelles | 0.00 | 0.00 | 100.00 | 0.00 | 100.00 | 100.00 | 100.00 | 100.00 | 100.00 | 100.00 |
| LP029-15_2017_A_Seychelles | 0.00 | 0.00 | 100.00 | 0.00 | 100.00 | 100.00 | 100.00 | 100.00 | 100.00 | 100.00 |
| LP030-1_2017_A_France-Martinique | 0.00 | 0.00 | 100.00 | 0.00 | 100.00 | 100.00 | 100.00 | 100.00 | 100.00 | 100.00 |
| LP031-1_2017_A_France-Martinique | 0.00 | 0.00 | 100.00 | 0.00 | 100.00 | 100.00 | 100.00 | 100.00 | 100.00 | 100.00 |
| LP032-1_2017_A_France-Martinique | 0.00 | 0.00 | 100.00 | 0.00 | 100.00 | 100.00 | 100.00 | 100.00 | 100.00 | 100.00 |
| LP033-1_2017_A_France-Martinique | 0.00 | 0.00 | 100.00 | 0.00 | 100.00 | 100.00 | 100.00 | 100.00 | 100.00 | 100.00 |
| LP034-1_2017_A_France-Martinique | 0.00 | 0.00 | 100.00 | 0.00 | 100.00 | 100.00 | 100.00 | 100.00 | 100.00 | 100.00 |
| LP035-1_2017_A_France-Martinique | 0.00 | 0.00 | 100.00 | 0.00 | 100.00 | 100.00 | 100.00 | 100.00 | 100.00 | 100.00 |
| LP036-1_2017_A_France-Martinique | 0.00 | 0.00 | 100.00 | 0.00 | 100.00 | 100.00 | 100.00 | 100.00 | 100.00 | 100.00 |
| LP162_2017_A_Pakistan | 0.00 | 0.00 | 100.00 | 0.00 | 100.00 | 100.00 | 100.00 | 100.00 | 100.00 | 100.00 |
| LP187_2017_A_Pakistan | 8.72 | 0.00 | 100.00 | 0.00 | 100.00 | 99.25 | 100.00 | 100.00 | 100.00 | 100.00 |
| LP188_2017_A_Pakistan | 0.00 | 0.00 | 100.00 | 0.00 | 100.00 | 98.82 | 100.00 | 100.00 | 100.00 | 100.00 |
| LP201_2017_A_Pakistan | 0.00 | 0.00 | 100.00 | 0.00 | 100.00 | 100.00 | 100.00 | 100.00 | 100.00 | 100.00 |
| LP215-01_2017_A_Pakistan | 0.00 | 0.00 | 100.00 | 0.00 | 100.00 | 100.00 | 100.00 | 100.00 | 100.00 | 100.00 |
| LP220-02_2017_A_Pakistan | 31.57 | 0.00 | 100.00 | 0.00 | 100.00 | 100.00 | 100.00 | 100.00 | 100.00 | 100.00 |
| LP229-03_2017_A_Pakistan | 0.00 | 0.00 | 100.00 | 0.00 | 100.00 | 100.00 | 100.00 | 100.00 | 100.00 | 100.00 |
| LP234-03_2017_A_Pakistan | 0.00 | 0.00 | 100.00 | 0.00 | 100.00 | 100.00 | 100.00 | 100.00 | 100.00 | 100.00 |
| LP235-06_2017_A_Pakistan | 30.62 | 0.00 | 100.00 | 0.00 | 100.00 | 100.00 | 100.00 | 100.00 | 100.00 | 100.00 |
| LP293-01_2017_A_Pakistan | 100.00 | 0.00 | 100.00 | 0.00 | 100.00 | 100.00 | 100.00 | 100.00 | 100.00 | 100.00 |

| | | | | | | | | | | |
|---|---|---|---|---|---|---|---|---|---|---|
| LP293-05_2017_A_Pakistan | 100.00 | 0.00 | 100.00 | 0.00 | 100.00 | 100.00 | 100.00 | 100.00 | 100.00 | 100.00 |
| LP295-05_2017_A_Pakistan | 0.00 | 0.00 | 100.00 | 0.00 | 100.00 | 98.93 | 100.00 | 100.00 | 100.00 | 100.00 |
| LP297-02_2017_A_Pakistan | 0.00 | 0.00 | 100.00 | 0.00 | 100.00 | 100.00 | 100.00 | 100.00 | 100.00 | 100.00 |
| NCPPB-1471_1963_A_China-Hong-Kong | 0.00 | 0.00 | 100.00 | 0.00 | 87.49 | 100.00 | 100.00 | 100.00 | 100.00 | 100.00 |
| NCPPB-211_1948_A_India | 5.65 | 0.00 | 100.00 | 0.00 | 100.00 | 100.00 | 100.00 | 100.00 | 100.00 | 100.00 |
| NCPPB-226_1949_A_New-Zealand | 0.00 | 0.00 | 100.00 | 0.00 | 100.00 | 100.00 | 100.00 | 100.00 | 100.00 | 100.00 |
| NCPPB-3562_1988_A_India | 100.00 | 0.00 | 100.00 | 7.94 | 100.00 | 100.00 | 100.00 | 100.00 | 100.00 | 100.00 |
| NCPPB-3607_1988_Astar_India | 100.00 | 0.00 | 100.00 | 4.03 | 100.00 | 92.82 | 10.30 | 100.00 | 0.00 | 98.61 |
| NCPPB-3608_1988_Aw_India | 100.00 | 100.00 | 100.00 | 0.00 | 100.00 | 11.15 | 100.00 | 100.00 | 100.00 | 100.00 |
| NCPPB-3610_1988_A_India | 7.66 | 0.00 | 100.00 | 0.00 | 100.00 | 100.00 | 100.00 | 100.00 | 100.00 | 100.00 |
| NCPPB-3612_1988_A_India | 100.00 | 0.00 | 100.00 | 0.00 | 100.00 | 100.00 | 100.00 | 100.00 | 100.00 | 100.00 |
| NCPPB-3615_1989_Astar_India | 100.00 | 0.00 | 100.00 | 0.00 | 100.00 | 93.25 | 0.00 | 100.00 | 0.00 | 100.00 |
| NCPPB-406_1957_A_New-Zealand | 0.00 | 0.00 | 100.00 | 0.00 | 100.00 | 100.00 | 100.00 | 100.00 | 100.00 | 100.00 |

### Supplementary References

Alegria MC, Docena C, Khater L, Ramos CH, da Silva AC & Farah CS (2004). New protein-protein interactions identified for the regulatory and structural components and substrates of the type III secretion system of the phytopathogen *Xanthomonas axonopodis* pathovar *citri*. **J. Bacteriol.** 186(18): 6186-6197. https://doi.org/10.1128/JB.186.18.6186-6197.2004

Astua-Monge G, Freitas-Astua J, Bacocina G, Roncoletta J, Carvalho SA & Machado MA (2005). Expression profiling of virulence and pathogenicity genes of *Xanthomonas axonopodis* pv. *citri*. **J. Bacteriol.** 187(3): 1201-1205. https://doi.org/10.1128/JB.187.3.1201-1205.2005

Bansal K, Midha S, Kumar S & Patil PB (2017). Ecological and evolutionary insights into *Xanthomonas* citri pathovar diversity. **Appl. Environ. Microbiol.** 83(9). https://doi.org/10.1128/AEM.02993-16

Facincani AP, Moreira LM, Soares MR, Ferreira CB, Ferreira RM, Ferro MI*, et al.* (2014). Comparative proteomic analysis reveals that T3SS, Tfp, and xanthan gum are key factors in initial stages of *Citrus sinensis* infection by *Xanthomonas citri* subsp. *citri*. **Funct. Integr. Genomics** 14(1): 205-217. https://doi.org/10.1007/s10142-013-0340-5

Ferreira CB, Moreira LM, Brigati JB, Lima LL, Ferro JA, Ferro MI*, et al.* (2017). Identification of new genes related to virulence of *Xanthomonas axonopodis* pv. *citri* during citrus host interactions. **Adv. Microbiol.** 7: 22-46. https://doi.org/10.4236/aim.2017.71003

Ferreira RM, Moreira LM, Ferro JA, Soares MRR, Laia ML, Varani AM*, et al.* (2016). Unravelling potential virulence factor candidates in *Xanthomonas citri* subsp. *citri* by secretome analysis. **PeerJ** 4: e1734. https://doi.org/10.7717/peerj.1734

Figueiredo JFL, Minsavage GV, Graham JH, White FF & Jones JB (2011). Mutational analysis of type III effector genes from *Xanthomonas citri* subsp. *citri*. **Eur. J. Plant Pathol.** 130(3): 339-347. https://doi.org/10.1007/s10658-011-9757-7

Gochez AM, Shantharaj D, Potnis N, Zhou X, Minsavage GV, White FF*, et al.* (2017). Molecular characterization of XopAG effector AvrGf2 from *Xanthomonas fuscans* ssp. *aurantifolii* in grapefruit. **Mol. Plant Pathol.** 18(3): 405-419. https://doi.org/10.1111/mpp.12408

Guo Y, Figueiredo F, Jones J & Wang N (2011). HrpG and HrpX play global roles in coordinating different virulence traits of *Xanthomonas axonopodis* pv. *citri*. **Mol. Plant Microbe Interact.** 24(6): 649-661. https://doi.org/10.1094/MPMI-09-10-0209

Huang CJ, Wu TL, Zheng PX, Ou JY, Ni HF & Lin YC (2021). Comparative genomic analysis uncovered evolution of pathogenicity factors, horizontal gene transfer events, and heavy metal resistance traits in citrus canker bacterium *Xanthomonas citri* subsp. *citri*. **Front. Microbiol.** 12: 731711. https://doi.org/10.3389/fmicb.2021.731711

Jalan N, Kumar D, Andrade MO, Yu F, Jones JB, Graham JH*, et al.* (2013). Comparative genomic and transcriptome analyses of pathotypes of *Xanthomonas citri* subsp. *citri* provide insights into mechanisms of bacterial virulence and host range. **BMC Genomics** 14: 551. https://doi.org/10.1186/1471-2164-14-551

Laia ML, Moreira LM, Dezajacomo J, Brigati JB, Ferreira CB, Ferro MI*, et al.* (2009). New genes of *Xanthomonas citri* subsp. *citri* involved in pathogenesis and adaptation revealed by a transposon-based mutant library. **BMC Microbiol.** 9: 12. https://doi.org/10.1186/1471-2180-9-12

Moreira LM, Almeida NF, Jr., Potnis N, Digiampietri LA, Adi SS, Bortolossi JC*, et al.* (2010). Novel insights into the genomic basis of citrus canker based on the genome sequences of two strains of *Xanthomonas fuscans* subsp. *aurantifolii*. **BMC Genomics** 11: 238. https://doi.org/10.1186/1471-2164-11-238

Patané JSL, Martins J, Jr., Rangel LT, Belasque J, Digiampietri LA, Facincani AP*, et al.* (2019). Origin and diversification of *Xanthomonas citri* subsp. *citri* pathotypes revealed by inclusive phylogenomic, dating, and biogeographic analyses. **BMC Genomics** 20(1): 700. https://doi.org/10.1186/s12864-019-6007-4

Rybak M, Minsavage GV, Stall RE & Jones JB (2009). Identification of *Xanthomonas citri* ssp. *citri* host specificity genes in a heterologous expression host. **Mol. Plant Pathol.** 10(2): 249-262. https://doi.org/10.1111/j.1364-3703.2008.00528.x

Yamazaki A, Hirata H & Tsuyumu S (2008). HrpG regulates type II secretory proteins in *Xanthomonas* axonopodis pv. *citri*. **J. Gen. Plant Pathol.** 74(2): 138-150. https://doi.org/10.1007/s10327-008-0075-7

Yan Q & Wang N (2012). High-throughput screening and analysis of genes of *Xanthomonas citri* subsp. *citri* involved in citrus canker symptom development. **Mol. Plant Microbe Interact.** 25(1): 69-84. https://doi.org/10.1094/MPMI-05-11-0121

Zhang Y, Jalan N, Zhou X, Goss E, Jones JB, Setubal JC*, et al.* (2015). Positive selection is the main driving force for evolution of citrus canker-causing *Xanthomonas*. **ISME J.** 9(10): 2128-2138. https://doi.org/10.1038/ismej.2015.15
